# Peripheral insulin resistance attenuates cerebral glucose metabolism and impairs working memory in healthy adults

**DOI:** 10.1101/2023.09.08.556948

**Authors:** Hamish A. Deery, Emma Liang, Robert Di Paolo, Katharina Voigt, Gerard Murray, M. Navyaan Siddiqui, Gary F. Egan, Chris Moran, Sharna D. Jamadar

## Abstract

People with insulin resistance are at increased risk for cognitive decline. Insulin resistance has previously been considered primarily a condition of ageing but it is increasingly seen in younger adults. Here, we explore the question that changes in insulin function in early adulthood have both proximal effects, and moderate or even accelerate changes in cerebral metabolism in ageing. Thirty-six younger (mean 27.8 years) and 43 older (mean 75.5) participants completed a battery of tests, including blood sampling, cognitive assessment and a simultaneous PET/MR scan. Cortical thickness and cerebral metabolic rate of glucose were derived for 100 regions and 17 functional networks. Older adults had lower rates of regional cerebral glucose metabolism than younger adults across the brain even after adjusting for lower cortical thickness in older adults. In younger adults, higher insulin resistance was associated with attenuated rates of regional cerebral glucose metabolism, but this was not seen in older adults. The largest effects of insulin resistance in younger adults were in prefrontal, parietal and temporal regions; and in the control, salience ventral attention, default and somatomotor networks. Higher rates of network glucose metabolism were associated with lower reaction time and psychomotor speed. Higher levels of insulin resistance were associated with lower working memory. Our results underscore the importance of insulin sensitivity to brain health and cognitive function across the adult lifespan, even in early adulthood.

**Significance Statement:** We show that preventing insulin resistance in early adulthood is important for ensuring efficient fuel supply for the brain and the maintenance of cognitive health across the adult lifespan. Glucose is the primary source of energy for the brain. Decreased glucose metabolism in the brain due to clinically significant levels of insulin resistance is associated with cognitive impairment. Although sub-clinical levels of insulin resistance have also been associated with brain changes, their impact on cerebral metabolism in healthy individuals is unclear. We showed for the first time that – while older adults have lower rates of cerebral metabolism - peripheral insulin resistance attenuates cerebral metabolism more so in healthy younger than healthy older adults, and impairs working memory.

## 1. Introduction

Insulin is central to the uptake of glucose into cells and the regulation of systemic blood glucose concentrations (1). Insulin enters the brain via insulin-independent mechanisms, where it used for a range of functions, such as cerebral glucose metabolism, the production of neurotransmitters, growth and regeneration of axons and neuron, circuit development, as well as the regulation of mood, behaviour and cognition (1, 2).

*Insulin resistance* describes how sensitive cells are to the effects of insulin. Greater insulin resistance (i.e., lower insulin sensitivity) results in greater amounts of insulin being required to transport glucose into cells. The underlying aetiology of insulin resistance has been well described (see (3) for a review). As the number of people with insulin resistance has increased (4, 5), so too has research on the pathophysiology of insulin in the brain. Much of this research has focussed on conditions in which insulin resistance is a hallmark characteristic, such as type 2 diabetes, Alzheimer’s disease and ageing (see (6-8) for a reviews). This research suggests that peripheral and central insulin resistance are linked (9, 10) and that insulin resistance is an important contributor to age-related diseases and cognitive decline (11-13).

Insulin resistance, even in those without type 2 diabetes, has been associated with grey matter atrophy and cognitive decline (14, 15). Longitudinal studies have linked peripheral insulin resistance in otherwise metabolically healthy people with subsequent declines in cognitive performance (16, 17). Variations in glycaemia in the absence of a clinical diagnoses of prediabetes or insulin resistance can cause grey matter atrophy (18, 19), reduced white matter integrity and impaired cognition in people in their late 20s, 30s and 40s (18-20).

Although the causes of cognitive decline in insulin resistance are multifaceted, altered cerebral glucose metabolism is considered a major factor (2, 8), with changes in insulin resistance in midlife or earlier becoming increasingly of interest (21). Although the impact of insulin resistance on whole-brain glucose metabolism is equivocal in healthy older adults, (6, 22-24), insulin resistance has been associated with lower glucose uptake regionally, including in the thalamus and caudate (23), regions of the prefrontal and temporal cortices, cingulate and insula (22) and the medial orbital frontal cortex (23). Age-related reductions in cerebral metabolism occur independently of cortical atrophy (25), indicating that metabolic reductions are not simply a result of loss of tissue or cell bodies in ageing but also occur from a loss of metabolic efficiency or function.

To date, most research exploring the links between insulin resistance and brain glucose metabolism has been with people in mid-to-late life. With greater recognition of the contribution of early adult factors to later life brain health, it is important to better understand whether there are associations between insulin resistance, cerebral glucose metabolism and cognition in younger people with few cardiometabolic risk factors. Understanding these associations can help guide the need for further research or health interventions and is especially important as rates of insulin resistance are growing in younger people (26).

Here we investigated the associations between age, insulin resistance, cerebral glucose metabolism, and cognition. We hypothesised that: 1) older people would have greater insulin resistance and lower cortical thickness than younger people; 2) older people would have lower regional cerebral metabolic rates of glucose than younger people, even after adjusting for lower cortical thickness in older people; 3) greater insulin resistance would be associated with lower cerebral metabolic rate of glucose and that this association would be moderated by age, with the effect being stronger in older adults; and 4) greater cerebral metabolic rate of glucose and lower insulin resistance would be associated with better cognitive test performance.

## 2 Results

### 2.1 Sample characteristics

The characteristics of the whole sample, as well as the younger and older participants, are shown in Table 1. The mean age of the whole sample was 53.8 years (SD=24.6) years. The proportion of women was 52%. The average years of education was 17.5; BMI was 25.0, resting heart rate was 78 BMP, systolic and diastolic blood pressure was 136 and 82 mmHg; and cortical thickness was 2.43 mm. Mean fasting blood glucose was 4.99 mmol/L, insulin 4.36 mIU/L, HOMA-IR 1.00 and HOMA-IR2 0.57.

**Table 1.** Demographics for the whole sample and comparison of older and younger groups. Continuous variables are mean (standard deviation); categorical variables are number and %.

|  | Whole Sample |  | Younger |  | Older |  | Younger vs Older p-value<sup>1</sup> |
| --- | --- | --- | --- | --- | --- | --- | --- |
|  | Mean | SD | Mean | SD | Mean | SD |  |
| Age (years) | 53.8 | 24.6 | 27.9 | 6.3 | 75.5 | 5.8 | <.001 |
| Sex (number and % female) | 41 (52%) |  | 21 (58%) |  | 20 (47%) |  | .295 |
| Education (years) | 17.5 | 3.3 | 18.1 | 2.9 | 16.9 | 3.6 | .167 |
| Body Mass Index (kg/m<sup>2</sup>) | 25.0 | 4.1 | 24.2 | 4.7 | 25.6 | 3.5 | .133 |
| Resting Heart Rate (bpm) | 77.9 | 15.7 | 82.7 | 17.4 | 73.9 | 13.0 | .013 |
| Systolic Blood Pressure (mmHg) | 135.5 | 26.0 | 119.6 | 17.7 | 148.8 | 24.4 | <.001 |
| Diastolic Blood Pressure (mmHg) | 81.7 | 12.7 | 79.4 | 13.4 | 83.6 | 11.9 | .144 |
| Cortical Thickness (mm) | 2.43 | 0.12 | 2.52 | 0.07 | 2.36 | 0.11 | <.001 |
| Fasting blood glucose (mmol/L) | 4.99 | 0.54 | 4.77 | 0.40 | 5.17 | 0.58 | <.001 |
| Fasting insulin (mIU/L) | 4.36 | 2.93 | 4.76 | 3.12 | 4.03 | 2.74 | .268 |
| HOMA-IR | 1.00 | 0.73 | 1.02 | 0.71 | 0.97 | 0.75 | .783 |
| HOMA-IR2 | 0.57 | 0.39 | 0.61 | 0.40 | 0.53 | 0.38 | .410 |
| HVLT: Delayed recall (total) | 8.29 | 2.66 | 9.39 | 2.49 | 7.37 | 2.47 | <.001 |
| HVLT: Recognition discrimination index | 9.83 | 2.00 | 10.63 | 1.70 | 9.19 | 2.02 | <.001 |
| Digit Span: Forward (longest) | 6.75 | 1.30 | 6.64 | 1.53 | 6.84 | 1.07 | .502 |
| Digit Span: Backwards (longest) | 5.28 | 1.28 | 5.47 | 1.34 | 5.12 | 1.22 | .221 |
| Category Switch: RT in switch trials (sec) | 1.80 | 0.64 | 1.42 | 0.40 | 2.12 | 0.63 | <.001 |
| Category Switch: SSRT | 394.6 | 354.2 | 284.1 | 244.8 | 487.1 | 404.7 | .010 |
| Digit Substitution: Correct count | 44.5 | 22.0 | 64.0 | 12.2 | 28.6 | 13.8 | <.001 |
| Digit Substitution: Sec per correct count | 4.37 | 6.95 | 3.55 | 9.68 | 5.08 | 2.95 | .017 |
| Stop Signal: Mean stop signal delays (sec) | 0.33 | 0.16 | 0.28 | 0.15 | 0.38 | 0.16 | .009 |
| Stop Signal: Stop signal reaction time (sec) | 0.56 | 0.12 | 0.52 | 0.11 | 0.59 | 0.12 | .004 |
| WASI FSIQ2 T-score | 116.8 | 15.3 | 109.0 | 12.1 | 123.2 | 14.8 | <.001 |

<sup>1</sup>P-values are based on T-test for continuous and Ch-square for categorical variables.

The mean age of the younger group was 27.8 years (SD=6.2) and the older group 75.5 years (SD=5.8). The proportion of women was higher in the younger group (58%) than the older group (47%) but this difference was not statistically significant (p=0.295). The average years of education was similar in the younger group (18.1) and the older group (17.1). Mean fasting blood glucose was greater in the older (5.2 mmol/L) than the younger (4.77 mmol/L) group (p<.001). Fasting insulin concentration were not significantly different between the two groups.

The older group had higher mean blood pressure than the younger group but this was only statistically significant for systolic blood pressure (149mmHg vs 120mmHg, p<.001). Although participants did not have a known history of hypertension upon study entry, a total of 29 people in the older group and nine in the younger group met guideline criteria for the diagnosis of hypertension from the measurements taken. Older people had a greater mean BMI than younger people (25.6 vs 24.2 kg/m^2^) but this difference was not statistically significant (p=.133). General liner models of age and insulin resistance differences in CMR_GLC,_ with blood pressure and the other demographics as covariates, are reported in the Supplement (Table S6).

#### 2.2.1 Age and insulin resistance and cortical thickness

Measures of insulin resistance (HOMA-IR and HOMA-IR2) were greater in the younger group than the older group but these differences were not statistically significant (Table 1). Four people in the younger age group and five in the older age group had HOMA-IR levels greater than 1.8. One person in the younger group and four people in the older group had HOMA-IR levels greater than 2.7. These thresholds of 1.8 and 2.7 have previously been correlated with clinical levels of insulin resistance and other cardiometabolic risk factors (27).

The older group had lower whole brain cortical thickness than the younger group (p<.001). Older people also had lower cortical thickness than those in the younger group in 94 of 100 regions.

#### 2.2.2 Age, cerebral cortical thickness and metabolic rate of glucose

Across age groups, greater cortical thickness was associated with greater regional CMR_GLC_ (Figure 1A, and Table S1) in 83 of the 100 regions with the largest effect sizes in the superior, middle and medial frontoparietal cortex. Taking the association between age and cortical thickness into account, those in the older group had lower CMR_GLC_ than those in the younger group in all regions (Figure 1B and Table S2). The effect sizes ranged from .20 to .24 in regions in the somatomotor, salience ventral attention, control and default networks, to less than .10 mostly in regions in the visual network and sub-cortical structures. The largest effects were in the medial, ventral and dorsal prefrontal cortices in the default network; the lateral prefrontal cortex, insula and the parietal operculum and lobule in the salience ventral attention network; the lateral prefrontal and temporal cortices in the control network; and regions of the somatomotor network.

**Figure 1.**
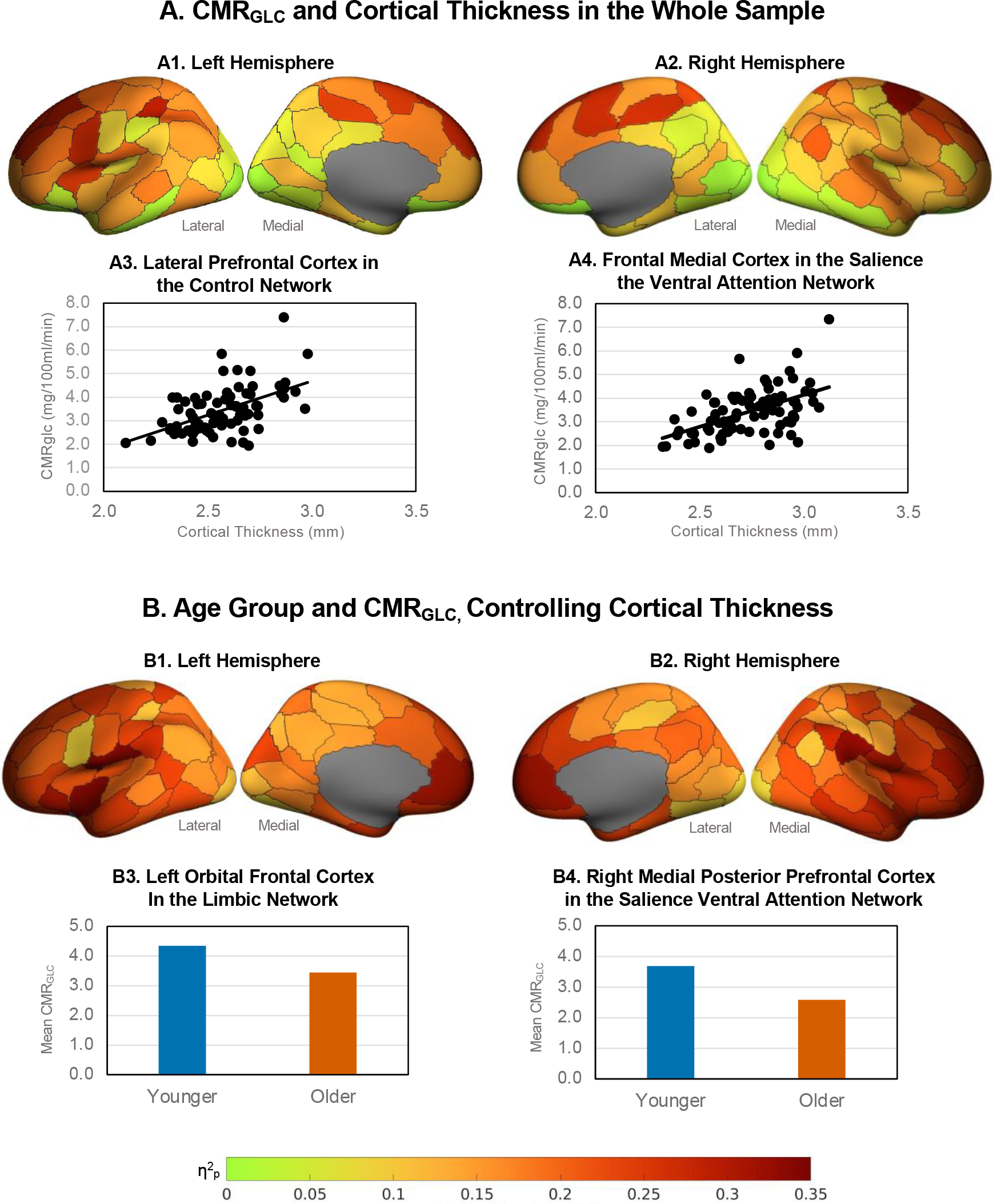
Association between age, cortical thickness and CMR_GLC_. A. Effect sizes (η^2^_p_) of higher regional CMR_GLC_ associations with higher cortical thickness in the whole sample in left (A1) and right (A2) hemispheres (data is from Table S1). Example associations in (A3) the lateral prefrontal cortex in the control network, and (A4) the frontal medial cortex in the salience ventral attention network. B. Association between age group and CMR_GLC_ controlling for cortical thickness in the left (B1) and right (B2) hemispheres (data is from Table S2. Example age group and CMR_GLC_ relationships in (B3) the left orbital frontal cortex in the limbic network, and (B4) the right medial posterior prefrontal cortex in the salience ventral attention network. Figures were produced using the toolbox at: https://github.com/StuartJO/plotSurfaceROIBoundary

### 2.3 Age, Insulin resistance and cerebral metabolic rate of glucose

Greater HOMA-IR was associated with lower CMR_GLC_ across all regions (Figure 2A and Table S3). The effect sizes ranged from .14 to .16 in regions in the salience ventral attention, somatomotor, default and control networks, to less than .08 in regions in the visual and control networks and the sub-cortical structures. The largest effects of HOMA-IR were in the lateral prefrontal, medial parietal and parietal operculum in the salience ventral attention network; the ventral and dorsal prefrontal cortices, parietal lobule and posterior cingulate in the default network; the lateral prefrontal cortex, precuneus and cingulate in the control network; and several somatomotor regions.

**Figure 2.**
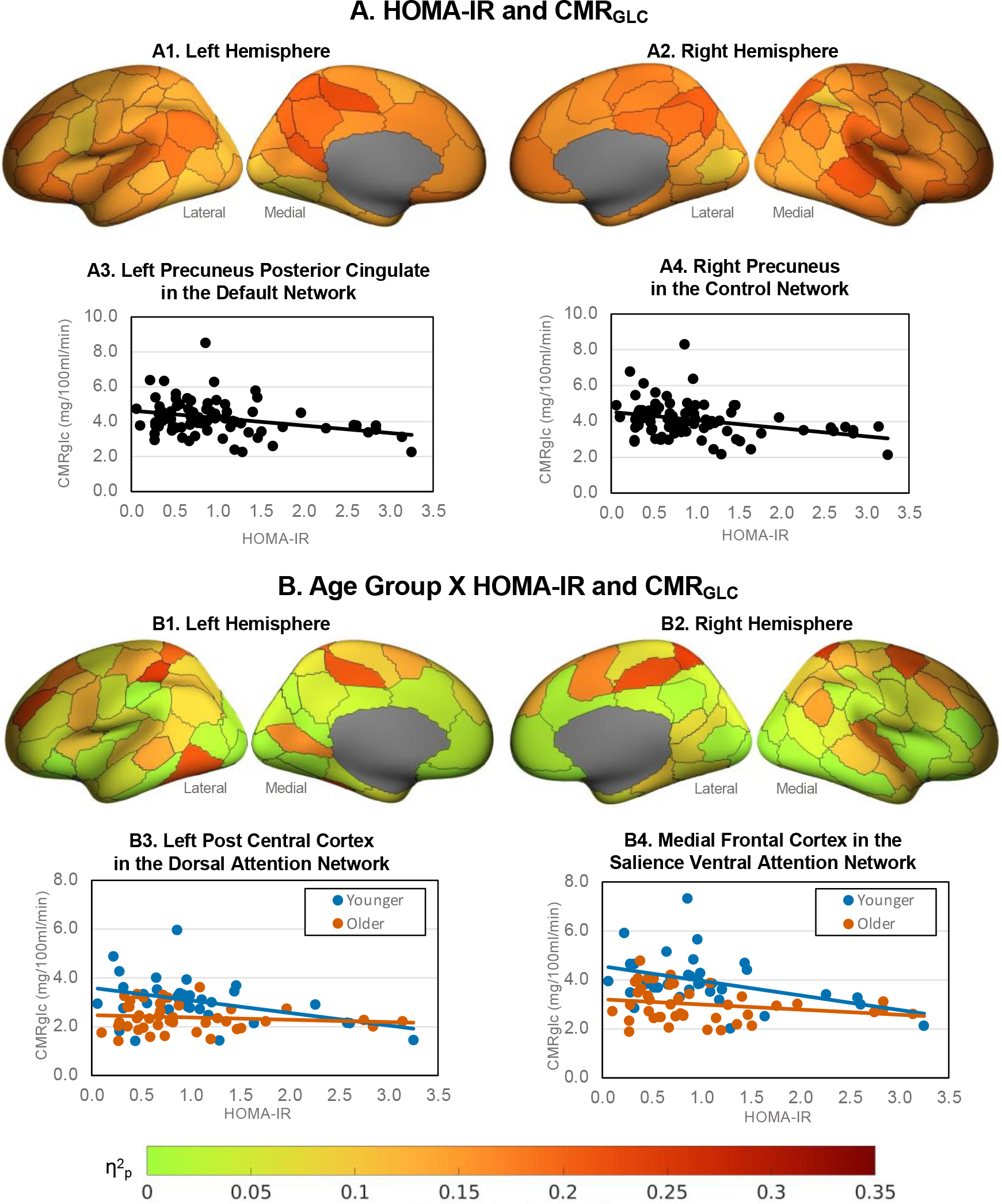
Effect sizes (η^2^_p_) of regional CMR_GLC_ associations with (A) HOMA-IR and (B) age group x HOMA-IR, controlling for cortical thickness (data in Table S2). Example HOMA-IR effects in (A3) the left precuneus posterior cingulate in the default network and (A4) right precuneus In the control network . Example age group X HOMA-IR interactions in (B3) left post central cortex in the dorsal attention network and (B4) right medial frontal cortex in the salience ventral attention network.

We found a statistically significant interaction between the older and younger groups and HOMA-IR levels on CMR_GLC_ in 41 regions (Figure 2B; Table S2). In post-hoc tests we found that greater insulin resistance was associated with lower CMR_GLC_ in the younger group but not the older group (Table S3). The nature of this interaction was such that a 10% increase in HOMA-IR in the younger group was associated with a lower CMR_GLC_ across regions of between 3.3% to 7.3%. In the older group a 10% increase in HOMA-IR was not statistically associated with a change in CMR_GLC_ (zero to -2.2%). The largest reductions for younger adults were in the dorsal prefrontal cortex and parietal medial regions of the default network; the lateral prefrontal cortices in the control and salience ventral attention networks; the medial frontal cortex in the default network; the superior parietal and post central regions of the dorsal attention network; the temporal parietal cortex and insula; and regions of the somatomotor network.

We also found statistically significant age group x HOMA-IR interactions in all Schaefer networks (Table S4). The nature of these interactions was similar to those at the regional level in that greater HOMA-IR was associated with lower network CMR_GLC_ in the younger but not the older group. Large effect sizes (η^2^_p_ >.20) of HOMA-IR were found for younger adults in the control, salience ventral attention, somatomotor, default and visual networks.

### 2.5 Cerebral Metabolic Rates of Glucose and Cognition

Five principal components (PCs) of cognition with eigenvalues greater than one were identified across the six cognitive tests and eleven measures, explaining 81% of the variance. The varimax rotation converged in five iterations (see Table S5). Cognitive control (task-switching and WASI FSIQ) loaded most strongly on the first principal component. Visuospatial processing (digit substitution) loaded most strongly on the second principal component; response inhibition (stop-signal) on the third component; working memory (digit span) on the fourth component; and verbal learning and memory (HVLT) on and the fifth component.

Higher CMR_GLC_ in the limbic, default and subcortical networks was associated with better performance on cognitive control (principal component 1, Table S6). In particular, higher network CMR_GLC_ was associated with a faster reaction time in task-switching (see Figure 3A). Higher CMR_GLC_ in the control and default networks was associated with faster visuospatial processing speed (principal component 2, digit substitution; Figure 3B). Higher CMR_GLC_ in the dorsal attention and control networks was associated with better response inhibition (principal component 3, stop signal; Figure 3C). Interestingly, lower CMR_GLC_ was associated with higher WASI FSIQ2 and number of correct responses in the digit substitution task.

**Figure 3.**
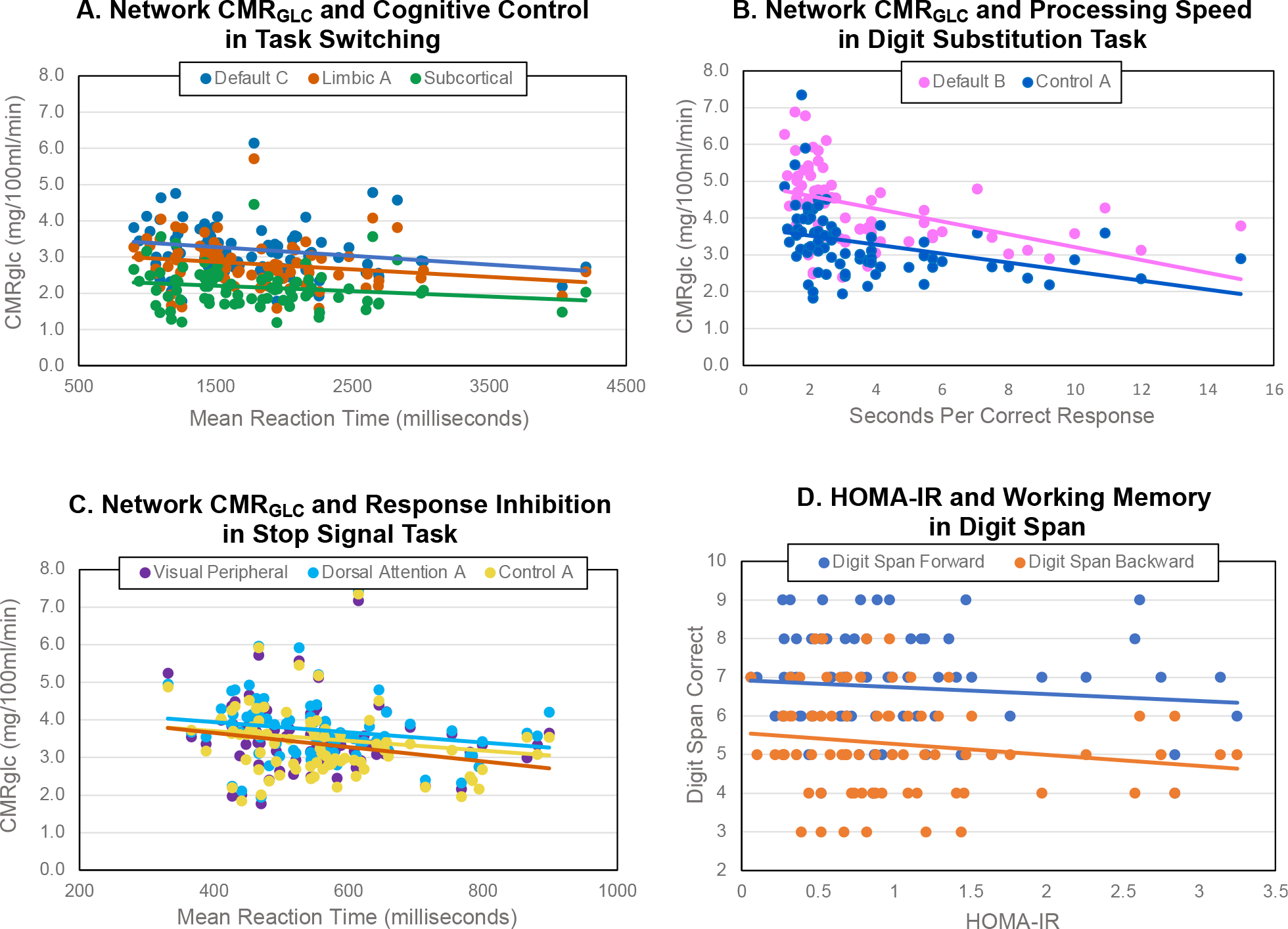
Relationship between network CMR_GLC_ and cognitive measures to illustrate significant effects from the general linear models (Tables S6). Significant negative associations of higher network CMR_GLC_ and: (A) shorter mean reaction time in task switching trials (PC1), (B) lower processing speed in the digit substitution task (loading on PC2) and (c) shorter mean reaction time in stop signal trials measuring response inhibition (PC3). (D) Significant negative association of higher HOMA-IR and lower longest forward and backward digit span performance.

Older adults had slower reaction time than younger adults in task switching but higher WASI FSIQ (first principal component; Table S6). Older adults also had slower visuospatial processing speed in the digit substitution task (second principal component). Higher HOMA-IR had a direct effect on worse working memory (principal component 4, digit span). However, none of the interaction terms of HOMA-IR and network CMR_GLC_ were significant in the stepwise regression models, indicating that levels of HOMA-IR did not moderate the relationship between CMR_GLC_ and cognition.

## 3. Discussion

### 3.1 The Association Between CMR_GLC_ and Age, HOMA-IR and Cortical Thickness

As expected, we found that older adults had lower cortical thickness than younger adults. However, unexpectedly, older adults in our sample were not more insulin resistant than younger adults. We also found that greater age and greater insulin resistance were associated with lower cerebral glucose metabolism across the brain, particularly in prefrontal and temporal cortices. Although we found that greater age was associated with lower cortical thickness and that lower cortical thickness was associated with lower cerebral glucose metabolism, the association between older age and lower cerebral glucose metabolism remained after adjusting for differences in cortical thickness. These results are consistent with previous primary research and a recently published systematic review and meta-analysis. (25). They suggest that older adults have less metabolic active cortical tissue than younger adults but they also have less efficient metabolism of glucose on a per tissue basis. Our results are also consistent with research in older adults in which higher HOMA-IR was associated with lower glucose metabolism in regions of the medial orbital, prefrontal and temporal cortices and the cingulate and insula (22, 28).

Consistent with our hypothesis, we found that the association between insulin resistance and cerebral glucose metabolism varied by age group in 41 regions, particularly in regions in the prefrontal, parietal, temporal and somatomotor cortices. However, in an unexpected finding, the association between greater insulin resistance and lower cerebral glucose metabolism was seen in the younger group but not the older group. Rates of cerebral metabolism in insulin resistant younger adults were also lower in all networks than in their more insulin sensitive counterparts, particularly in the control, salience ventral attention, somatomotor and default mode networks. Previous research has also shown a negative effect from metabolic dysfunction on grey matter volume, white matter integrity and cognition in people in their late 20s, 30s and 40s (18-20). Our results add to this by suggesting that cerebral metabolism is also attenuated by insulin resistance in otherwise healthy younger adults.

It is striking that higher insulin resistance was associated with greater CMR_GLC_ reductions in younger than older adults. This is somewhat counterintuitive, as we had expected that insulin resistance would be related to CMR_GLC_ reductions more so in older adults. Our results are also different to the one other study by Nugent et al. (28) that also included younger adults, in which HOMA-IR did not correlate with CMR_GLC_ in any brain region. One possible reason for these different results is that our younger sample had a broader age range by 12 years and slightly higher HOMA-IR2 than Nugent et al. (20-42 years vs 18-30 years; HOMA-IR2 0.6 vs 0.5). We also note that our older adult group had slightly lower mean and standard deviation fasting insulin and HOMA-IR levels than our younger adult group (see Table S1). This is in contrast to the association between ageing and increasing insulin resistance shown in studies of the general population (29, 30). As such, our older cohort seems to be more insulin sensitive compared to the general older adult population. We do not rule out the possibility of an association between greater insulin resistance and lower cerebral glucose metabolism in other adult samples. Such patterns have been previously reported in other studies and we add to this work by suggesting that greater insulin resistance and lower cerebral glucose metabolism is seen in younger samples than previously examined. Replicating and extending our findings with other younger adult populations and with a wider range of insulin resistance levels in older adults is warranted.

### 3.2 CMR_GLC_ Primarily in “Higher Order” Brain Networks is Associated with Cognition

Consistent with our hypothesis, higher regional CMR_GLC_ in the default, control, dorsal attention, limbic and subcortical networks was associated with faster psychomotor speed and reaction time in the digit substitution, task switching and stop signal tasks (see Figure 3). However, these associations were not moderated by levels of peripheral insulin resistance. The networks in which higher CMR_GLC_ was associated with faster processing speed and reaction time reflect mostly “higher order” (e.g., control and attention) as opposed to primary sensory networks (e.g., visual). These results suggest that higher rates of glucose metabolism in the “higher order” networks supports flexible, adaptive responses in goal-directed behaviour, as well as inhibition of inappropriate actions. These results also likely reflect the fact that the cognitive battery primarily indexed memory, spatiotemporal processing and attentional control.

Unexpectedly, higher CMR_GLC_ was associated with lower WASI scores. WASI, which is an estimate of full scale IQ (FSIQ), was acquired as a more robust estimate of cognitive reserve, which is often measured using proxy indices such as reading tests or educational attainment (31). Our sample had a relatively high mean WASI FSIQ2 of 116 and seven participants scored above 140, all in the older age group, indicating high levels of cognitive reserve. Two younger participants scored above 130. Together these results suggests that reduced rates of cerebral glucose in the brain of adults primarily attenuate processing speed rather than task accuracy, even in older adults who retain relatively high cognitive reserve.

We did not find significant network CMR_GLC_ associations with episodic and working memory (PC4 and PC5). However, we did find higher HOMA-IR to be associated with worse working memory, an effect also found in previous research (32, 33), suggesting that insulin resistance is a risk factor for working memory impairment. Unexpectedly, we did not find age group differences for episodic and working memory. A large body of research has shown that older adults typically show a decline in episodic and working memory compared with younger adults (see (34) (35)), further highlighting that our sample of older adults were particularly cognitively-healthy with high cognitive reserve. Additional research is warranted to investigate how closely cerebral glucose metabolism and insulin are coupled with cognition across the adult lifespan.

### 3.3 Possible Mechanisms and Future Directions

The mechanisms through which insulin resistance leads to changes in brain function remain to be fully understood. However, a growing body of research suggests that the brain is sensitive to the levels of peripheral insulin and uses insulin in a range of functions (36-38). Research also suggesting shared pathways or mechanisms driving changes to the brain in age and insulin resistance (8). The mechanism include metabolic disturbances from an increase in neuronal insulin resistance, decreased brain insulin receptor number and function, impaired insulin signalling, a pro-inflammatory state and mitochondrial dysfunction (39, 40). Prolonged peripheral hyperinsulinemia can also decrease insulin receptors at the blood-brain barrier, thereby reducing insulin transport into the brain (39). Neurons in the hypothalamus and brainstem that are responsible for energy homoestatis and feeding are also impaired in insulin resistance (2). Much of the research in this area has been with participants with a diagnosis of insulin resistance or Type 2 diabetes (7) or been limited to older participants (6, 22, 24). However, our results indicate that variability in peripheral insulin resistance within a healthy range is also associated with changes in cerebral metabolism, particularly in younger adults.

The current study used a cross-sectional design, limiting conclusions about any causal relationships. The differences we found between groups may also reflect underlying cohort differences rather than age-related changes. As noted above, our older adult population appears to be particularly healthy and relatively homogenous, at least in terms of insulin resistance and cognitive reserve. Longitudinal research could test whether changes in insulin function in early adulthood not only have a proximal effect, such as those reported here, but also moderate or even accelerate cerebral metabolic changes in ageing.

Our study is also limited by the absence of middle aged adults. Research on structural and functional brain networks have reported quadratic trajectories of age differences, with an inflection point somewhere in the third to fifth decade of life (see (41) for review). However, the lack of middle aged adults precludes the identification and quantification of ageing trajectories across the full adult lifespan. Additional research is needed to elucidate these patterns.

HOMA-IR is considered a reliable clinical and research tool for the assessment of levels of insulin resistance. Nevertheless, a limitation of HOMA-IR is that it represents a single snapshot of the complex glucose-insulin system (42). Research using dynamic measures (e.g., hyperinsulinemic-euglycemic clamp or 2-hour glucose tolerance tests) could improve our understanding of the complexity of insulin signalling and metabolism in the periphery and the brain in ageing and across the spectrum of health and metabolic-related diseases.

### 3.4 Implications for the Maintenance of Brain Health Across the Adult Lifespan

The results of the current study suggest that insulin resistance may contribute to brain health even in younger adults. Diet, lifestyle (e.g., sleep, exercise, stress), and genetics are risk factors for an increase in insulin resistance and would appear to be targets for public health interventions and clinical application to optimise both peripheral and central glucose metabolism (2, 43). Pharmaceutical treatments for diabetes that target peripheral glucose and bodyweight reductions may reduce the risk for cognitive decline (44, 45). Medications that increase sensitivity to insulin have been used for years in people with diabetes and are now being considered in people without diabetes to improve brain health. Our work suggests that future research should consider including people in early adulthood given the signals between insulin resistance and cerebral glucose metabolism we report here.

## 4. Method

Full details of the methods are provided in the Supplement. This study design, hypotheses and analyses were preregistered at OSF registrations (https://osf.io/93mnd).

### 4.1 Participants

Participants were recruited from the general community via local advertising. The final sample included 79 individuals, 36 younger (mean 27.8; SD 6.2;) and 43 older (mean 75.5; SD 5.8; range 66-86 years) adults (see Table S1). Exclusion criteria included a known diagnosis or history of hypertension, diabetes, or both, reported by participants at the time of recruitment to the study.

### 4.2 Data Acquisitions

Participants completed an online demographic and lifestyle questionnaire and a cognitive test battery. Briefly, the following cognitive measure were used (see Supplement for details): a combined and age-normalised index of verbal comprehension and perceptual reasoning from the Wechsler Abbreviated Scale of Intelligence; delayed recall and a recognition discrimination index from the Hopkins Verbal Learning Test; length of longest correct series of forward and backward recall from a digit span test to index working memory; switching cost in a task switching test to index cognitive flexibility; reaction time in a stop signal task to measure response inhibition; and number and seconds per correct response in a digit substitution task to measure visuospatial performance and processing speed.

Participants underwent a 90-minute simultaneous MR-PET scan in a Siemens (Erlangen) Biograph 3-Tesla molecular MR scanner (for scan parameters see Supplementary Information). At the beginning of the scan, half of the 260 MBq FDG tracer was administered via the left forearm as a bolus, the remaining 130 MBq of the FDG tracer dose was infused at a rate of 36ml/hour over 50 minutes (46). Non-functional MRI scans were acquired during the first 12 minutes, including a T1 3DMPRAGE and T2 FLAIR. Thirteen minutes into the scan, list-mode PET and T2* EPI BOLD-EPI sequences were initiated. A 40-minute resting-state scan was undertaken in naturalistic viewing conditions. Plasma radioactivity levels were measured every 10 minutes throughout the scan.

### 4.3 MRI Pre-Processing, Cortical Thickness and Grey matter Volume

For the T1 images, the brain was extracted in Freesurfer; quality of the pial/white matter surface was manually checked, corrected and registered to MNI152 space using Advanced Normalization Tools (ANTs). Cortical thickness for the Schaefer 100 regions was obtained from the Freesurfer reconstruction statistics for each participant.

### 4.4 PET Image Reconstruction and Pre-Processing

The list-mode PET data for each subject were binned into 344 3D sinogram frames of 16s intervals. Attenuation was corrected via the pseudo-CT method for hybrid PET-MR scanners (47). Ordinary Poisson-Ordered Subset Expectation Maximization algorithm (3 iterations, 21 subsets) with point spread function correction was used to reconstruct 3D volumes from the sinogram frames. The reconstructed DICOM slices were converted to NIFTI format with size 344 × 344 × 127 (d size: 1.39 × 1.39 × 2.03 mm^3^) for each volume. All 3D volumes were temporally concatenated to form a single 4D NIFTI volume. After concatenation, the PET volumes were motion corrected using FSL MCFLIRT (48), with the mean PET image used to mask the 4D data. PET images were corrected for partial volume effects using the modified Müller-Gartner method (49) implemented in Petsurf.

### 4.5 Cerebral Metabolic Rates of Glucose

Calculations of regional CMR_GLC_ were undertaken in PMOD 4.4 (http://www.pmod.com) using the FDG time activity curves for the Schaefer 100 atlas parcellation and AAL subcortical structures. The FDG in the plasma samples was decay-corrected for the time between sampling and counting, and used as the input function to Patlak models. A lumped constant of 0.89 was used (50), and equilibrium (t) set at 10 mins, the time corresponding to the peak of the bolus and onset of a stable signal (46). The fractional blood space (vB) was set at 0.05 (51). Participant’s plasma glucose (mmol) was entered in the model from their baseline blood sample.

CMR_GLC_ in the 17 networks was calculated from the regional CMR_GLC_ values for each participant. Because the regions within a network differ in cortical volume, the regional CMR_GLC_ values could not simply be averaged. Rather, each regional CMR_GLC_ value was weighted by the percentage its volume represented within the total network cortical volume. An overall subcortical CMR_GLC_ value was calculated by weighting each structure by the percentage its volume represented from the total volume of the subcortical structures.

### 4.6 HOMA-IR

A blood sample taken prior to FD infusion was used to collect 2ml of plasma for insulin and glucose measurement, which was undertaken by a commercial laboratory. HOMA-IR was calculated as fasting glucose (mmol/L) x fasting insulin (µU/ml) / 22.5 (52). The constant of 22.5 is a normalising factor for normal fasting plasma insulin and glucose (i.e., 4.5 mmol/L x 5 μU/ml = 22.5). Higher HOMA-IR values indicate greater insulin resistance. We also calculated HOMA-IR2 (https://www.rdm.ox.ac.uk/). We compared the relationship between HOMA-IR and HOMA-IR2 with CMR_GLC_ and found minimal to no differences (see Table S2 and S7). This was expected as HOMA-IR2 models increases in the insulin secretion curve for plasma glucose concentrations above 10 mmol/L (53); a threshold that than none of our participants reached. Hence, the results reported here are based on HOMA-IR.

### 4.7 Data Analysis

The CMR_GLC_ and HOMA-IR data was inspected for and found to be satisfactory for assumptions of normality and potential impact of any outliers (see Supplement Figure S2).

#### 4.7.1 Age, Cortical Thickness, HOMA-IR and CMR_GLC_

*Hypothesis 1:* Independent sample T-tests were run to test hypothesis 1 that older people would have greater insulin resistance and lower cortical thickness than younger people. *Hypothesis 2 and 3:* A series of general linear models (GLMs) was run in which regional CMR_GLC_ was the dependent variable and age group, HOMA-IR and age group x HOMA-IR were the predictors. Cortical thickness in the same region was included as a covariate. For the subcortical structures, whole brain average cortical thickness was used as the covariate. The age group main effect was used to assess hypothesis 2 that older people would have lower regional cerebral metabolic rates of glucose than younger people, even after adjusting for lower cortical thickness in older people. The age group x HOMA-IR effects were used to test hypothesis 3 that greater insulin resistance would be associated with lower cerebral metabolic rate of glucose and that this association would be moderated by age, with the effect being stronger in older adults. For significant age group x HOMA-IR effects, post-hoc GLMs were run separately for younger and older adults.

A series of GLMs were also run for CMR_GLC_ at the 17 network level with the same design. Partial eta squared (η^2^_p_) was used to quantify the effect sizes in the GLMs. Each series of analyses was also FDR-corrected at p < .05 for the overall GLM. We also calculated the percentage change in regional CMR_GCL_ from a 10% change in HOMA-IR from the slope of the regression lines for younger and older adults separately.

#### 4.7.2 CMR_GLC_ and Cognition

*Hypothesis 4:* We applied data reduction techniques to reduce the dimensions in the cognitive test data. The cognitive scores (see Supplement) were converted to Z-scores and entered in a Principal Component Analysis (PCA). Principal components (PCs) with eigenvalues greater than one were retained and subject to varimax rotation to optimally reduce dimensionality (54). Participant component scores were saved for further analyses.

A series of GLMs was run to test hypothesis 4 that greater cerebral metabolic rate of glucose and lower insulin resistance would be associated with better cognitive test performance. The five cognition PCs were entered as the dependent variable, with CMR_GLC_ in 17 networks entered as independent variables, together with age group, whole brain cortical thickness and HOMA-IR. To test for a moderating effect of HOMA-IR, where a network CMR_GLC_ predicted a principal component of cognition, a product term was created between CMR_GLC_ and HOMA-IR. Stepwise regression was used with the significant CMR_GLC_ network(s) entered in block 1, and the product term(s) with HOMA-IR in block 2. A significant increase in variance explained from block 1 to block 2 was indicative of moderation.

## Funding Information

Jamadar is supported by an Australian National Health and Medical Research Council (NHMRC) Fellowship (APP1174164).

## Notes

**Competing Interest Statement:** The authors declare no conflicts of interest.

### Competing Interest Statement

The authors have declared no competing interest.

### Summary of Updates

Based on reviewer feedback update the introduction and discussion, as well as additional analyses in supplement using other demographics and alternative measure of insulin resistance (HOMA-IR2).

