## Supplementary material for "Peripheral insulin resistance attenuates cerebral glucose metabolism and impairs working memory in healthy adults": PNAS_Supplementary Materials_Deery et al.pdf

**~ Supplementary Information ~**

**Supplementary Methods**

***1. Ethical Considerations***

The study protocol was reviewed and approved by the Monash University Human Research Ethics Committee in accordance with Australian Code for the Responsible Conduct of Research (2007) and the Australian National Statement on Ethical Conduct in Human Research (2007). Administration of ionizing radiation was approved by the Monash Health Principal Medical Physicist, following the Australian Radiation Protection and Nuclear Safety Agency Code of Practice (2005). For participants older than 18 years, the annual radiation exposure limit of 5 mSv applies. The effective dose in this study was 4.9 mSv.

***2. Recruitment***

Ninety participants were recruited from the general community via local advertising. An initial screening interview ensured that participants had the capacity to provide informed consent, did not have a history of hypertension, a diagnosis of diabetes, neurological or psychiatric illness, and were not taking psychoactive medication that could affect cognitive function or metabolism. Participants were also screened for claustrophobia, non-MR compatible implants, and clinical or research PET scan in the past 12 months. Women were screened for current or suspected pregnancy. Participants received a $100 voucher for participating in the study.

Eleven participants were excluded from further analyses due to blood haemolysis or well counter issues preventing insulin measurement or kinetic modelling (n=7), excessive head motion (n=2) or incomplete PET scan or image reconstruction (n=2).

***3. Cognitive Battery***

Prior to the scan, participants completed an online demographic and lifestyle questionnaire including age, sex, education, height and weight, history of smoking, alcohol and recreational drug use. Participants also completed a cognitive test battery consisting of measures of general intelligence, working memory, cognitive flexibility, inhibitory control and verbal learning.

**Wechsler Abbreviated Scale of Intelligence (WASI-IQ).** An assessment of intelligence suitable for ages 6-90 years (1). There are 4 subtests: block design, vocabulary, matrix reasoning and similarities. WASI-IQ was scored by converting raw scores into a scale score, which were transformed into a composite score reflecting verbal comprehension and perceptual reasoning abilities (FSIQ2). This score was converted to an age-based T scores established in a normal population.

**Hopkins Verbal Learning Test (HVLT).** A three-trial list learning and free recall task comprising 12 words, four words from each of three semantic categories (2). Approximately 20–25 minutes later, a delayed recall trial and a recognition trial was completed. The delayed recall required free recall of any words remembered. The recognition trial comprised 24 words, including the 12 target words and 12 false-positives, six semantically related, and six semantically unrelated. Delayed recall (total words recalled) and a recognition discrimination index (number of correct minus number of false positives in the recognition task) were calculated.

**Digit Span.** A measure of verbal short term and working memory used in two formats: Forward and backward digit span (3). Participants were presented with a series of digits, and are asked to repeat them in either the order presented (forward span) or in reverse order (backwards span). After two consecutive failures of the same length, the test was stopped. Scores were derived as the length of longest correct series for both forward and backward recall.

**Task Switching.** A computer-based test in which participants were given a word and had to perform one of two simple categorisation tasks, depending on the cue that appeared with the word: 1) 'living' task. If the cue was a heart, participants were asked to categorise the word via a key press based on whether it represents a LIVING versus a NON-LIVING object; and 2) 'size' task. If the cue was an arrow-cross, participants were asked to categorise the word via a key press based on whether it represents an object that is BIGGER or SMALLER than a basketball. The cue selection for each new trial was randomised. Half the test trials were switch trials; half non-switch trials. Half the switch and non-switch trials were congruent in the key presses for either task, half were incongruent. The measures used included the mean latency of correctly responding to a switch trial and switch cost. Switch cost is the difference between mean correct latency of switch trials and nonswitch trials with positive value indicating participants were slower on switch trials, that is, there was a latency cost to switching (4).

**Stop Signal.** A computer-based test in which participants were presented an arrow that pointed either right or left (5). The task was to press the left response key if the arrow pointed to the left and press the right response key if the arrow pointed to the right, unless a signal beep is played after the presentation of the arrow. In this case the response should be stopped before execution. The delay between presentation of arrow and signal beep (starting at 250ms) was adjusted up or down (by 50ms) depending on performance. The delay got longer if the previous signal stop was successful (up to 1150ms) and smaller if the previous signal stop was not successful (down to 50ms). The stimulus onset asynchrony between the start of each trial (onset of fixation circles) was 2000ms. Variables were the mean reaction time in stop signal trials and stop signal reaction time. Stop signal reaction time is an estimate of inhibition ability, that is, the time required to stop the initiated go-process. The slower the stop signal reaction time, the more difficult to stop the go-process.

**Digit Symbol Substitution.** A computer-based task in which participant were presented with an 18 columns x 16 rows matrix (6). The task was to translate symbols shown above the matrix (key) into digits in the matrix within a two minute period. Total count of correct responses and seconds per correct response were recorded.

#### *4. MR-PET Data Acquisition*

Participants underwent a 90-minute simultaneous MR-PET scan in a Siemens (Erlangen) Biograph 3-Tesla molecular MR scanner. Participants were directed to consume a high-protein/low-sugar diet for the 24 hours prior to the scan. They were also instructed to fast for six hours and to drink 2–6 glasses of water. Prior to FDG infusion, participants were cannulated in the vein in each forearm and a 10ml baseline blood sample taken. At the beginning of the scan, half of the 260 MBq FDG tracer was administered via the left forearm as a bolus, providing a strong PET signal from the beginning of the scan. The remaining 130 MBq of the FDG tracer dose was infused at a rate of 36ml/hour over 50 minutes, minimising the amount of signal decay over the course of the data acquisition. We have previously demonstrated that this protocol provides a good balance between a fast increase in signal-to-noise ratio at the start of the scan, and maintenance of signal-to-noise ratio over the duration of the scan (7).

Participants were positioned supine in the scanner bore with their head in a 32-channel radiofrequency head coil and were instructed to lie as still as possible. The scan sequence was as follows. Non-functional MRI scans were acquired during the first 12 minutes, including a T1 3DMPRAGE (TA = 3.49 min, TR = 1640ms, TE = 234ms, flip angle = 8°, field of view = 256 × 256 mm2, voxel size = 1.0 × 1.0 × 1.0 mm3, 176 slices, sagittal acquisition) and T2 FLAIR (TA = 5.52 min, TR = 5,000ms, TE = 396ms, field of view = 250 × 250 mm2, voxel size = .5 × .5 × 1 mm3, 160 slices) to image the anatomical grey and white matter structures, respectively. Thirteen minutes into the scan, list-mode PET (voxel size = 1.39 x 1.39 x 5.0mm3) and T2* EPI BOLD-fMRI (TA = 40 minutes; TR = 1000ms, TE = 39ms, FOV = 210 mm2, 2.4 × 2.4 × 2.4 mm3 voxels, 64 slices, ascending axial acquisition) sequences were initiated. A 40-minute resting-state scan was undertaken in naturalistic viewing conditions watching a movie of a drone flying over the Hawaii Islands. At 53 minutes, pseudo-continuous arterial spin labelling (pc-ASL) began, and at 58 minutes, diffusion-weighted imaging (DWI) was acquired with 71 directions to index white matter connectivity. pcASL, DWI and fMRI results are not reported here.

Plasma radioactivity levels were measured throughout the duration of the scan. Beginning at 10-minutes post infusion onset, 5ml blood samples were taken from the right forearm using a vacutainer at 10-minute intervals for a total of nine samples. The blood sample were immediately placed in a Heraeus Megafuge 16 centrifuge (ThermoFisher Scientific, Osterode, Germany) and spun at 2,000 rpm (RCF ~ 515g) for 5 minutes. 1,000-µL plasma was pipetted, transferred to a counting tube, and placed in a well counter for four minutes. The count start time, total number of counts, and counts per minute were recorded for each sample.

#### *5. Correction for Partial Volume Effects*

PET images were corrected for partial volume effects using the modified Müller-Gartner method implemented in PetSurf (https://surfer.nmr.mgh.harvard.edu/fswiki/PetSurfer) (8, 9). The method corrects for white matter spill in and grey matter spill out of the PET signal. The equation subtracts from the grey matter voxel signal the white matter signal (convoluted by the point spread function) and divides by the grey matter signal. This division can introduce over-correction at the grey matter boundary, and hence a grey matter binary mask is recommended, with the threshold level needing to be chosen. A grey matter threshold of 20-30% is recommended in ageing because atrophy can influence results (8). For our analyses, we chose a 25% grey matter threshold and surface-based spatial smoothing (9). We used a Gaussian kernel with a full width at half maximum of 12 mm to increase the signal-to-noise ratio. Subcortical structures were partial volume corrected and spatially smoothed in volume space and merged with the cortical data.

**Supplementary Results**

***1. Normality and Outlier Considerations***

A.

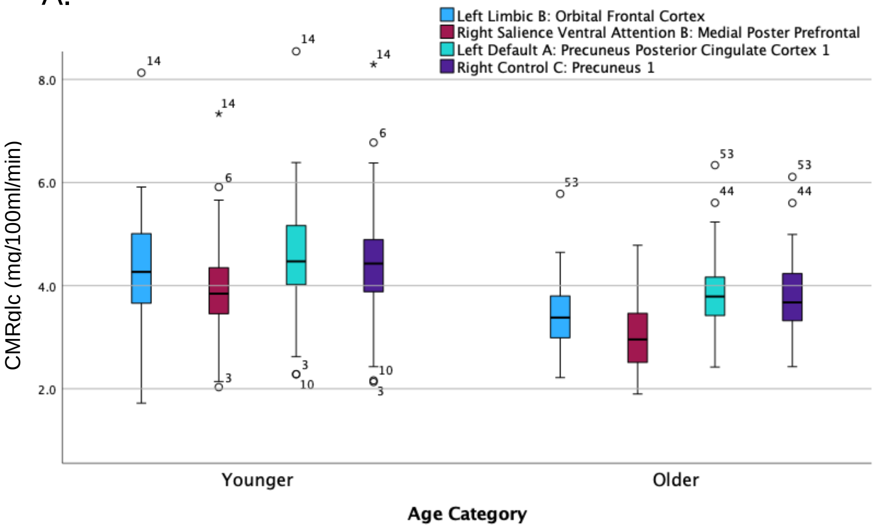

B.

| Partial Correlation: Age Group and CRMglc, Controlling for Cortical Thickness | | | | | | |
|---|---|---|---|---|---|---|
| | All Participants | | 2 Outliers Excluded | | Log Transformed | |
| | r | p | r | p | r | p |
| Left LimbiC: B: Orbital Frontal Cortex 1 | -0.43 | 0.001 | -0.46 | <.001 | -0.40 | <.001 |
| Right Salience Ventral Attention B: Medial Posterior Prefrontal 1 | -0.44 | 0.001 | -0.47 | <.001 | -0.45 | <.001 |
| Partial Correlation: HOMA-IR and CRMglc, Controlling for Age Group | | | | | | |
| | All Participants | | 2 Outliers Excluded | | Log Transformed | |
| | r | p | r | p | r | p |
| Left Default A: Precuneus Posterior Cingulate Cortex 1 | -0.34 | 0.032 | -0.36 | 0.011 | -0.37 | <.001 |
| Right Control C: Precuneus 1 | -0.36 | 0.050 | -0.38 | 0.011 | -0.38 | <.001 |

Figure S1. A. Boxplot of younger and older adult for regions shown in the Figures in main manuscript. We found the data to have slight non-normality based on the Kolmogorov-Smirnov Test. Cases were also identified where regional $CMR_{GLC}$ values were high or low for some participants relative the to sample distribution, although they are within a physiological plausible range (A). Analyses were run log transforming and excluding outliers in regions and compared to the results for all participants without transformation (B). Results are similar in all cases, suggesting that the analyses are robust to the outliers and any non-normality. Hence, all participants and raw $CRM_{GLC}$ data was used for the analyses presented in the manuscript.

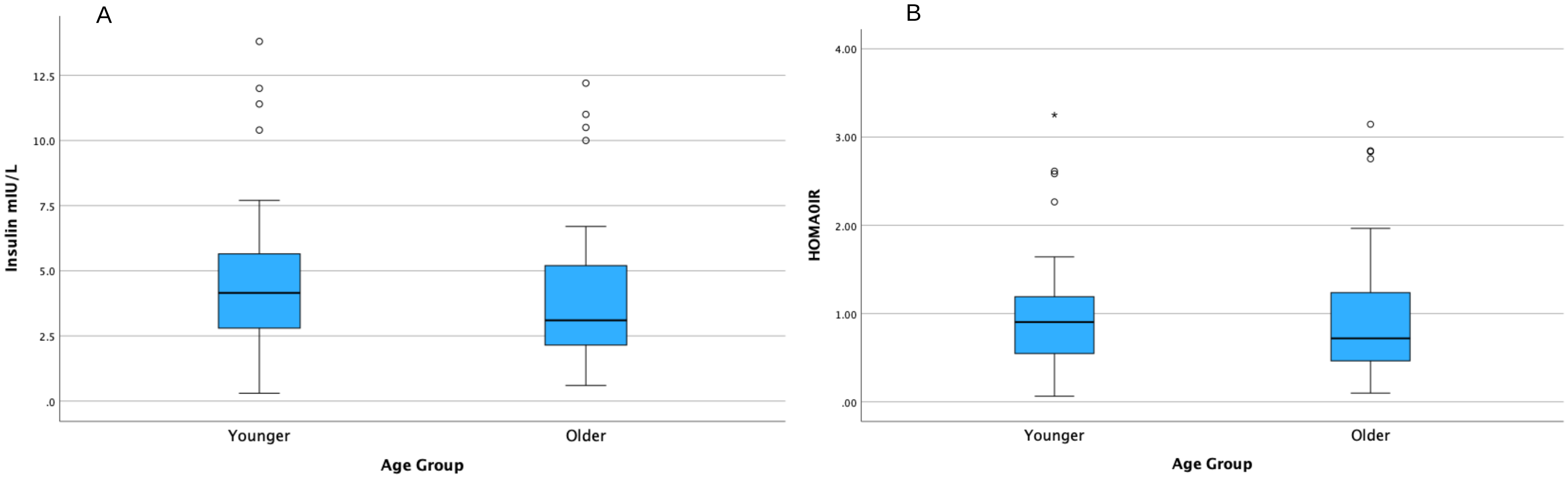

Figure S2. Boxplot of younger and older adult: (A) insulin, and (B) HOMA-IR

### *2. Cortical Thickness, Age, HOMA-IR and $CMR_{GLC}$*

Tables S2 to S4 present the results from the main analyses in the manuscript.

Table S1. General linear models of regional $CMR_{GLC}$ and cortical thickness in the whole sample. The $\eta^2_p$ values are plotted on the brain surface in Figure 1 of the main manuscript.

| Left Hemisphere | | | | Right Hemisphere | | | |
|---|---|---|---|---|---|---|---|
| | F | p-FDR | $\eta^2_p$ | | F | p-FDR | $\eta^2_p$ |
| Visual Central: Extra Striate Cortex 1 | 2.5 | 0.133 | 0.029 | Visual Central: Extra Striate Cortex 1 | 3.2 | 0.088 | 0.038 |
| Visual Central: Extra Striate Cortex 2 | 0.5 | 0.515 | 0.006 | Visual Central: Extra Striate Cortex 2 | 0.1 | 0.789 | 0.001 |
| Visual Central: Striate Cortex 1 | 0.5 | 0.500 | 0.006 | Visual Central: Extra Striate Cortex 3 | 4.1 | 0.054 | 0.048 |
| Visual Central: Extra Striate Cortex 3 | 4.7 | 0.039 | 0.055 | Visual Peripheral: Striate Cortex Calcarine 1 | 0.4 | 0.540 | 0.005 |
| Visual Peripheral: Extra Striate Inferior 1 | 3.3 | 0.083 | 0.039 | Visual Peripheral: Extra Striate Inferior 1 | 11.7 | 0.002 | 0.126 |
| Visual Peripheral: Striate Cortex Calcarine 1 | 10.8 | 0.003 | 0.117 | Visual Peripheral: Extra Striate Superior 1 | 6.2 | 0.019 | 0.071 |
| Visual Peripheral: Extra Striate Cortex Sup 1 | 5.6 | 0.026 | 0.065 | | | | |
| Somatomotor A: 1 | 15.9 | 0.000 | 0.164 | Somatomotor A: 1 | 7.0 | 0.013 | 0.080 |
| Somatomotor A: 2 | 15.8 | 0.000 | 0.163 | Somatomotor A: 2 | 7.2 | 0.013 | 0.081 |
| Somatomotor B: Auditory 1 | 15.4 | 0.001 | 0.160 | Somatomotor A: 3 | 14.8 | 0.001 | 0.154 |
| Somatomotor B: S2 1 | 6.8 | 0.015 | 0.077 | Somatomotor A: 4 | 13.7 | 0.001 | 0.145 |
| Somatomotor B: S2 2 | 19.0 | 0.000 | 0.190 | Somatomotor B: Auditory 1 | 18.6 | 0.000 | 0.187 |
| Somatomotor B: Central 1 | 7.5 | 0.011 | 0.084 | Somatomotor B: S2 1 | 6.6 | 0.016 | 0.075 |
| | | | | Somatomotor B: S2 2 | 14.8 | 0.001 | 0.154 |
| | | | | Somatomotor B: Central 1 | 8.3 | 0.008 | 0.093 |
| Dorsal Attention A: Temporal Occipital 1 | 14.9 | 0.001 | 0.156 | Dorsal Attention A: Temporal Occipital 1 | 2.4 | 0.138 | 0.029 |
| Dorsal Attention A: Parietal Occipital 1 | 7.1 | 0.013 | 0.081 | Dorsal Attention A: Parietal Occipital 1 | 8.6 | 0.007 | 0.096 |
| Dorsal Attention A: Superior Parietal Lobule 1 | 14.6 | 0.001 | 0.153 | Dorsal Attention A: Superior Parietal Lobule 1 | 9.6 | 0.005 | 0.106 |
| Dorsal Attention B: Post Central 1 | 4.6 | 0.042 | 0.054 | Dorsal Attention B: Post Central 1 | 15.2 | 0.001 | 0.158 |
| Dorsal Attention B: Post Central 2 | 33.4 | 0.000 | 0.292 | Dorsal Attention B: Post Central 2 | 24.2 | 0.000 | 0.230 |
| Dorsal Attention B: Post Central 3 | 16.4 | 0.000 | 0.168 | Dorsal Attention B: Frontal Eye Fields 1 | 39.1 | 0.000 | 0.325 |
| Dorsal Attention B: Frontal Eye Fields 1 | 19.8 | 0.000 | 0.196 | | | | |
| Salience Ventral Attention A: Parietal Operculum 1 | 8.8 | 0.006 | 0.098 | Salience Ventral Attention A: Parietal Operculum 1 | 10.2 | 0.003 | 0.112 |
| Salience Ventral Attention A: Insula: 1 | 11.7 | 0.002 | 0.126 | Salience Ventral Attention A: Insula: 1 | 10.3 | 0.003 | 0.113 |
| Salience Ventral Attention A: Insula: 2 | 26.7 | 0.000 | 0.248 | Salience Ventral Attention A: Parietal Medial 1 | 28.6 | 0.000 | 0.261 |
| Salience Ventral Attention A: Parietal Medial 1 | 25.6 | 0.000 | 0.240 | Salience Ventral Attention A: Frontal Medial 1 | 29.3 | 0.000 | 0.265 |
| Salience Ventral Attention A: Frontal Medial 1 | 26.9 | 0.000 | 0.250 | Salience Ventral Attention B: Inferior Parietal Lobule 1 | 10.2 | 0.003 | 0.112 |
| Salience Ventral Attention B: Lateral Prefrontal Cortex 1 | 32.1 | 0.000 | 0.284 | Salience Ventral Attention B: Lateral Prefrontal Cortex 1 | 19.7 | 0.000 | 0.196 |
| Salience Ventral Attention B: Medial Posterior Prefrontal 1 | 16.6 | 0.000 | 0.170 | Salience Ventral Attention B: Medial Posterior Prefrontal 1 | 16.7 | 0.000 | 0.171 |
| Limbic A: Temporal Pole 1 | 5.5 | 0.026 | 0.064 | Limbic A: Temporal Pole 1 | 7.0 | 0.013 | 0.080 |
| Limbic A: Temporal Pole 2 | 1.4 | 0.250 | 0.017 | Limbic B: Orbital Frontal Cortex 1 | 0.0 | 0.900 | 0.000 |
| Limbic B: Orbital Frontal Cortex 1 | 0.7 | 0.427 | 0.009 | | | | |
| Control A: Intraparietal Sulcus 1 | 20.4 | 0.000 | 0.201 | Control A: Intraparietal Sulcus 1 | 15.5 | 0.001 | 0.161 |
| Control A: Lateral Prefrontal Cortex 1 | 18.4 | 0.000 | 0.185 | Control A: Lateral Prefrontal Cortex 1 | 6.3 | 0.019 | 0.072 |
| Control A: Lateral Prefrontal Cortex 2 | 30.4 | 0.000 | 0.273 | Control A: Lateral Prefrontal Cortex 2 | 13.9 | 0.001 | 0.147 |
| Control B: Lateral Prefrontal Cortex 1 | 14.3 | 0.001 | 0.150 | Control B: Temporal 1 | 2.4 | 0.137 | 0.028 |
| Control C: Precuneus 1 | 5.2 | 0.031 | 0.060 | Control B: inferior parietal lobule 1 | 15.1 | 0.001 | 0.157 |
| Control C: Precuneus 2 | 9.1 | 0.005 | 0.101 | Control B: Lateral Prefrontal Cortexd 1 | 24.8 | 0.000 | 0.234 |
| Control C: Cingulate Posterior 1 | 6.2 | 0.019 | 0.072 | Control B: Lateral Prefrontal Cortexv 1 | 13.1 | 0.001 | 0.139 |
| | | | | Control C: Cingulate Posterior 1 | 7.9 | 0.009 | 0.089 |
| | | | | Control C: Precuneus 1 | 5.6 | 0.026 | 0.064 |
| Default A: Dorsal Prefrontal Cortex 1 | 33.3 | 0.000 | 0.291 | Default A: Inferior Parietal Lobule 1 | 21.3 | 0.000 | 0.208 |
| Default A: Precuneus Posterior Cingulate Cortex1 | 8.4 | 0.007 | 0.094 | Default A: Dorsal Prefrontal Cortex 1 | 23.4 | 0.000 | 0.224 |
| Default A: Medial Prefrontal Cortex 1 | 11.6 | 0.002 | 0.125 | Default A: Precuneus Posterior Cingulate Cortex 1 | 4.1 | 0.053 | 0.049 |
| Default B: Temp 1 | 13.0 | 0.001 | 0.138 | Default A: Medial Prefrontal Cortex 1 | 13.3 | 0.001 | 0.141 |
| Default B: Temp 2 | 17.2 | 0.000 | 0.175 | Default B: Dorsal Prefrontal Cortex 1 | 30.2 | 0.000 | 0.272 |
| Default B: Inferior Parietal Lobule 1 | 14.6 | 0.001 | 0.153 | Default B: Ventral Prefrontal Cortex 1 | 9.5 | 0.005 | 0.105 |
| Default B: Dorsal Prefrontal Cortex 1 | 31.1 | 0.000 | 0.277 | Default B: Ventral Prefrontal Cortex 2 | 16.1 | 0.000 | 0.166 |
| Default B: Lateral Prefrontal Cortex 1 | 20.6 | 0.000 | 0.202 | Default C: Retro Superior Parietal Lobuleenial 1 | 7.5 | 0.011 | 0.085 |
| Default B: Ventral Prefrontal Cortex 1 | 2.8 | 0.106 | 0.034 | Default C: Parahippocampal Cortex 1 | 8.7 | 0.007 | 0.097 |
| Default B: Ventral Prefrontal Cortex 2 | 14.8 | 0.001 | 0.154 | | | | |
| Default C: Retro Superior Parietal Lobuleenial 1 | 2.9 | 0.102 | 0.035 | | | | |
| Default C: Parahippocampal Cortex 1 | 9.4 | 0.005 | 0.104 | | | | |
| Temporal Parietal 1 | 12.3 | 0.002 | 0.132 | Temporal Parietal 1 | 15.5 | 0.001 | 0.160 |
| | | | | Temporal Parietal 2 | 16.5 | 0.000 | 0.169 |
| | | | | Temporal Parietal 3 | 10.6 | 0.003 | 0.116 |

Table S2. General linear models of age group, HOMA-IR and age group x HOMA-IR effect on regional $CMR_{GLC}$, including cortical thickness as a covariate. The $\eta^2_p$ values are plotted on the brain surface in Figure 2 of the main manuscript.

**Left Hemisphere**

| | Overall Model | | | Cortical Thickness | | | Age Group | | | HOMA-IR | | | Age Group x HOMA-IR | | |
|---|---|---|---|---|---|---|---|---|---|---|---|---|---|---|---|
| | F | p-FDR | $\eta^2_p$ | F | p | $\eta^2_p$ | F | p | $\eta^2_p$ | F | p | $\eta^2_p$ | F | p | $\eta^2_p$ |
| Visual Central: Extra Striate Cortex 1 | 2.5 | 0.053 | 0.117 | 2.8 | 0.101 | 0.036 | 4.6 | 0.036 | 0.058 | 4.1 | 0.048 | 0.052 | 1.1 | 0.298 | 0.015 |
| Visual Central: Extra Striate Cortex 2 | 3.1 | 0.022 | 0.142 | 4.3 | 0.042 | 0.055 | 6.0 | 0.017 | 0.074 | 7.5 | 0.008 | 0.092 | 0.2 | 0.635 | 0.003 |
| Visual Central: Striate Cortex 1 | 3.0 | 0.023 | 0.141 | 3.0 | 0.089 | 0.039 | 6.8 | 0.011 | 0.084 | 5.9 | 0.018 | 0.073 | 0.3 | 0.603 | 0.004 |
| Visual Central: Extra Striate Cortex 3 | 4.9 | 0.001 | 0.211 | 7.0 | 0.010 | 0.086 | 10.8 | 0.002 | 0.127 | 6.4 | 0.014 | 0.079 | 1.2 | 0.274 | 0.016 |
| Visual Peripheral: Extra Striate Inferior 1 | 4.5 | 0.003 | 0.196 | 3.8 | 0.055 | 0.049 | 10.0 | 0.002 | 0.119 | 4.8 | 0.032 | 0.060 | 0.5 | 0.491 | 0.006 |
| Visual Peripheral: Striate Cortex Calcarine 1 | 7.3 | 0.000 | 0.284 | 6.5 | 0.013 | 0.080 | 9.9 | 0.002 | 0.118 | 8.9 | 0.004 | 0.107 | 9.4 | 0.003 | 0.113 |
| Visual Peripheral: Extra Striate CortexSup 1 | 7.2 | 0.000 | 0.281 | 7.2 | 0.009 | 0.088 | 15.3 | 0.000 | 0.172 | 8.9 | 0.004 | 0.108 | 2.0 | 0.162 | 0.026 |
| Somatomotor A: 1 | 10.0 | 0.000 | 0.351 | 5.0 | 0.029 | 0.063 | 13.9 | 0.000 | 0.158 | 7.8 | 0.007 | 0.095 | 7.3 | 0.009 | 0.090 |
| Somatomotor A: 2 | 8.7 | 0.000 | 0.319 | 3.9 | 0.051 | 0.050 | 8.1 | 0.006 | 0.099 | 9.3 | 0.003 | 0.112 | 5.1 | 0.027 | 0.065 |
| Somatomotor B: Auditory 1 | 9.6 | 0.000 | 0.342 | 7.1 | 0.010 | 0.087 | 11.7 | 0.001 | 0.136 | 11.7 | 0.001 | 0.136 | 5.4 | 0.022 | 0.069 |
| Somatomotor B: S2 1 | 7.2 | 0.000 | 0.280 | 4.8 | 0.032 | 0.061 | 13.7 | 0.000 | 0.157 | 8.4 | 0.005 | 0.102 | 1.4 | 0.244 | 0.018 |
| Somatomotor B: S2 2 | 13.6 | 0.000 | 0.424 | 7.4 | 0.008 | 0.090 | 23.4 | 0.000 | 0.240 | 10.0 | 0.002 | 0.119 | 3.5 | 0.064 | 0.046 |
| Somatomotor B: Central 1 | 6.8 | 0.000 | 0.269 | 5.2 | 0.026 | 0.065 | 12.6 | 0.001 | 0.145 | 8.1 | 0.006 | 0.099 | 2.7 | 0.107 | 0.035 |
| Dorsal Attention A: Temporal Occipital 1 | 9.7 | 0.000 | 0.344 | 9.9 | 0.002 | 0.118 | 14.4 | 0.000 | 0.163 | 6.1 | 0.016 | 0.076 | 13.5 | 0.000 | 0.154 |
| Dorsal Attention A: Parietal Occipital 1 | 5.9 | 0.000 | 0.241 | 4.9 | 0.029 | 0.063 | 10.0 | 0.002 | 0.119 | 7.8 | 0.007 | 0.096 | 0.9 | 0.343 | 0.012 |
| Dorsal Attention A: Superior Parietal Lobule 1 | 8.0 | 0.000 | 0.303 | 5.7 | 0.020 | 0.071 | 10.7 | 0.002 | 0.126 | 7.6 | 0.007 | 0.093 | 4.6 | 0.035 | 0.059 |
| Dorsal Attention B: Post Central 1 | 7.8 | 0.000 | 0.296 | 3.1 | 0.081 | 0.041 | 15.2 | 0.000 | 0.170 | 7.6 | 0.007 | 0.093 | 0.1 | 0.710 | 0.002 |
| Dorsal Attention B: Post Central 2 | 12.6 | 0.000 | 0.406 | 3.3 | 0.075 | 0.042 | 6.2 | 0.015 | 0.078 | 8.3 | 0.005 | 0.101 | 16.0 | 0.000 | 0.178 |
| Dorsal Attention B: Post Central 3 | 11.6 | 0.000 | 0.386 | 8.1 | 0.006 | 0.099 | 16.1 | 0.000 | 0.179 | 8.7 | 0.004 | 0.105 | 12.0 | 0.001 | 0.140 |
| Dorsal Attention B: Frontal Eye Fields 1 | 11.5 | 0.000 | 0.383 | 4.3 | 0.041 | 0.055 | 16.2 | 0.000 | 0.179 | 7.5 | 0.008 | 0.092 | 3.2 | 0.076 | 0.042 |
| Salience Ventral Attention A: Parietal Operculum 1 | 10.1 | 0.000 | 0.353 | 5.3 | 0.024 | 0.067 | 17.6 | 0.000 | 0.192 | 10.7 | 0.002 | 0.126 | 1.3 | 0.256 | 0.017 |
| Salience Ventral Attention A: Insula: 1 | 13.0 | 0.000 | 0.413 | 5.4 | 0.023 | 0.068 | 23.3 | 0.000 | 0.239 | 9.6 | 0.003 | 0.114 | 1.2 | 0.274 | 0.016 |
| Salience Ventral Attention A: Insula: 2 | 12.3 | 0.000 | 0.400 | 4.2 | 0.043 | 0.054 | 13.4 | 0.000 | 0.153 | 7.3 | 0.008 | 0.090 | 5.3 | 0.024 | 0.067 |
| Salience Ventral Attention A: Parietal Medial 1 | 12.8 | 0.000 | 0.409 | 4.6 | 0.035 | 0.059 | 7.7 | 0.007 | 0.094 | 14.2 | 0.000 | 0.161 | 13.2 | 0.001 | 0.152 |
| Salience Ventral Attention A: Frontal Medial 1 | 12.0 | 0.000 | 0.393 | 4.8 | 0.031 | 0.061 | 10.0 | 0.002 | 0.119 | 7.6 | 0.007 | 0.093 | 7.6 | 0.007 | 0.093 |
| Salience Ventral Attention B: Lateral Prefrontal Corte | 16.6 | 0.000 | 0.473 | 4.6 | 0.036 | 0.058 | 14.7 | 0.000 | 0.166 | 11.7 | 0.001 | 0.136 | 14.3 | 0.000 | 0.162 |
| Salience Ventral Attention B: Medial Posterior Prefror | 11.1 | 0.000 | 0.374 | 2.1 | 0.154 | 0.027 | 12.3 | 0.001 | 0.142 | 9.7 | 0.003 | 0.116 | 1.8 | 0.188 | 0.023 |
| Limbic A Temporal Pole 1 | 7.8 | 0.000 | 0.295 | 4.4 | 0.040 | 0.056 | 16.1 | 0.000 | 0.178 | 7.3 | 0.009 | 0.090 | 3.6 | 0.063 | 0.046 |
| Limbic A: Temporal Pole 2 | 6.2 | 0.000 | 0.250 | 4.0 | 0.050 | 0.051 | 13.7 | 0.000 | 0.156 | 8.8 | 0.004 | 0.106 | 0.4 | 0.529 | 0.005 |
| Limbic B: Orbital Frontal Cortex 1 | 8.2 | 0.000 | 0.308 | 3.0 | 0.090 | 0.038 | 16.4 | 0.000 | 0.182 | 10.3 | 0.002 | 0.122 | 0.2 | 0.686 | 0.002 |
| Control A: Intraparietal Sulcus 1 | 10.3 | 0.000 | 0.357 | 5.1 | 0.027 | 0.064 | 12.7 | 0.001 | 0.147 | 7.4 | 0.008 | 0.091 | 5.9 | 0.017 | 0.074 |
| Control A: Lateral Prefrontal Cortex 1 | 9.7 | 0.000 | 0.345 | 3.4 | 0.067 | 0.045 | 12.0 | 0.001 | 0.139 | 6.9 | 0.010 | 0.086 | 3.3 | 0.073 | 0.043 |
| Control A: Lateral Prefrontal Cortex 2 | 10.5 | 0.000 | 0.361 | 1.7 | 0.191 | 0.023 | 5.0 | 0.028 | 0.064 | 5.6 | 0.021 | 0.070 | 6.8 | 0.011 | 0.084 |
| Control B: Lateral Prefrontal Cortexv 1 | 10.3 | 0.000 | 0.358 | 4.4 | 0.039 | 0.056 | 16.0 | 0.000 | 0.177 | 8.6 | 0.005 | 0.104 | 6.0 | 0.017 | 0.075 |
| Control C: Precuneus 1 | 6.3 | 0.000 | 0.253 | 2.8 | 0.101 | 0.036 | 7.5 | 0.008 | 0.092 | 12.3 | 0.001 | 0.143 | 0.9 | 0.357 | 0.011 |
| Control C: Precuneus 2 | 8.1 | 0.000 | 0.305 | 5.4 | 0.023 | 0.067 | 11.4 | 0.001 | 0.133 | 12.9 | 0.001 | 0.148 | 3.7 | 0.059 | 0.047 |
| Control C: Cingulate Posterior 1 | 6.2 | 0.000 | 0.250 | 4.2 | 0.044 | 0.054 | 8.9 | 0.004 | 0.107 | 9.5 | 0.003 | 0.114 | 1.9 | 0.167 | 0.026 |
| Default A: Dorsal Prefrontal Cortex 1 | 16.5 | 0.000 | 0.471 | 3.2 | 0.077 | 0.042 | 15.4 | 0.000 | 0.173 | 8.5 | 0.005 | 0.103 | 10.6 | 0.002 | 0.125 |
| Default A: Precuneus Posterior Cingulate Cortex1 | 6.7 | 0.000 | 0.266 | 3.8 | 0.054 | 0.049 | 8.3 | 0.005 | 0.101 | 11.8 | 0.001 | 0.138 | 2.8 | 0.100 | 0.036 |
| Default A: Medial Prefrontal Cortex 1 | 12.5 | 0.000 | 0.403 | 3.3 | 0.074 | 0.042 | 21.0 | 0.000 | 0.221 | 8.9 | 0.004 | 0.107 | 1.3 | 0.264 | 0.017 |
| Default B: Temp 1 | 9.5 | 0.000 | 0.338 | 5.8 | 0.018 | 0.073 | 15.9 | 0.000 | 0.176 | 8.2 | 0.005 | 0.100 | 3.1 | 0.083 | 0.040 |
| Default B: Temp 2 | 9.5 | 0.000 | 0.340 | 4.9 | 0.029 | 0.063 | 10.4 | 0.002 | 0.123 | 7.2 | 0.009 | 0.089 | 7.8 | 0.007 | 0.096 |
| Default B: Inferior Parietal Lobule 1 | 9.2 | 0.000 | 0.333 | 6.1 | 0.016 | 0.076 | 8.8 | 0.004 | 0.107 | 11.4 | 0.001 | 0.133 | 4.8 | 0.031 | 0.061 |
| Default B: Dorsal Prefrontal Cortex 1 | 15.3 | 0.000 | 0.452 | 3.6 | 0.060 | 0.047 | 15.3 | 0.000 | 0.171 | 8.8 | 0.004 | 0.106 | 3.9 | 0.051 | 0.050 |
| Default B: Lateral Prefrontal Cortex 1 | 10.8 | 0.000 | 0.369 | 3.8 | 0.055 | 0.049 | 12.9 | 0.001 | 0.149 | 7.1 | 0.009 | 0.088 | 5.9 | 0.017 | 0.074 |
| Default B: Ventral Prefrontal Cortex 1 | 11.5 | 0.000 | 0.383 | 2.8 | 0.100 | 0.036 | 20.9 | 0.000 | 0.220 | 10.8 | 0.002 | 0.128 | 0.0 | 0.971 | 0.000 |
| Default B: Ventral Prefrontal Cortex 2 | 10.3 | 0.000 | 0.357 | 4.8 | 0.031 | 0.061 | 15.1 | 0.000 | 0.170 | 10.4 | 0.002 | 0.123 | 2.5 | 0.119 | 0.033 |
| Default C: RetroSuperior Parietal Lobuleenial 1 | 7.4 | 0.000 | 0.286 | 2.7 | 0.108 | 0.035 | 11.1 | 0.001 | 0.130 | 13.5 | 0.000 | 0.155 | 0.7 | 0.422 | 0.009 |
| Default C: Parahippocampal Cortex 1 | 7.4 | 0.000 | 0.285 | 3.3 | 0.074 | 0.043 | 12.9 | 0.001 | 0.149 | 6.7 | 0.012 | 0.083 | 2.1 | 0.150 | 0.028 |
| Temporal Parietal 1 | 10.0 | 0.000 | 0.350 | 7.7 | 0.007 | 0.095 | 15.5 | 0.000 | 0.174 | 11.5 | 0.001 | 0.134 | 5.2 | 0.026 | 0.066 |
| Caudate | 5.9 | 0.000 | 0.241 | 3.3 | 0.074 | 0.043 | 6.8 | 0.011 | 0.085 | 6.5 | 0.013 | 0.081 | 2.8 | 0.098 | 0.037 |
| Putamen | 5.8 | 0.000 | 0.239 | 1.8 | 0.184 | 0.024 | 7.6 | 0.007 | 0.093 | 4.4 | 0.040 | 0.056 | 1.4 | 0.237 | 0.019 |
| Pallidum | 5.7 | 0.001 | 0.236 | 0.7 | 0.422 | 0.009 | 6.0 | 0.017 | 0.075 | 5.8 | 0.019 | 0.072 | 0.8 | 0.373 | 0.011 |
| Thalamus | 5.6 | 0.001 | 0.232 | 0.9 | 0.353 | 0.012 | 4.3 | 0.041 | 0.055 | 9.6 | 0.003 | 0.115 | 1.4 | 0.245 | 0.018 |

**Right Hemisphere**

| | Overall Model | | | Cortical Thickness | | | Age Group | | | HOMA-IR | | | Age Group x HOMA-IR | | |
|---|---|---|---|---|---|---|---|---|---|---|---|---|---|---|---|
| | F | p-FDR | $\eta^2_p$ | F | p | $\eta^2_p$ | F | p | $\eta^2_p$ | F | p | $\eta^2_p$ | F | p | $\eta^2_p$ |
| Visual Central: Extra Striate Cortex 1 | 4.1 | 0.005 | 0.183 | 4.6 | 0.035 | 0.059 | 5.0 | 0.028 | 0.064 | 8.6 | 0.005 | 0.104 | 3.5 | 0.064 | 0.045 |
| Visual Central: Extra Striate Cortex 2 | 2.9 | 0.030 | 0.134 | 2.8 | 0.096 | 0.037 | 4.6 | 0.035 | 0.058 | 8.3 | 0.005 | 0.101 | 0.0 | 0.896 | 0.000 |
| Visual Central: Extra Striate Cortex 3 | 5.0 | 0.001 | 0.214 | 6.8 | 0.011 | 0.084 | 8.3 | 0.005 | 0.100 | 9.6 | 0.003 | 0.115 | 1.0 | 0.310 | 0.014 |
| Visual Peripheral: Striate Cortex Calcarine 1 | 3.1 | 0.020 | 0.145 | 3.6 | 0.061 | 0.047 | 8.1 | 0.006 | 0.099 | 5.2 | 0.025 | 0.066 | 0.4 | 0.540 | 0.005 |
| Visual Peripheral: Extra Striate Inferior 1 | 7.0 | 0.000 | 0.274 | 4.9 | 0.029 | 0.062 | 9.1 | 0.004 | 0.109 | 8.3 | 0.005 | 0.100 | 4.1 | 0.045 | 0.053 |
| Visual Peripheral: Extra Striate Superior 1 | 6.2 | 0.000 | 0.251 | 7.6 | 0.007 | 0.093 | 11.6 | 0.001 | 0.135 | 9.1 | 0.004 | 0.109 | 2.9 | 0.092 | 0.038 |
| Somatomotor A: 1 | 8.2 | 0.000 | 0.306 | 3.3 | 0.071 | 0.043 | 13.4 | 0.000 | 0.153 | 10.2 | 0.002 | 0.121 | 1.9 | 0.171 | 0.025 |
| Somatomotor A: 2 | 8.1 | 0.000 | 0.305 | 5.9 | 0.017 | 0.074 | 15.0 | 0.000 | 0.169 | 9.8 | 0.002 | 0.117 | 5.9 | 0.018 | 0.073 |
| Somatomotor A: 3 | 8.5 | 0.000 | 0.315 | 3.7 | 0.058 | 0.048 | 9.9 | 0.002 | 0.118 | 9.0 | 0.004 | 0.109 | 7.0 | 0.010 | 0.086 |
| Somatomotor A: 4 | 8.8 | 0.000 | 0.322 | 3.8 | 0.055 | 0.049 | 9.3 | 0.003 | 0.111 | 10.6 | 0.002 | 0.125 | 3.8 | 0.056 | 0.049 |
| Somatomotor B: Auditory 1 | 11.8 | 0.000 | 0.389 | 10.0 | 0.002 | 0.119 | 11.5 | 0.001 | 0.135 | 14.1 | 0.000 | 0.160 | 9.9 | 0.002 | 0.118 |
| Somatomotor B: S2 1 | 7.8 | 0.000 | 0.297 | 3.5 | 0.064 | 0.045 | 14.8 | 0.000 | 0.166 | 8.8 | 0.004 | 0.106 | 0.8 | 0.381 | 0.010 |
| Somatomotor B: S2 2 | 12.0 | 0.000 | 0.393 | 6.9 | 0.011 | 0.085 | 20.5 | 0.000 | 0.217 | 12.2 | 0.001 | 0.141 | 3.3 | 0.071 | 0.043 |
| Somatomotor B: Central 1 | 6.0 | 0.000 | 0.246 | 4.5 | 0.037 | 0.058 | 8.1 | 0.006 | 0.099 | 8.7 | 0.004 | 0.105 | 2.7 | 0.103 | 0.036 |
| Dorsal Attention A: Temporal Occipital 1 | 5.2 | 0.001 | 0.220 | 6.8 | 0.011 | 0.084 | 12.8 | 0.001 | 0.147 | 8.1 | 0.006 | 0.099 | 0.5 | 0.481 | 0.007 |
| Dorsal Attention A: Parietal Occipital 1 | 7.4 | 0.000 | 0.284 | 8.5 | 0.005 | 0.103 | 13.8 | 0.000 | 0.157 | 8.9 | 0.004 | 0.107 | 2.3 | 0.130 | 0.031 |
| Dorsal Attention A: Superior Parietal Lobule 1 | 9.2 | 0.000 | 0.331 | 6.8 | 0.011 | 0.085 | 14.3 | 0.000 | 0.162 | 12.5 | 0.001 | 0.144 | 4.6 | 0.035 | 0.059 |
| Dorsal Attention B: Post Central 1 | 7.9 | 0.000 | 0.300 | 3.4 | 0.070 | 0.044 | 7.1 | 0.010 | 0.087 | 9.1 | 0.003 | 0.110 | 2.3 | 0.134 | 0.030 |
| Dorsal Attention B: Post Central 2 | 11.2 | 0.000 | 0.377 | 7.1 | 0.010 | 0.087 | 9.7 | 0.003 | 0.115 | 9.4 | 0.003 | 0.113 | 15.8 | 0.000 | 0.176 |
| Dorsal Attention B: Frontal Eye Fields 1 | 15.3 | 0.000 | 0.453 | 2.3 | 0.137 | 0.030 | 9.6 | 0.003 | 0.115 | 7.3 | 0.009 | 0.090 | 13.2 | 0.001 | 0.152 |
| Salience Ventral Attention A: Parietal Operculum 1 | 11.6 | 0.000 | 0.386 | 6.0 | 0.016 | 0.076 | 22.8 | 0.000 | 0.236 | 12.3 | 0.001 | 0.142 | 1.6 | 0.211 | 0.021 |
| Salience Ventral Attention A: Insula: 1 | 10.1 | 0.000 | 0.353 | 4.6 | 0.035 | 0.059 | 18.4 | 0.000 | 0.200 | 9.9 | 0.002 | 0.118 | 0.2 | 0.637 | 0.003 |
| Salience Ventral Attention A: Parietal Medial 1 | 11.7 | 0.000 | 0.388 | 2.9 | 0.094 | 0.038 | 6.1 | 0.016 | 0.076 | 10.2 | 0.002 | 0.121 | 13.8 | 0.000 | 0.157 |
| Salience Ventral Attention A: Frontal Medial 1 | 13.0 | 0.000 | 0.413 | 5.3 | 0.024 | 0.067 | 9.6 | 0.003 | 0.115 | 9.4 | 0.003 | 0.113 | 9.9 | 0.002 | 0.118 |
| Salience Ventral Attention B: Inferior Parietal Lobule | 9.6 | 0.000 | 0.340 | 5.7 | 0.020 | 0.072 | 18.4 | 0.000 | 0.199 | 8.7 | 0.004 | 0.106 | 1.2 | 0.278 | 0.016 |
| Salience Ventral Attention B: Lateral Prefrontal Cor | 12.1 | 0.000 | 0.396 | 5.4 | 0.023 | 0.068 | 18.7 | 0.000 | 0.201 | 7.6 | 0.007 | 0.093 | 5.5 | 0.022 | 0.069 |
| Salience Ventral Attention B: Medial Posterior Prefr | 12.7 | 0.000 | 0.408 | 1.6 | 0.217 | 0.021 | 15.6 | 0.000 | 0.174 | 8.9 | 0.004 | 0.107 | 0.9 | 0.345 | 0.012 |
| Limbic A: Temporal Pole 1 | 8.1 | 0.000 | 0.305 | 4.0 | 0.049 | 0.051 | 15.7 | 0.000 | 0.175 | 8.4 | 0.005 | 0.102 | 3.3 | 0.075 | 0.042 |
| Limbic B: Orbital Frontal Cortex 1 | 7.7 | 0.000 | 0.293 | 3.2 | 0.079 | 0.041 | 16.9 | 0.000 | 0.186 | 8.7 | 0.004 | 0.105 | 0.1 | 0.794 | 0.001 |
| Control A: Intraparietal Sulcus 1 | 7.0 | 0.000 | 0.275 | 4.9 | 0.030 | 0.062 | 8.5 | 0.005 | 0.103 | 5.4 | 0.023 | 0.068 | 5.7 | 0.020 | 0.071 |
| Control A: Lateral Prefrontal Cortex 1 | 9.2 | 0.000 | 0.333 | 4.5 | 0.036 | 0.058 | 20.1 | 0.000 | 0.214 | 9.0 | 0.004 | 0.109 | 0.0 | 0.993 | 0.000 |
| Control A: Lateral Prefrontal Cortex 2 | 10.1 | 0.000 | 0.353 | 5.2 | 0.025 | 0.066 | 15.9 | 0.000 | 0.177 | 8.9 | 0.004 | 0.108 | 3.1 | 0.082 | 0.040 |
| Control B: Temporal 1 | 7.9 | 0.000 | 0.298 | 4.0 | 0.048 | 0.052 | 17.2 | 0.000 | 0.188 | 10.7 | 0.002 | 0.127 | 0.1 | 0.751 | 0.001 |
| Control B: inferior parietal lobule 1 | 10.9 | 0.000 | 0.370 | 5.2 | 0.026 | 0.065 | 14.6 | 0.000 | 0.165 | 11.3 | 0.001 | 0.132 | 6.2 | 0.015 | 0.078 |
| Control B: Lateral Prefrontal Cortexd 1 | 12.9 | 0.000 | 0.411 | 5.5 | 0.022 | 0.069 | 13.3 | 0.000 | 0.153 | 10.3 | 0.002 | 0.122 | 9.0 | 0.004 | 0.109 |
| Control B: Lateral Prefrontal Cortexv 1 | 11.2 | 0.000 | 0.378 | 6.3 | 0.014 | 0.079 | 19.3 | 0.000 | 0.207 | 11.3 | 0.001 | 0.133 | 5.1 | 0.026 | 0.065 |
| Control C: Cingulate Posterior 1 | 7.8 | 0.000 | 0.296 | 3.5 | 0.064 | 0.046 | 11.2 | 0.001 | 0.132 | 11.4 | 0.001 | 0.134 | 1.8 | 0.181 | 0.024 |
| Control C: Precuneus 1 | 7.3 | 0.000 | 0.284 | 6.2 | 0.015 | 0.078 | 12.5 | 0.001 | 0.144 | 12.9 | 0.001 | 0.149 | 2.1 | 0.152 | 0.027 |
| Default A: Inferior Parietal Lobule 1 | 10.1 | 0.000 | 0.352 | 5.8 | 0.019 | 0.072 | 7.0 | 0.010 | 0.087 | 10.7 | 0.002 | 0.127 | 9.0 | 0.004 | 0.108 |
| Default A: Dorsal Prefrontal Cortex 1 | 13.6 | 0.000 | 0.424 | 4.9 | 0.030 | 0.062 | 19.3 | 0.000 | 0.207 | 6.1 | 0.016 | 0.076 | 6.7 | 0.012 | 0.082 |
| Default A: Precuneus Posterior Cingulate Cortex 1 | 6.3 | 0.000 | 0.255 | 5.4 | 0.023 | 0.068 | 12.5 | 0.001 | 0.144 | 11.6 | 0.001 | 0.136 | 0.6 | 0.432 | 0.008 |
| Default A: Medial Prefrontal Cortex 1 | 13.2 | 0.000 | 0.417 | 3.2 | 0.079 | 0.041 | 21.3 | 0.000 | 0.223 | 10.5 | 0.002 | 0.125 | 1.1 | 0.304 | 0.014 |
| Default B: Dorsal Prefrontal Cortex 1 | 15.8 | 0.000 | 0.460 | 4.7 | 0.033 | 0.060 | 17.2 | 0.000 | 0.189 | 10.1 | 0.002 | 0.120 | 7.0 | 0.010 | 0.087 |
| Default B: Ventral Prefrontal Cortex 1 | 10.5 | 0.000 | 0.363 | 3.9 | 0.053 | 0.049 | 17.3 | 0.000 | 0.189 | 11.9 | 0.001 | 0.138 | 1.5 | 0.219 | 0.020 |
| Default B: Ventral Prefrontal Cortex 2 | 11.1 | 0.000 | 0.376 | 3.5 | 0.066 | 0.045 | 16.4 | 0.000 | 0.181 | 9.6 | 0.003 | 0.115 | 1.8 | 0.178 | 0.024 |
| Default C: RetroSuperior Parietal Lobuleenial 1 | 6.6 | 0.000 | 0.262 | 4.4 | 0.039 | 0.056 | 9.8 | 0.003 | 0.116 | 9.6 | 0.003 | 0.115 | 4.1 | 0.047 | 0.052 |
| Default C: Parahippocampal Cortex 1 | 7.4 | 0.000 | 0.287 | 7.5 | 0.008 | 0.092 | 14.0 | 0.000 | 0.159 | 9.2 | 0.003 | 0.110 | 5.2 | 0.025 | 0.066 |
| Temporal Parietal 1 | 11.4 | 0.000 | 0.381 | 7.5 | 0.008 | 0.092 | 18.6 | 0.000 | 0.201 | 8.0 | 0.006 | 0.097 | 6.0 | 0.016 | 0.075 |
| Temporal Parietal 2 | 11.4 | 0.000 | 0.382 | 7.7 | 0.007 | 0.094 | 13.6 | 0.000 | 0.155 | 13.5 | 0.000 | 0.155 | 5.6 | 0.020 | 0.071 |
| Temporal Parietal 3 | 7.2 | 0.000 | 0.281 | 5.9 | 0.018 | 0.074 | 11.3 | 0.001 | 0.132 | 7.9 | 0.006 | 0.096 | 3.1 | 0.083 | 0.040 |
| Caudate | 4.7 | 0.002 | 0.204 | 3.0 | 0.086 | 0.039 | 5.3 | 0.025 | 0.067 | 7.2 | 0.009 | 0.088 | 2.1 | 0.149 | 0.030 |
| Putamen | 5.5 | 0.001 | 0.229 | 2.4 | 0.125 | 0.032 | 7.1 | 0.010 | 0.087 | 5.6 | 0.021 | 0.070 | 1.7 | 0.193 | 0.020 |
| Pallidum | 5.8 | 0.001 | 0.237 | 1.3 | 0.263 | 0.017 | 6.7 | 0.012 | 0.083 | 5.8 | 0.019 | 0.073 | 1.1 | 0.288 | 0.020 |
| Thalamus | 5.5 | 0.001 | 0.228 | 0.9 | 0.334 | 0.013 | 4.1 | 0.045 | 0.053 | 10.4 | 0.002 | 0.123 | 1.2 | 0.273 | 0.020 |

Table S3. Post hoc analyses of significant age group x HOMA-IR interactions; separate general linear models of $CMR_{GLC}$ for younger and older adults. The models include cortical thickness as a covariate. Δ% is the percentage change in $CMR_{GLC}$ from a 10% change in HOMA-IR, calculated from the slope of the regression lines.

**Left Hemisphere**

| | Age Group x HOMA-IR | | | Post-Hoc Younger | | | Post-Hoc Older | | |
|---|---|---|---|---|---|---|---|---|---|
| | F | p | $\eta^2_p$ | F | p | Δ% | F | p | Δ% |
| Visual Central: Extra Striate Cortex 1 | 1.1 | 0.298 | 0.015 | | | | | | |
| Visual Central: Extra Striate Cortex 2 | 0.2 | 0.635 | 0.003 | | | | | | |
| Visual Central: Striate Cortex 1 | 0.3 | 0.603 | 0.004 | | | | | | |
| Visual Central: Extra Striate Cortex 3 | 1.2 | 0.274 | 0.016 | | | | | | |
| Visual Peripheral: Extra Striate Inferior 1 | 0.5 | 0.491 | 0.006 | | | | | | |
| Visual Peripheral: Striate Cortex Calcarine 1 | 9.4 | 0.003 | 0.113 | 9.5 | 0.004 | -6.1 | 0.1 | 0.799 | 0.0 |
| Visual Peripheral: Extra Striate CortexSup 1 | 2.0 | 0.162 | 0.026 | | | | | | |
| Somatomotor A: 1 | 7.3 | 0.009 | 0.090 | 7.1 | 0.012 | -5.1 | 0.3 | 0.574 | -0.5 |
| Somatomotor A: 2 | 5.1 | 0.027 | 0.065 | 7.9 | 0.008 | -5.1 | 1.0 | 0.315 | -1.0 |
| Somatomotor B: Auditory 1 | 5.4 | 0.022 | 0.069 | 10.8 | 0.002 | -6.3 | 0.8 | 0.366 | -1.3 |
| Somatomotor B: S2 1 | 1.4 | 0.244 | 0.018 | | | | | | |
| Somatomotor B: S2 2 | 3.5 | 0.064 | 0.046 | | | | | | |
| Somatomotor B: Central 1 | 2.7 | 0.107 | 0.035 | | | | | | |
| Dorsal Attention A: Temporal Occipital 1 | 13.5 | 0.000 | 0.154 | 8.5 | 0.006 | -5.2 | 0.4 | 0.509 | -0.5 |
| Dorsal Attention A: Parietal Occipital 1 | 0.9 | 0.343 | 0.012 | | | | | | |
| Dorsal Attention A: Superior Parietal Lobule 1 | 4.6 | 0.035 | 0.059 | 7.6 | 0.009 | -6.0 | 0.1 | 0.819 | -0.9 |
| Dorsal Attention B: Post Central 1 | 0.1 | 0.710 | 0.002 | | | | | | |
| Dorsal Attention B: Post Central 2 | 16.0 | 0.000 | 0.178 | 6.1 | 0.019 | -5.1 | 1.0 | 0.315 | -0.9 |
| Dorsal Attention B: Post Central 3 | 12.0 | 0.001 | 0.140 | 8.6 | 0.006 | -5.1 | 0.0 | 0.925 | -0.6 |
| Dorsal Attention B: Frontal Eye Fields 1 | 3.2 | 0.076 | 0.042 | | | | | | |
| Salience Ventral Attention A: Parietal Operculum 1 | 1.3 | 0.256 | 0.017 | | | | | | |
| Salience Ventral Attention A: Insula: 1 | 1.2 | 0.274 | 0.016 | | | | | | |
| Salience Ventral Attention A: Insula: 2 | 5.3 | 0.024 | 0.067 | 7.4 | 0.010 | -6.4 | 0.6 | 0.425 | -1.4 |
| Salience Ventral Attention A: Parietal Medial 1 | 13.2 | 0.001 | 0.152 | 11.1 | 0.002 | -6.0 | 2.1 | 0.155 | -1.7 |
| Salience Ventral Attention A: Frontal Medial 1 | 7.6 | 0.007 | 0.093 | 6.3 | 0.017 | -5.8 | 0.0 | 0.915 | -2.2 |
| Salience Ventral Attention B: Lateral Prefrontal Cortex 1 | 14.3 | 0.000 | 0.162 | 7.7 | 0.009 | -6.1 | 2.0 | 0.164 | -2.0 |
| Salience Ventral Attention B: Medial Posterior Prefrontal | 1.8 | 0.188 | 0.023 | | | | | | |
| Limbic A Temporal Pole 1 | 3.6 | 0.063 | 0.046 | | | | | | |
| Limbic A: Temporal Pole 2 | 0.4 | 0.529 | 0.005 | | | | | | |
| Limbic B: Orbital Frontal Cortex 1 | 0.2 | 0.686 | 0.002 | | | | | | |
| Control A: Intraparietal Sulcus 1 | 5.9 | 0.017 | 0.074 | 6.6 | 0.015 | -5.7 | 0.2 | 0.655 | -0.8 |
| Control A: Lateral Prefrontal Cortex 1 | 3.3 | 0.073 | 0.043 | | | | | | |
| Control A: Lateral Prefrontal Cortex 2 | 6.8 | 0.011 | 0.084 | 3.6 | 0.065 | -5.5 | 1.2 | 0.288 | -1.6 |
| Control B: Lateral Prefrontal Cortexv 1 | 6.0 | 0.017 | 0.075 | 7.3 | 0.011 | -7.0 | 0.8 | 0.368 | -1.5 |
| Control C: Precuneus 1 | 0.9 | 0.357 | 0.011 | | | | | | |
| Control C: Precuneus 2 | 3.7 | 0.059 | 0.047 | | | | | | |
| Control C: Cingulate Posterior 1 | 1.9 | 0.167 | 0.026 | 6.5 | 0.015 | -5.5 | 0.7 | 0.423 | -1.3 |
| Default A: Dorsal Prefrontal Cortex 1 | 10.6 | 0.002 | 0.125 | 6.3 | 0.017 | -6.2 | 1.5 | 0.226 | -2.2 |
| Default A: Precuneus Posterior Cingulate Cortex1 | 2.8 | 0.100 | 0.036 | | | | | | |
| Default A: Medial Prefrontal Cortex 1 | 1.3 | 0.264 | 0.017 | | | | | | |
| Default B: Temp 1 | 3.1 | 0.083 | 0.040 | | | | | | |
| Default B: Temp 2 | 7.8 | 0.007 | 0.096 | 6.5 | 0.016 | -4.9 | 0.2 | 0.673 | -1.0 |
| Default B: Inferior Parietal Lobule 1 | 4.8 | 0.031 | 0.061 | 8.9 | 0.005 | -6.1 | 0.7 | 0.424 | -1.3 |
| Default B: Dorsal Prefrontal Cortex 1 | 3.9 | 0.051 | 0.050 | | | | | | |
| Default B: Lateral Prefrontal Cortex 1 | 5.9 | 0.017 | 0.074 | 5.5 | 0.025 | -5.5 | 0.4 | 0.509 | -1.5 |
| Default B: Ventral Prefrontal Cortex 1 | 0.0 | 0.971 | 0.000 | | | | | | |
| Default B: Ventral Prefrontal Cortex 2 | 2.5 | 0.119 | 0.033 | | | | | | |
| Default C: RetroSuperior Parietal Lobuleenial 1 | 0.7 | 0.422 | 0.009 | | | | | | |
| Default C: Parahippocampal Cortex 1 | 2.1 | 0.150 | 0.028 | | | | | | |
| Temporal Parietal 1 | 5.2 | 0.026 | 0.066 | 10.8 | 0.002 | -6.1 | 0.3 | 0.585 | -0.8 |
| Caudate | 2.8 | 0.098 | 0.037 | | | | | | |
| Putamen | 1.4 | 0.237 | 0.019 | | | | | | |
| Pallidum | 0.8 | 0.373 | 0.011 | | | | | | |
| Thalamus | 1.4 | 0.245 | 0.018 | | | | | | |

**Right Hemisphere**

| | Age Group x HOMA-IR | | | Post-Hoc Younger | | | Post-Hoc Older | | |
|---|---|---|---|---|---|---|---|---|---|
| | F | p | $\eta^2_p$ | F | p | Δ% | F | p | Δ% |
| Visual Central: Extra Striate Cortex 1 | 3.5 | 0.064 | 0.045 | | | | | | |
| Visual Central: Extra Striate Cortex 2 | 0.0 | 0.896 | 0.000 | | | | | | |
| Visual Central: Extra Striate Cortex 3 | 1.0 | 0.310 | 0.014 | | | | | | |
| Visual Peripheral: Striate Cortex Calcarine 1 | 0.4 | 0.540 | 0.005 | | | | | | |
| Visual Peripheral: Extra Striate Inferior 1 | 4.1 | 0.045 | 0.053 | 8.6 | 0.006 | -5.9 | 0.3 | 0.562 | -1.3 |
| Visual Peripheral: Extra Striate Superior 1 | 2.9 | 0.092 | 0.038 | | | | | | |
| Somatomotor A: 1 | 1.9 | 0.171 | 0.025 | | | | | | |
| Somatomotor A: 2 | 5.9 | 0.018 | 0.073 | 6.7 | 0.014 | -5.2 | 0.5 | 0.485 | -1.3 |
| Somatomotor A: 3 | 7.0 | 0.010 | 0.086 | 7.5 | 0.010 | -5.2 | 0.7 | 0.400 | -1.1 |
| Somatomotor A: 4 | 3.8 | 0.056 | 0.049 | | | | | | |
| Somatomotor B: Auditory 1 | 9.9 | 0.002 | 0.118 | 15.4 | 0.000 | -5.7 | 0.4 | 0.535 | -0.3 |
| Somatomotor B: S2 1 | 0.8 | 0.381 | 0.010 | | | | | | |
| Somatomotor B: S2 2 | 3.3 | 0.071 | 0.043 | | | | | | |
| Somatomotor B: Central 1 | 2.7 | 0.103 | 0.036 | | | | | | |
| Dorsal Attention A: Temporal Occipital 1 | 0.5 | 0.481 | 0.007 | | | | | | |
| Dorsal Attention A: Parietal Occipital 1 | 2.3 | 0.130 | 0.031 | | | | | | |
| Dorsal Attention A: Superior Parietal Lobule 1 | 4.6 | 0.035 | 0.059 | 10.4 | 0.003 | -5.4 | 0.9 | 0.359 | -1.3 |
| Dorsal Attention B: Post Central 1 | 2.3 | 0.134 | 0.030 | | | | | | |
| Dorsal Attention B: Post Central 2 | 15.8 | 0.000 | 0.176 | 9.3 | 0.004 | -4.7 | 0.1 | 0.784 | -1.7 |
| Dorsal Attention B: Frontal Eye Fields 1 | 13.2 | 0.001 | 0.152 | 4.9 | 0.034 | -5.2 | 0.9 | 0.341 | -1.6 |
| Salience Ventral Attention A: Parietal Operculum 1 | 1.6 | 0.211 | 0.021 | | | | | | |
| Salience Ventral Attention A: Insula: 1 | 0.2 | 0.637 | 0.003 | | | | | | |
| Salience Ventral Attention A: Parietal Medial 1 | 13.8 | 0.000 | 0.157 | 7.6 | 0.009 | -4.9 | 1.9 | 0.177 | -2.0 |
| Salience Ventral Attention A: Frontal Medial 1 | 9.9 | 0.002 | 0.118 | 8.6 | 0.006 | -5.6 | 0.3 | 0.562 | -1.8 |
| Salience Ventral Attention B: Inferior Parietal Lobule 1 | 1.2 | 0.278 | 0.016 | | | | | | |
| Salience Ventral Attention B: Lateral Prefrontal Cortex 1 | 5.5 | 0.022 | 0.069 | 7.3 | 0.011 | -6.3 | 0.0 | 0.976 | -0.9 |
| Salience Ventral Attention B: Medial Posterior Prefrontal 1 | 0.9 | 0.345 | 0.012 | | | | | | |
| Limbic A: Temporal Pole 1 | 3.3 | 0.075 | 0.042 | | | | | | |
| Limbic B: Orbital Frontal Cortex 1 | | | | | | | | | |
| | 0.1 | 0.794 | 0.001 | | | | | | |
| Control A: Intraparietal Sulcus 1 | 5.7 | 0.020 | 0.071 | 6.4 | 0.017 | -6.3 | 0.0 | 0.950 | -1.5 |
| Control A: Lateral Prefrontal Cortex 1 | 0.0 | 0.993 | 0.000 | | | | | | |
| Control A: Lateral Prefrontal Cortex 2 | 3.1 | 0.082 | 0.040 | | | | | | |
| Control B: Temporal 1 | 0.1 | 0.751 | 0.001 | | | | | | |
| Control B: inferior parietal lobule 1 | 6.2 | 0.015 | 0.078 | 11.1 | 0.002 | -7.3 | 0.7 | 0.417 | -2.0 |
| Control B: Lateral Prefrontal Cortexd 1 | 9.0 | 0.004 | 0.109 | 7.4 | 0.010 | -7.7 | 0.0 | 0.832 | -2.0 |
| Control B: Lateral Prefrontal Cortexv 1 | 5.1 | 0.026 | 0.065 | 11.2 | 0.002 | -5.9 | 1.2 | 0.279 | -1.6 |
| Control C: Cingulate Posterior 1 | 1.8 | 0.181 | 0.024 | | | | | | |
| Control C: Precuneus 1 | 2.1 | 0.152 | 0.027 | | | | | | |
| Default A: Inferior Parietal Lobule 1 | 9.0 | 0.004 | 0.108 | 9.3 | 0.005 | -5.5 | 0.2 | 0.660 | -2.0 |
| Default A: Dorsal Prefrontal Cortex 1 | 6.7 | 0.012 | 0.082 | 5.8 | 0.022 | -7.0 | 0.1 | 0.722 | -1.8 |
| Default A: Precuneus Posterior Cingulate Cortex 1 | 0.6 | 0.432 | 0.008 | | | | | | |
| Default A: Medial Prefrontal Cortex 1 | 1.1 | 0.304 | 0.014 | | | | | | |
| Default B: Dorsal Prefrontal Cortex 1 | 7.0 | 0.010 | 0.087 | 7.0 | 0.012 | -6.6 | 0.4 | 0.506 | -0.9 |
| Default B: Ventral Prefrontal Cortex 1 | 1.5 | 0.219 | 0.020 | | | | | | |
| Default B: Ventral Prefrontal Cortex 2 | 1.8 | 0.178 | 0.024 | | | | | | |
| Default C: RetroSuperior Parietal Lobuleenial 1 | 4.1 | 0.047 | 0.052 | 9.2 | 0.005 | -7.0 | 0.7 | 0.393 | -1.1 |
| Default C: Parahippocampal Cortex 1 | 5.2 | 0.025 | 0.066 | 9.9 | 0.004 | -3.9 | 0.1 | 0.796 | -0.8 |
| Temporal Parietal 1 | 6.0 | 0.016 | 0.075 | 8.5 | 0.006 | -3.3 | 0.0 | 0.958 | -0.9 |
| Temporal Parietal 2 | 5.6 | 0.020 | 0.071 | 12.8 | 0.001 | -4.0 | 0.9 | 0.359 | -0.9 |
| Temporal Parietal 3 | 3.1 | 0.083 | 0.040 | | | | | | |
| Caudate | 2.1 | 0.149 | 0.028 | | | | | | |
| Putamen | 1.7 | 0.193 | 0.023 | | | | | | |
| Pallidum | 1.1 | 0.288 | 0.015 | | | | | | |
| Thalamus | 1.2 | 0.273 | 0.016 | | | | | | |

Table S4. General linear models of age group, HOMA-IR and age group x HOMA-IR effects on network $CMR_{GLC}$, including cortical thickness as a covariate. Includes post-hoc analyses of significant interactions as separate GLM for younger and older adults separately.

| | Overall Model | | | Cortical Thickness | | | Age Cat | | | HOMA-IR | | | Age Group x HOMA-IR | | | Post hoc - Younger | | | Post hoc Older | | |
|---|---|---|---|---|---|---|---|---|---|---|---|---|---|---|---|---|---|---|---|---|---|
| | F | p-FDR | $\eta^2_p$ | F | p | $\eta^2_p$ | F | p | $\eta^2_p$ | F | p | $\eta^2_p$ | F | p | $\eta^2_p$ | F | p | $\eta^2_p$ | F | p | $\eta^2_p$ |
| Visual Central | 3.8 | 0.007 | 0.171 | 1.2 | 0.283 | 0.016 | 6.5 | 0.013 | 0.081 | 6.5 | 0.013 | 0.081 | 7.7 | 0.007 | 0.095 | 7.7 | 0.009 | 0.188 | 0.4 | 0.523 | 0.010 |
| Visual Peripheral | 6.3 | 0.000 | 0.254 | 4.0 | 0.048 | 0.052 | 10.2 | 0.002 | 0.121 | 10.2 | 0.002 | 0.121 | 8.4 | 0.005 | 0.101 | 9.0 | 0.005 | 0.215 | 0.1 | 0.749 | 0.003 |
| Somatomotor A | 10.0 | 0.000 | 0.352 | 6.8 | 0.011 | 0.084 | 10.5 | 0.002 | 0.124 | 10.5 | 0.002 | 0.124 | 10.8 | 0.002 | 0.128 | 8.8 | 0.006 | 0.210 | 0.9 | 0.351 | 0.022 |
| Somatomotor B | 11.4 | 0.000 | 0.381 | 6.3 | 0.014 | 0.079 | 14.5 | 0.000 | 0.163 | 14.5 | 0.000 | 0.163 | 11.6 | 0.001 | 0.136 | 14.2 | 0.001 | 0.300 | 0.5 | 0.488 | 0.012 |
| Dors Attention A | 8.6 | 0.000 | 0.318 | 6.0 | 0.017 | 0.075 | 10.6 | 0.002 | 0.125 | 10.6 | 0.002 | 0.125 | 9.1 | 0.004 | 0.109 | 8.8 | 0.006 | 0.210 | 0.0 | 0.858 | 0.001 |
| Dors Attention B | 13.8 | 0.000 | 0.427 | 13.0 | 0.001 | 0.149 | 7.8 | 0.007 | 0.095 | 7.8 | 0.007 | 0.095 | 8.1 | 0.006 | 0.099 | 7.1 | 0.012 | 0.177 | 0.3 | 0.606 | 0.007 |
| Salience Ventral Attention A | 13.0 | 0.000 | 0.412 | 6.5 | 0.013 | 0.080 | 10.3 | 0.002 | 0.122 | 10.3 | 0.002 | 0.122 | 10.5 | 0.002 | 0.124 | 9.7 | 0.004 | 0.227 | 0.5 | 0.490 | 0.012 |
| Salience Ventral Attention B | 13.8 | 0.000 | 0.427 | 5.8 | 0.018 | 0.073 | 12.0 | 0.001 | 0.139 | 12.0 | 0.001 | 0.139 | 10.6 | 0.002 | 0.125 | 7.5 | 0.010 | 0.185 | 1.4 | 0.249 | 0.033 |
| Limbic A | 8.1 | 0.000 | 0.304 | 3.8 | 0.054 | 0.049 | 16.2 | 0.000 | 0.180 | 16.2 | 0.000 | 0.180 | 8.5 | 0.005 | 0.103 | 6.8 | 0.014 | 0.170 | 1.9 | 0.175 | 0.046 |
| Limbic B | 8.0 | 0.000 | 0.301 | 0.1 | 0.707 | 0.002 | 16.7 | 0.000 | 0.184 | 16.7 | 0.000 | 0.184 | 9.5 | 0.003 | 0.114 | 7.2 | 0.011 | 0.179 | 0.3 | 0.586 | 0.007 |
| Control A | 11.0 | 0.000 | 0.372 | 6.4 | 0.014 | 0.079 | 9.1 | 0.003 | 0.110 | 9.1 | 0.003 | 0.110 | 7.3 | 0.008 | 0.090 | 6.8 | 0.013 | 0.172 | 0.1 | 0.708 | 0.004 |
| Control B | 11.5 | 0.000 | 0.384 | 6.8 | 0.011 | 0.084 | 14.5 | 0.000 | 0.164 | 14.5 | 0.000 | 0.164 | 10.5 | 0.002 | 0.124 | 8.6 | 0.006 | 0.207 | 0.3 | 0.576 | 0.008 |
| Control C | 9.9 | 0.000 | 0.349 | 6.6 | 0.012 | 0.082 | 9.8 | 0.002 | 0.117 | 9.8 | 0.002 | 0.117 | 12.5 | 0.001 | 0.145 | 12.2 | 0.001 | 0.270 | 0.9 | 0.362 | 0.021 |
| Default A | 12.4 | 0.000 | 0.401 | 8.5 | 0.005 | 0.103 | 11.4 | 0.001 | 0.133 | 11.4 | 0.001 | 0.133 | 10.3 | 0.002 | 0.122 | 8.9 | 0.005 | 0.212 | 0.4 | 0.522 | 0.010 |
| Default B | 13.7 | 0.000 | 0.425 | 6.6 | 0.012 | 0.082 | 11.8 | 0.001 | 0.137 | 11.8 | 0.001 | 0.137 | 10.8 | 0.002 | 0.128 | 8.0 | 0.008 | 0.195 | 0.5 | 0.481 | 0.012 |
| Default C | 8.1 | 0.000 | 0.303 | 5.1 | 0.027 | 0.064 | 11.4 | 0.001 | 0.133 | 11.4 | 0.001 | 0.133 | 9.7 | 0.003 | 0.116 | 9.0 | 0.005 | 0.214 | 0.5 | 0.464 | 0.013 |
| Temporal Parietal | 11.4 | 0.000 | 0.382 | 7.6 | 0.007 | 0.093 | 12.5 | 0.001 | 0.144 | 12.5 | 0.001 | 0.144 | 11.9 | 0.001 | 0.139 | 12.8 | 0.001 | 0.279 | 0.2 | 0.644 | 0.005 |
| Subcortical | 5.8 | 0.000 | 0.237 | 1.7 | 0.202 | 0.022 | 5.7 | 0.020 | 0.071 | 5.7 | 0.020 | 0.071 | 8.4 | 0.005 | 0.102 | 6.0 | 0.020 | 0.154 | 2.5 | 0.119 | 0.060 |

#### *3. Principal Component Analysis for Cognitive Variables*

Table S5. Principal component analysis (loadings) of cognitive variables identifying five components explaining 81% of the variance.

| | Component | | | | |
|---|---|---|---|---|---|
| | 1 | 2 | 3 | 4 | 5 |
| Variance Explained (%) | 18.3 | 17.8 | 17.0 | 14.1 | 13.8 |
| HVLT: Delayed recall (total) | -0.209 | 0.064 | 0.058 | 0.266 | 0.779 |
| HVLT: Recognition discrimination index | -0.005 | -0.256 | -0.130 | -0.029 | 0.748 |
| Digit Span: Forward (longest) | 0.124 | 0.052 | 0.052 | 0.888 | -0.010 |
| Digit Span: Backwards (longest) | -0.160 | -0.144 | -0.094 | 0.817 | 0.212 |
| Category Switch: Reaction time in switch trials | 0.848 | 0.339 | 0.191 | -0.087 | -0.039 |
| Category Switch: SSRT | 0.938 | 0.105 | 0.075 | 0.042 | -0.084 |
| Digit Substitution: Correct count | -0.260 | -0.915 | -0.165 | 0.057 | 0.047 |
| Digit Substitution: Seconds per correct count | 0.156 | 0.916 | 0.017 | -0.024 | -0.040 |
| Stop Signal: Mean stop signal delays | 0.118 | 0.045 | 0.944 | 0.038 | 0.034 |
| Stop Signal: Stop signal reaction time | 0.114 | 0.120 | 0.930 | -0.069 | -0.063 |
| WASI FSIQ2 T-score | 0.456 | 0.206 | 0.097 | 0.016 | 0.534 |

Table S6. General Linear Models predicting each of the 5 principal components from network $CMR_{GLC}$ 17 Networks, age group, HOMA-IR and cortical thickness.

| | Principal Component 1 | | | Principal Component 2 | | | Principal Component 3 | | | Principal Component 4 | | | Principal Component 5 | | |
|---|---|---|---|---|---|---|---|---|---|---|---|---|---|---|---|
| | F | p | $\eta^2_p$ | F | p | $\eta^2_p$ | F | p | $\eta^2_p$ | F | p | $\eta^2_p$ | F | p | $\eta^2_p$ |
| Visual Central | 1.6 | 0.206 | 0.031 | 0.9 | 0.358 | 0.016 | 6.3 | 0.015 | 0.108 | 0.5 | 0.498 | 0.009 | 1.3 | 0.256 | 0.025 |
| Visual Peripheral | 0.0 | 0.917 | 0.000 | 0.2 | 0.665 | 0.004 | 4.7 | 0.034 | 0.084 | 0.2 | 0.682 | 0.003 | 0.6 | 0.457 | 0.011 |
| Somatomotor A | 0.0 | 0.988 | 0.000 | 0.0 | 0.839 | 0.001 | 1.6 | 0.209 | 0.030 | 0.7 | 0.394 | 0.014 | 0.3 | 0.582 | 0.006 |
| Somatomotor B | 0.1 | 0.781 | 0.001 | 0.3 | 0.599 | 0.005 | 0.1 | 0.735 | 0.002 | 0.0 | 0.851 | 0.001 | 0.3 | 0.587 | 0.006 |
| Dors Attention A | 0.0 | 0.934 | 0.000 | 0.2 | 0.637 | 0.004 | 5.2 | 0.027 | 0.090 | 0.2 | 0.691 | 0.003 | 0.9 | 0.359 | 0.016 |
| Dors Attention B | 0.7 | 0.393 | 0.014 | 1.6 | 0.209 | 0.030 | 1.7 | 0.196 | 0.032 | 0.3 | 0.582 | 0.006 | 0.3 | 0.588 | 0.006 |
| Salience Ventral Attention A | 0.8 | 0.362 | 0.016 | 1.4 | 0.237 | 0.027 | 0.1 | 0.782 | 0.001 | 3.6 | 0.065 | 0.064 | 0.0 | 0.996 | 0.000 |
| Salience Ventral Attention B | 0.3 | 0.579 | 0.006 | 0.6 | 0.451 | 0.011 | 0.4 | 0.553 | 0.007 | 1.5 | 0.224 | 0.028 | 0.3 | 0.607 | 0.005 |
| Limbic A | 4.4 | 0.042 | 0.077 | 0.3 | 0.600 | 0.005 | 0.2 | 0.684 | 0.003 | 0.7 | 0.413 | 0.013 | 2.0 | 0.164 | 0.037 |
| Limbic B | 0.1 | 0.721 | 0.002 | 0.5 | 0.471 | 0.010 | 1.2 | 0.276 | 0.023 | 0.1 | 0.749 | 0.002 | 0.1 | 0.780 | 0.002 |
| Control A | 0.6 | 0.447 | 0.011 | 4.2 | 0.046 | 0.074 | 5.3 | 0.026 | 0.092 | 1.1 | 0.303 | 0.020 | 1.5 | 0.223 | 0.028 |
| Control B | 0.6 | 0.459 | 0.011 | 1.0 | 0.330 | 0.018 | 1.9 | 0.175 | 0.035 | 0.5 | 0.467 | 0.010 | 0.0 | 0.934 | 0.000 |
| Control C | 0.1 | 0.751 | 0.002 | 0.6 | 0.431 | 0.012 | 0.3 | 0.604 | 0.005 | 0.0 | 0.846 | 0.001 | 0.1 | 0.798 | 0.001 |
| Default A | 0.1 | 0.751 | 0.002 | 0.1 | 0.811 | 0.001 | 3.2 | 0.079 | 0.058 | 0.0 | 0.903 | 0.000 | 0.2 | 0.669 | 0.004 |
| Default B | 0.0 | 0.837 | 0.001 | 4.7 | 0.035 | 0.083 | 1.1 | 0.293 | 0.021 | 0.2 | 0.682 | 0.003 | 0.6 | 0.425 | 0.012 |
| Default C | 5.9 | 0.018 | 0.102 | 0.3 | 0.564 | 0.006 | 1.6 | 0.206 | 0.031 | 0.0 | 0.846 | 0.001 | 0.8 | 0.390 | 0.014 |
| Temporal Parietal | 1.0 | 0.313 | 0.020 | 0.5 | 0.493 | 0.009 | 0.7 | 0.398 | 0.014 | 2.8 | 0.101 | 0.051 | 0.0 | 0.912 | 0.000 |
| Subcortical | 4.7 | 0.035 | 0.083 | 0.9 | 0.358 | 0.016 | 0.6 | 0.427 | 0.012 | 0.1 | 0.776 | 0.002 | 0.1 | 0.774 | 0.002 |
| Cortical Thickness | 2.4 | 0.129 | 0.044 | 1.2 | 0.278 | 0.023 | 1.6 | 0.213 | 0.030 | 0.1 | 0.788 | 0.001 | 0.8 | 0.383 | 0.015 |
| Age Category | 9.5 | 0.003 | 0.154 | 5.5 | 0.023 | 0.095 | 0.3 | 0.601 | 0.005 | 1.3 | 0.264 | 0.024 | 0.7 | 0.421 | 0.012 |
| HOMA-IR | 0.2 | 0.664 | 0.004 | 0.0 | 0.944 | 0.000 | 0.5 | 0.470 | 0.010 | 4.9 | 0.031 | 0.086 | 0.0 | 0.942 | 0.000 |

#### *4. Effects of demographic variables on $CMR_{GLC}$ and cortical thickness*

We examined the effects of the demographic variables on the dependent variables by repeating the GLM models from the main analyses with the demographics included as covariates.

For the relationship of the variables with $CMR_{GLC}$, age, cortical thickness and HOMA-IR remained the predominant predictors across most regions when the other demographics were added to the GLM, although the relationships were somewhat attenuated or mediated by the other factors (see $\eta^2_{p,}$ Table S6).

Table S6. General linear models testing the association of regional $CMR_{GLC}$ with cortical thickness and other demographic variables.

| | Overall Model | | | Cortical Thickness | | | Age Group | | | HOMA-IR | | | Systolic BP | | | Diastolic BP | | | Resting Heart Rate | | | BMI | | | Sex | | | Years of Education | | |
|---|---|---|---|---|---|---|---|---|---|---|---|---|---|---|---|---|---|---|---|---|---|---|---|---|---|---|---|---|---|---|
| | F | p-FDR | $\eta^2_p$ | F | p | $\eta^2_p$ | F | p | $\eta^2_p$ | F | p | $\eta^2_p$ | F | p | $\eta^2_p$ | F | p | $\eta^2_p$ | F | p | $\eta^2_p$ | F | p | $\eta^2_p$ | F | p | $\eta^2_p$ | F | p | $\eta^2_p$ |
| L_Visual Central: Extra Striate Cortex 1 | 1.2 | 0.343 | 0.15 | 1.9 | 0.170 | 0.03 | 2.7 | 0.104 | 0.05 | 1.3 | 0.260 | 0.02 | 1.5 | 0.233 | 0.02 | 1.6 | 0.210 | 0.03 | 1.3 | 0.255 | 0.02 | 0.1 | 0.701 | 0.00 | 0.0 | 0.912 | 0.00 | 2.2 | 0.143 | 0.04 |
| L_Visual Central: Extra Striate Cortex 2 | 1.0 | 0.422 | 0.14 | 0.4 | 0.522 | 0.01 | 1.9 | 0.177 | 0.03 | 2.4 | 0.124 | 0.04 | 0.6 | 0.427 | 0.01 | 1.2 | 0.282 | 0.02 | 1.1 | 0.289 | 0.02 | 0.0 | 0.864 | 0.00 | 0.0 | 0.866 | 0.00 | 1.5 | 0.223 | 0.03 |
| L_Visual Central: Striate Cortex 1 | 1.3 | 0.275 | 0.17 | 0.5 | 0.474 | 0.01 | 3.1 | 0.081 | 0.05 | 2.2 | 0.146 | 0.04 | 0.9 | 0.349 | 0.02 | 1.3 | 0.259 | 0.02 | 2.6 | 0.115 | 0.04 | 0.0 | 0.917 | 0.00 | 0.0 | 0.935 | 0.00 | 0.8 | 0.366 | 0.01 |
| L_Visual Central: Extra Striate Cortex 3 | 1.6 | 0.156 | 0.20 | 2.3 | 0.134 | 0.04 | 4.3 | 0.042 | 0.07 | 2.5 | 0.118 | 0.04 | 1.4 | 0.239 | 0.02 | 1.9 | 0.175 | 0.03 | 1.0 | 0.325 | 0.02 | 0.2 | 0.669 | 0.00 | 0.0 | 0.903 | 0.00 | 1.2 | 0.278 | 0.02 |
| L_Visual Peripheral: Extra Striate Inferior 1 | 1.7 | 0.111 | 0.21 | 1.0 | 0.316 | 0.02 | 4.8 | 0.033 | 0.08 | 1.6 | 0.215 | 0.03 | 1.7 | 0.196 | 0.03 | 2.4 | 0.124 | 0.04 | 1.6 | 0.204 | 0.03 | 0.0 | 0.868 | 0.00 | 0.3 | 0.575 | 0.01 | 0.6 | 0.440 | 0.01 |
| L_Visual Peripheral: Striate Cortex Calcarine 1 | 2.4 | 0.027 | 0.27 | 6.4 | 0.014 | 0.10 | 2.5 | 0.120 | 0.04 | 3.4 | 0.071 | 0.06 | 1.3 | 0.260 | 0.02 | 1.8 | 0.181 | 0.03 | 1.7 | 0.203 | 0.03 | 0.0 | 0.975 | 0.00 | 0.2 | 0.665 | 0.00 | 1.1 | 0.297 | 0.02 |
| L_Visual Peripheral: Extra Striate Cortex Sup 1 | 2.4 | 0.027 | 0.27 | 1.0 | 0.311 | 0.02 | 4.1 | 0.046 | 0.07 | 3.8 | 0.055 | 0.06 | 0.5 | 0.467 | 0.01 | 2.1 | 0.153 | 0.03 | 1.6 | 0.217 | 0.03 | 0.1 | 0.737 | 0.00 | 0.4 | 0.520 | 0.01 | 0.2 | 0.662 | 0.00 |
| L_Somatomotor A: 1 | 4.3 | 0.001 | 0.40 | 11.9 | 0.001 | 0.17 | 6.3 | 0.015 | 0.10 | 5.3 | 0.025 | 0.08 | 3.3 | 0.076 | 0.05 | 5.6 | 0.021 | 0.09 | 2.5 | 0.122 | 0.04 | 0.3 | 0.610 | 0.00 | 0.2 | 0.683 | 0.00 | 1.3 | 0.258 | 0.02 |
| L_Somatomotor A: 2 | 3.7 | 0.003 | 0.36 | 8.0 | 0.006 | 0.12 | 3.7 | 0.060 | 0.06 | 4.6 | 0.036 | 0.07 | 3.8 | 0.055 | 0.06 | 5.5 | 0.023 | 0.09 | 1.7 | 0.200 | 0.03 | 0.3 | 0.619 | 0.00 | 0.7 | 0.404 | 0.01 | 1.1 | 0.303 | 0.02 |
| L_Somatomotor B: Auditory 1 | 3.4 | 0.004 | 0.35 | 4.9 | 0.031 | 0.08 | 2.8 | 0.099 | 0.05 | 3.8 | 0.055 | 0.06 | 1.2 | 0.283 | 0.02 | 2.3 | 0.138 | 0.04 | 0.3 | 0.560 | 0.01 | 0.3 | 0.607 | 0.00 | 0.0 | 0.825 | 0.00 | 1.7 | 0.191 | 0.03 |
| L_Somatomotor B: S2 1 | 2.8 | 0.010 | 0.31 | 0.2 | 0.650 | 0.00 | 5.3 | 0.025 | 0.08 | 2.8 | 0.102 | 0.05 | 1.1 | 0.294 | 0.02 | 3.0 | 0.090 | 0.05 | 1.1 | 0.294 | 0.02 | 0.1 | 0.753 | 0.00 | 0.4 | 0.517 | 0.01 | 0.9 | 0.348 | 0.02 |
| L_Somatomotor B: S2 2 | 4.8 | 0.001 | 0.42 | 3.0 | 0.089 | 0.05 | 6.3 | 0.015 | 0.10 | 1.8 | 0.187 | 0.03 | 0.8 | 0.369 | 0.01 | 2.0 | 0.161 | 0.03 | 0.0 | 0.833 | 0.00 | 0.7 | 0.409 | 0.01 | 1.2 | 0.270 | 0.02 | 1.0 | 0.314 | 0.02 |
| L_Somatomotor B: Central 1 | 2.4 | 0.026 | 0.27 | 3.7 | 0.059 | 0.06 | 4.8 | 0.032 | 0.08 | 3.3 | 0.073 | 0.05 | 1.5 | 0.230 | 0.02 | 1.7 | 0.193 | 0.03 | 0.4 | 0.520 | 0.01 | 0.0 | 0.871 | 0.00 | 0.0 | 0.838 | 0.00 | 0.9 | 0.343 | 0.02 |
| L_Dorsal Attention A: Temporal Occipital 1 | 3.1 | 0.006 | 0.33 | 9.2 | 0.004 | 0.14 | 0.8 | 0.388 | 0.01 | 1.9 | 0.172 | 0.03 | 0.0 | 0.866 | 0.00 | 1.7 | 0.202 | 0.03 | 1.9 | 0.176 | 0.03 | 0.0 | 0.895 | 0.00 | 0.0 | 0.831 | 0.00 | 1.4 | 0.240 | 0.02 |
| L_Dorsal Attention A: Parietal Occipital 1 | 2.6 | 0.020 | 0.28 | 3.3 | 0.074 | 0.05 | 5.3 | 0.025 | 0.08 | 1.8 | 0.180 | 0.03 | 1.8 | 0.180 | 0.03 | 1.5 | 0.220 | 0.03 | 0.1 | 0.736 | 0.00 | 0.7 | 0.413 | 0.01 | 0.1 | 0.773 | 0.00 | 2.0 | 0.163 | 0.03 |
| L_Dorsal Attention A: Superior Parietal Lobule 1 | 3.3 | 0.004 | 0.34 | 6.3 | 0.015 | 0.10 | 3.5 | 0.065 | 0.06 | 2.4 | 0.125 | 0.04 | 1.6 | 0.209 | 0.03 | 3.7 | 0.059 | 0.06 | 1.2 | 0.272 | 0.02 | 0.0 | 0.998 | 0.00 | 0.0 | 0.860 | 0.00 | 1.0 | 0.326 | 0.02 |
| L_Dorsal Attention B: Post Central 1 | 3.0 | 0.007 | 0.32 | 0.2 | 0.655 | 0.00 | 8.7 | 0.005 | 0.13 | 2.6 | 0.115 | 0.04 | 0.4 | 0.513 | 0.01 | 1.0 | 0.329 | 0.02 | 0.5 | 0.494 | 0.01 | 0.3 | 0.557 | 0.01 | 0.3 | 0.580 | 0.01 | 1.4 | 0.241 | 0.02 |
| L_Dorsal Attention B: Post Central 2 | 5.3 | 0.000 | 0.45 | 17.5 | 0.000 | 0.23 | 1.7 | 0.198 | 0.03 | 6.5 | 0.013 | 0.10 | 0.3 | 0.597 | 0.00 | 1.3 | 0.266 | 0.02 | 1.1 | 0.304 | 0.02 | 2.0 | 0.159 | 0.03 | 0.0 | 0.904 | 0.00 | 1.9 | 0.178 | 0.03 |
| L_Dorsal Attention B: Post Central 3 | 3.6 | 0.003 | 0.36 | 9.3 | 0.003 | 0.14 | 6.2 | 0.016 | 0.10 | 3.2 | 0.078 | 0.05 | 2.4 | 0.124 | 0.04 | 3.8 | 0.056 | 0.06 | 0.3 | 0.568 | 0.01 | 0.0 | 0.996 | 0.00 | 0.0 | 0.855 | 0.00 | 1.2 | 0.279 | 0.02 |
| L_Dorsal Attention B: Frontal Eye Fields 1 | 4.4 | 0.001 | 0.41 | 4.5 | 0.039 | 0.07 | 7.9 | 0.007 | 0.12 | 3.6 | 0.062 | 0.06 | 2.8 | 0.098 | 0.05 | 3.7 | 0.058 | 0.06 | 1.1 | 0.299 | 0.02 | 0.0 | 0.849 | 0.00 | 0.0 | 0.919 | 0.00 | 0.8 | 0.370 | 0.01 |
| L_Salience Ventral Attention A: Parietal Operculum 1 | 3.9 | 0.002 | 0.38 | 2.3 | 0.135 | 0.04 | 8.1 | 0.006 | 0.12 | 3.3 | 0.073 | 0.05 | 0.9 | 0.345 | 0.02 | 1.9 | 0.174 | 0.03 | 0.5 | 0.476 | 0.01 | 0.5 | 0.488 | 0.01 | 0.1 | 0.816 | 0.00 | 1.8 | 0.187 | 0.03 |
| L_Salience Ventral Attention A: Insula: 1 | 4.5 | 0.001 | 0.41 | 0.9 | 0.337 | 0.02 | 12.3 | 0.001 | 0.17 | 3.5 | 0.068 | 0.06 | 0.8 | 0.361 | 0.01 | 1.7 | 0.196 | 0.03 | 0.4 | 0.530 | 0.01 | 0.0 | 0.947 | 0.00 | 0.5 | 0.503 | 0.01 | 1.3 | 0.252 | 0.02 |
| L_Salience Ventral Attention A: Insula: 2 | 5.4 | 0.000 | 0.46 | 7.6 | 0.008 | 0.12 | 7.0 | 0.010 | 0.11 | 1.0 | 0.312 | 0.02 | 2.9 | 0.094 | 0.05 | 1.9 | 0.178 | 0.03 | 0.3 | 0.578 | 0.01 | 0.1 | 0.711 | 0.00 | 1.8 | 0.187 | 0.03 | 3.5 | 0.066 | 0.06 |
| L_Salience Ventral Attention A: Parietal Medial 1 | 5.3 | 0.000 | 0.45 | 14.6 | 0.000 | 0.20 | 0.9 | 0.340 | 0.02 | 4.4 | 0.040 | 0.07 | 1.7 | 0.196 | 0.03 | 3.9 | 0.055 | 0.06 | 1.0 | 0.322 | 0.02 | 0.5 | 0.468 | 0.01 | 1.6 | 0.216 | 0.03 | 0.5 | 0.497 | 0.01 |
| L_Salience Ventral Attention A: Frontal Medial 1 | 4.7 | 0.001 | 0.42 | 4.3 | 0.043 | 0.07 | 4.7 | 0.035 | 0.07 | 2.4 | 0.126 | 0.04 | 2.5 | 0.122 | 0.04 | 3.7 | 0.059 | 0.06 | 1.4 | 0.243 | 0.02 | 0.0 | 0.868 | 0.00 | 1.2 | 0.275 | 0.02 | 0.6 | 0.434 | 0.01 |
| L_Salience Ventral Attention B: Lateral Prefrontal Cortex 1 | 6.2 | 0.000 | 0.49 | 10.5 | 0.002 | 0.15 | 9.1 | 0.004 | 0.14 | 4.9 | 0.031 | 0.08 | 2.9 | 0.093 | 0.05 | 3.4 | 0.072 | 0.05 | 0.6 | 0.427 | 0.01 | 0.0 | 0.936 | 0.00 | 0.0 | 0.907 | 0.00 | 0.9 | 0.350 | 0.02 |
| L_Salience Ventral Attention B: Medial Posterior Prefrontal | 4.3 | 0.001 | 0.40 | 0.3 | 0.612 | 0.00 | 8.4 | 0.005 | 0.13 | 3.9 | 0.055 | 0.06 | 1.3 | 0.260 | 0.02 | 2.0 | 0.162 | 0.03 | 1.2 | 0.269 | 0.02 | 0.0 | 0.928 | 0.00 | 0.4 | 0.517 | 0.01 | 1.5 | 0.232 | 0.02 |
| L_Limbic A: Temporal Pole 1 | 2.8 | 0.012 | 0.30 | 2.0 | 0.162 | 0.03 | 6.5 | 0.014 | 0.10 | 2.0 | 0.161 | 0.03 | 0.5 | 0.462 | 0.01 | 1.5 | 0.222 | 0.03 | 0.5 | 0.496 | 0.01 | 0.0 | 0.870 | 0.00 | 2.1 | 0.157 | 0.03 | 0.9 | 0.353 | 0.01 |
| L_Limbic A: Temporal Pole 2 | 2.4 | 0.029 | 0.27 | 0.3 | 0.580 | 0.01 | 7.4 | 0.009 | 0.11 | 2.4 | 0.129 | 0.04 | 1.6 | 0.212 | 0.03 | 2.3 | 0.134 | 0.04 | 0.6 | 0.459 | 0.01 | 0.2 | 0.679 | 0.00 | 0.5 | 0.501 | 0.01 | 1.5 | 0.232 | 0.02 |
| L_Limbic B: Orbital Frontal Cortex 1 | 3.0 | 0.008 | 0.31 | 0.2 | 0.650 | 0.00 | 10.1 | 0.002 | 0.15 | 2.8 | 0.099 | 0.05 | 1.1 | 0.296 | 0.02 | 1.3 | 0.253 | 0.02 | 0.7 | 0.409 | 0.01 | 0.1 | 0.715 | 0.00 | 0.4 | 0.530 | 0.01 | 0.5 | 0.473 | 0.01 |
| L_Control A: Intraparietal Sulcus 1 | 4.4 | 0.001 | 0.41 | 6.9 | 0.011 | 0.11 | 4.7 | 0.033 | 0.08 | 3.0 | 0.089 | 0.05 | 1.9 | 0.171 | 0.03 | 4.8 | 0.033 | 0.08 | 1.0 | 0.332 | 0.02 | 0.3 | 0.568 | 0.01 | 0.0 | 0.950 | 0.00 | 2.8 | 0.103 | 0.05 |
| L_Control A: Lateral Prefrontal Cortex 1 | 4.0 | 0.002 | 0.38 | 3.3 | 0.076 | 0.05 | 7.1 | 0.010 | 0.11 | 1.6 | 0.218 | 0.03 | 2.2 | 0.148 | 0.04 | 2.2 | 0.141 | 0.04 | 0.5 | 0.476 | 0.01 | 0.4 | 0.552 | 0.01 | 1.0 | 0.325 | 0.02 | 1.9 | 0.168 | 0.03 |
| L_Control A: Lateral Prefrontal Cortex 2 | 4.7 | 0.001 | 0.42 | 9.9 | 0.003 | 0.15 | 2.7 | 0.103 | 0.05 | 0.7 | 0.395 | 0.01 | 2.5 | 0.116 | 0.04 | 1.5 | 0.225 | 0.03 | 0.1 | 0.822 | 0.00 | 0.2 | 0.638 | 0.00 | 1.9 | 0.170 | 0.03 | 2.0 | 0.160 | 0.03 |
| L_Control B: Lateral Prefrontal Cortex 1 | 4.0 | 0.002 | 0.38 | 5.4 | 0.024 | 0.08 | 9.9 | 0.003 | 0.15 | 1.6 | 0.208 | 0.03 | 2.6 | 0.111 | 0.04 | 2.4 | 0.127 | 0.04 | 0.7 | 0.391 | 0.01 | 0.7 | 0.423 | 0.01 | 0.2 | 0.669 | 0.00 | 1.0 | 0.319 | 0.02 |
| L_Control C: Precuneus 1 | 3.1 | 0.006 | 0.33 | 1.8 | 0.183 | 0.03 | 5.9 | 0.018 | 0.09 | 6.7 | 0.012 | 0.10 | 3.1 | 0.081 | 0.05 | 3.9 | 0.054 | 0.06 | 1.9 | 0.175 | 0.03 | 0.0 | 0.910 | 0.00 | 0.3 | 0.596 | 0.00 | 0.8 | 0.382 | 0.01 |
| L_Control C: Precuneus 2 | 3.5 | 0.003 | 0.35 | 5.1 | 0.028 | 0.08 | 4.3 | 0.043 | 0.07 | 5.1 | 0.027 | 0.08 | 1.2 | 0.272 | 0.02 | 2.0 | 0.164 | 0.03 | 0.7 | 0.412 | 0.01 | 0.2 | 0.631 | 0.00 | 1.5 | 0.230 | 0.02 | 1.4 | 0.241 | 0.02 |
| L_Control C: Cingulate Posterior 1 | 2.1 | 0.048 | 0.25 | 1.3 | 0.252 | 0.02 | 3.2 | 0.080 | 0.05 | 2.4 | 0.131 | 0.04 | 1.3 | 0.265 | 0.02 | 0.9 | 0.351 | 0.02 | 0.4 | 0.553 | 0.01 | 0.1 | 0.708 | 0.00 | 0.9 | 0.357 | 0.01 | 0.5 | 0.471 | 0.01 |
| L_Default A: Dorsal Prefrontal Cortex 1 | 6.8 | 0.000 | 0.51 | 9.7 | 0.003 | 0.14 | 11.9 | 0.001 | 0.17 | 2.1 | 0.149 | 0.04 | 3.2 | 0.079 | 0.05 | 3.3 | 0.075 | 0.05 | 0.7 | 0.422 | 0.01 | 0.2 | 0.673 | 0.00 | 0.4 | 0.532 | 0.01 | 1.6 | 0.216 | 0.03 |
| L_Default A: Precuneus Posterior Cingulate Cortex1 | 2.4 | 0.026 | 0.27 | 2.0 | 0.160 | 0.03 | 2.7 | 0.106 | 0.04 | 4.2 | 0.044 | 0.07 | 1.0 | 0.323 | 0.02 | 1.5 | 0.233 | 0.02 | 0.6 | 0.457 | 0.01 | 0.1 | 0.788 | 0.00 | 0.3 | 0.563 | 0.01 | 0.6 | 0.446 | 0.01 |
| L_Default A: Medial Prefrontal Cortex 1 | 4.8 | 0.001 | 0.43 | 1.1 | 0.309 | 0.02 | 13.4 | 0.001 | 0.19 | 2.5 | 0.117 | 0.04 | 1.0 | 0.320 | 0.02 | 1.7 | 0.197 | 0.03 | 0.7 | 0.394 | 0.01 | 0.2 | 0.667 | 0.00 | 0.5 | 0.498 | 0.01 | 1.1 | 0.307 | 0.02 |
| L_Default B: Temp 1 | 3.2 | 0.006 | 0.33 | 1.2 | 0.285 | 0.02 | 5.7 | 0.020 | 0.09 | 2.1 | 0.156 | 0.03 | 0.8 | 0.360 | 0.01 | 1.5 | 0.231 | 0.02 | 0.5 | 0.463 | 0.01 | 0.1 | 0.720 | 0.00 | 0.5 | 0.463 | 0.01 | 1.3 | 0.260 | 0.02 |
| L_Default B: Temp 2 | 4.0 | 0.002 | 0.38 | 9.6 | 0.003 | 0.14 | 2.7 | 0.103 | 0.05 | 2.0 | 0.159 | 0.03 | 0.8 | 0.390 | 0.01 | 2.2 | 0.143 | 0.04 | 0.9 | 0.352 | 0.01 | 0.3 | 0.597 | 0.00 | 0.2 | 0.641 | 0.00 | 2.3 | 0.139 | 0.04 |
| L_Default B: Inferior Parietal Lobule 1 | 3.8 | 0.002 | 0.37 | 6.4 | 0.014 | 0.10 | 2.2 | 0.144 | 0.04 | 3.5 | 0.066 | 0.06 | 2.0 | 0.159 | 0.03 | 2.7 | 0.105 | 0.04 | 0.4 | 0.521 | 0.01 | 0.3 | 0.581 | 0.01 | 0.1 | 0.707 | 0.00 | 1.5 | 0.231 | 0.02 |
| L_Default B: Dorsal Prefrontal Cortex 1 | 5.9 | 0.000 | 0.48 | 1.6 | 0.207 | 0.03 | 10.7 | 0.002 | 0.16 | 2.9 | 0.092 | 0.05 | 1.9 | 0.171 | 0.03 | 2.3 | 0.131 | 0.04 | 0.8 | 0.387 | 0.01 | 0.1 | 0.749 | 0.00 | 0.5 | 0.491 | 0.01 | 1.9 | 0.170 | 0.03 |
| L_Default B: Lateral Prefrontal Cortex 1 | 4.0 | 0.002 | 0.38 | 4.9 | 0.030 | 0.08 | 6.7 | 0.012 | 0.10 | 2.5 | 0.116 | 0.04 | 1.6 | 0.210 | 0.03 | 2.2 | 0.141 | 0.04 | 1.0 | 0.327 | 0.02 | 0.1 | 0.707 | 0.00 | 0.1 | 0.796 | 0.00 | 1.4 | 0.241 | 0.02 |
| L_Default B: Ventral Prefrontal Cortex 1 | 4.1 | 0.001 | 0.39 | 0.1 | 0.753 | 0.00 | 13.9 | 0.000 | 0.19 | 4.4 | 0.041 | 0.07 | 1.0 | 0.334 | 0.02 | 1.4 | 0.234 | 0.02 | 0.5 | 0.484 | 0.01 | 0.0 | 0.942 | 0.00 | 0.4 | 0.544 | 0.01 | 1.2 | 0.283 | 0.02 |
| L_Default B: Ventral Prefrontal Cortex 2 | 3.9 | 0.002 | 0.38 | 1.1 | 0.298 | 0.02 | 8.3 | 0.006 | 0.13 | 3.0 | 0.088 | 0.05 | 2.0 | 0.159 | 0.03 | 2.3 | 0.131 | 0.04 | 1.1 | 0.309 | 0.02 | 0.1 | 0.722 | 0.00 | 0.3 | 0.604 | 0.00 | 1.6 | 0.217 | 0.03 |
| L_Default C: Retro Superior Parietal Lobuleenial 1 | 2.9 | 0.009 | 0.31 | 0.6 | 0.440 | 0.01 | 6.8 | 0.012 | 0.10 | 5.2 | 0.026 | 0.08 | 1.2 | 0.283 | 0.02 | 1.7 | 0.195 | 0.03 | 0.2 | 0.697 | 0.00 | 0.0 | 0.915 | 0.00 | 0.9 | 0.336 | 0.02 | 1.4 | 0.235 | 0.02 |
| L_Default C: Parahippocampal Cortex 1 | 3.3 | 0.005 | 0.34 | 2.2 | 0.147 | 0.04 | 8.4 | 0.005 | 0.13 | 1.6 | 0.209 | 0.03 | 1.4 | 0.237 | 0.02 | 2.4 | 0.126 | 0.04 | 0.3 | 0.612 | 0.00 | 0.2 | 0.621 | 0.00 | 0.9 | 0.338 | 0.02 | 2.0 | 0.160 | 0.03 |
| L_Temporal Parietal 1 | 3.6 | 0.003 | 0.36 | 6.5 | 0.013 | 0.10 | 3.7 | 0.059 | 0.06 | 3.4 | 0.069 | 0.06 | 0.9 | 0.340 | 0.02 | 2.1 | 0.153 | 0.03 | 0.8 | 0.387 | 0.01 | 0.5 | 0.476 | 0.01 | 0.2 | 0.628 | 0.00 | 2.6 | 0.113 | 0.04 |

*continued …*

*… Table S6 continued*

| | Overall Model | | | Cortical Thickness | | | Age Group | | | HOMA-IR | | | Systolic BP | | | Diastolic BP | | | Resting Heart Rate | | | BMI | | | Sex | | | Years of Education | | |
|---|---|---|---|---|---|---|---|---|---|---|---|---|---|---|---|---|---|---|---|---|---|---|---|---|---|---|---|---|---|---|
| | F | p-FDR | $\eta^2_p$ | F | p | $\eta^2_p$ | F | p | $\eta^2_p$ | F | p | $\eta^2_p$ | F | p | $\eta^2_p$ | F | p | $\eta^2_p$ | F | p | $\eta^2_p$ | F | p | $\eta^2_p$ | F | p | $\eta^2_p$ | F | p | $\eta^2_p$ |
| R_Visual Central: Extra Striate Cortex 1 | 1.5 | 0.172 | 0.19 | 1.6 | 0.208 | 0.03 | 2.0 | 0.164 | 0.03 | 3.3 | 0.075 | 0.05 | 2.3 | 0.136 | 0.04 | 3.3 | 0.076 | 0.05 | 2.4 | 0.125 | 0.04 | 0.1 | 0.776 | 0.00 | 0.1 | 0.748 | 0.00 | 0.9 | 0.354 | 0.01 |
| R_Visual Central: Extra Striate Cortex 2 | 1.0 | 0.452 | 0.13 | 0.1 | 0.744 | 0.00 | 1.9 | 0.172 | 0.03 | 3.3 | 0.074 | 0.05 | 0.7 | 0.404 | 0.01 | 0.9 | 0.336 | 0.02 | 1.2 | 0.274 | 0.02 | 0.0 | 0.941 | 0.00 | 0.0 | 0.979 | 0.00 | 1.0 | 0.325 | 0.02 |
| R_Visual Central: Extra Striate Cortex 3 | 1.4 | 0.222 | 0.18 | 1.7 | 0.195 | 0.03 | 1.4 | 0.235 | 0.02 | 3.5 | 0.068 | 0.06 | 0.7 | 0.416 | 0.01 | 0.8 | 0.383 | 0.01 | 0.8 | 0.364 | 0.01 | 0.0 | 0.949 | 0.00 | 0.0 | 0.930 | 0.00 | 0.4 | 0.516 | 0.01 |
| R_Visual Peripheral: Striate Cortex Calcarine 1 | 1.2 | 0.341 | 0.15 | 0.1 | 0.787 | 0.00 | 2.8 | 0.100 | 0.05 | 1.8 | 0.187 | 0.03 | 0.7 | 0.406 | 0.01 | 1.4 | 0.249 | 0.02 | 2.3 | 0.134 | 0.04 | 0.0 | 0.896 | 0.00 | 0.0 | 1.000 | 0.00 | 0.3 | 0.594 | 0.00 |
| R_Visual Peripheral: Extra Striate Inferior 1 | 3.0 | 0.007 | 0.32 | 5.1 | 0.027 | 0.08 | 3.8 | 0.056 | 0.06 | 3.6 | 0.063 | 0.06 | 2.6 | 0.113 | 0.04 | 3.8 | 0.056 | 0.06 | 1.9 | 0.179 | 0.03 | 0.0 | 0.895 | 0.00 | 0.2 | 0.653 | 0.00 | 1.3 | 0.262 | 0.02 |
| R_Visual Peripheral: Extra Striate Superior 1 | 2.1 | 0.054 | 0.24 | 3.0 | 0.088 | 0.05 | 3.3 | 0.076 | 0.05 | 4.2 | 0.045 | 0.07 | 1.9 | 0.170 | 0.03 | 3.0 | 0.087 | 0.05 | 2.8 | 0.099 | 0.05 | 0.2 | 0.658 | 0.00 | 0.3 | 0.617 | 0.00 | 0.2 | 0.641 | 0.00 |
| R_Somatomotor A: 1 | 3.1 | 0.007 | 0.32 | 2.6 | 0.111 | 0.04 | 7.0 | 0.010 | 0.11 | 4.6 | 0.036 | 0.07 | 1.3 | 0.253 | 0.02 | 2.1 | 0.155 | 0.03 | 0.8 | 0.387 | 0.01 | 0.0 | 0.863 | 0.00 | 0.2 | 0.662 | 0.00 | 1.5 | 0.231 | 0.02 |
| R_Somatomotor A: 2 | 3.1 | 0.007 | 0.32 | 7.2 | 0.009 | 0.11 | 5.2 | 0.026 | 0.08 | 6.8 | 0.012 | 0.10 | 1.6 | 0.207 | 0.03 | 3.0 | 0.090 | 0.05 | 1.5 | 0.223 | 0.03 | 0.0 | 0.896 | 0.00 | 0.1 | 0.810 | 0.00 | 1.9 | 0.178 | 0.03 |
| R_Somatomotor A: 3 | 3.0 | 0.008 | 0.32 | 6.1 | 0.016 | 0.10 | 3.8 | 0.057 | 0.06 | 4.5 | 0.039 | 0.07 | 0.8 | 0.369 | 0.01 | 2.2 | 0.147 | 0.04 | 0.3 | 0.573 | 0.01 | 0.0 | 0.936 | 0.00 | 0.0 | 0.879 | 0.00 | 1.0 | 0.317 | 0.02 |
| R_Somatomotor A: 4 | 4.1 | 0.002 | 0.39 | 7.6 | 0.008 | 0.12 | 5.1 | 0.028 | 0.08 | 4.8 | 0.032 | 0.08 | 3.7 | 0.059 | 0.06 | 5.8 | 0.019 | 0.09 | 1.2 | 0.272 | 0.02 | 0.4 | 0.545 | 0.01 | 1.1 | 0.299 | 0.02 | 2.1 | 0.155 | 0.03 |
| R_Somatomotor B: Auditory 1 | 3.9 | 0.002 | 0.37 | 8.8 | 0.004 | 0.13 | 1.5 | 0.225 | 0.03 | 3.9 | 0.052 | 0.06 | 1.7 | 0.201 | 0.03 | 2.9 | 0.091 | 0.05 | 0.3 | 0.602 | 0.00 | 0.1 | 0.758 | 0.00 | 0.3 | 0.585 | 0.01 | 1.4 | 0.246 | 0.02 |
| R_Somatomotor B: S2 1 | 3.1 | 0.006 | 0.33 | 1.1 | 0.304 | 0.02 | 4.4 | 0.040 | 0.07 | 1.7 | 0.199 | 0.03 | 0.8 | 0.386 | 0.01 | 1.5 | 0.223 | 0.03 | 0.5 | 0.475 | 0.01 | 0.8 | 0.364 | 0.01 | 1.0 | 0.318 | 0.02 | 1.0 | 0.312 | 0.02 |
| R_Somatomotor B: S2 2 | 4.0 | 0.002 | 0.38 | 2.5 | 0.116 | 0.04 | 8.8 | 0.004 | 0.13 | 2.7 | 0.105 | 0.04 | 2.3 | 0.135 | 0.04 | 2.5 | 0.117 | 0.04 | 0.1 | 0.717 | 0.00 | 0.4 | 0.515 | 0.01 | 1.1 | 0.297 | 0.02 | 0.5 | 0.474 | 0.01 |
| R_Somatomotor B: Central 1 | 2.3 | 0.033 | 0.26 | 6.1 | 0.016 | 0.10 | 2.2 | 0.144 | 0.04 | 3.2 | 0.080 | 0.05 | 2.5 | 0.122 | 0.04 | 2.6 | 0.111 | 0.04 | 0.5 | 0.486 | 0.01 | 0.1 | 0.785 | 0.00 | 0.4 | 0.511 | 0.01 | 0.9 | 0.347 | 0.02 |
| R_Dorsal Attention A: Temporal Occipital 1 | 1.6 | 0.135 | 0.20 | 0.5 | 0.469 | 0.01 | 4.8 | 0.032 | 0.08 | 2.4 | 0.130 | 0.04 | 1.7 | 0.193 | 0.03 | 2.5 | 0.119 | 0.04 | 0.8 | 0.379 | 0.01 | 0.0 | 0.836 | 0.00 | 0.1 | 0.815 | 0.00 | 1.2 | 0.274 | 0.02 |
| R_Dorsal Attention A: Parietal Occipital 1 | 2.2 | 0.044 | 0.25 | 1.8 | 0.191 | 0.03 | 3.2 | 0.080 | 0.05 | 2.1 | 0.154 | 0.03 | 2.2 | 0.143 | 0.04 | 2.9 | 0.095 | 0.05 | 0.8 | 0.368 | 0.01 | 0.5 | 0.497 | 0.01 | 0.1 | 0.708 | 0.00 | 0.8 | 0.384 | 0.01 |
| R_Dorsal Attention A: Superior Parietal Lobule 1 | 3.6 | 0.003 | 0.36 | 6.4 | 0.014 | 0.10 | 5.9 | 0.019 | 0.09 | 5.2 | 0.026 | 0.08 | 1.8 | 0.185 | 0.03 | 3.9 | 0.054 | 0.06 | 1.1 | 0.309 | 0.02 | 0.0 | 0.887 | 0.00 | 0.0 | 0.856 | 0.00 | 1.3 | 0.266 | 0.02 |
| R_Dorsal Attention B: Post Central 1 | 3.2 | 0.006 | 0.33 | 2.5 | 0.116 | 0.04 | 1.8 | 0.184 | 0.03 | 4.2 | 0.045 | 0.07 | 0.6 | 0.435 | 0.01 | 2.2 | 0.147 | 0.04 | 0.9 | 0.354 | 0.01 | 0.0 | 0.864 | 0.00 | 0.3 | 0.561 | 0.01 | 0.2 | 0.686 | 0.00 |
| R_Dorsal Attention B: Post Central 2 | 3.6 | 0.003 | 0.36 | 10.7 | 0.002 | 0.16 | 0.9 | 0.343 | 0.02 | 3.1 | 0.082 | 0.05 | 0.6 | 0.442 | 0.01 | 2.4 | 0.130 | 0.04 | 0.8 | 0.366 | 0.01 | 0.1 | 0.766 | 0.00 | 0.2 | 0.690 | 0.00 | 0.5 | 0.470 | 0.01 |
| R_Dorsal Attention B: Frontal Eye Fields 1 | 6.6 | 0.000 | 0.51 | 16.9 | 0.000 | 0.23 | 8.6 | 0.005 | 0.13 | 4.1 | 0.047 | 0.07 | 6.0 | 0.017 | 0.09 | 5.9 | 0.018 | 0.09 | 2.5 | 0.116 | 0.04 | 0.1 | 0.719 | 0.00 | 0.1 | 0.812 | 0.00 | 1.1 | 0.289 | 0.02 |
| R_Salience Ventral Attention A: Parietal Operculum 1 | 4.0 | 0.002 | 0.38 | 0.7 | 0.419 | 0.01 | 8.2 | 0.006 | 0.12 | 3.6 | 0.061 | 0.06 | 0.9 | 0.343 | 0.02 | 1.8 | 0.180 | 0.03 | 0.2 | 0.674 | 0.00 | 0.7 | 0.413 | 0.01 | 0.1 | 0.776 | 0.00 | 0.4 | 0.510 | 0.01 |
| R_Salience Ventral Attention A: Insula: 1 | 3.6 | 0.003 | 0.36 | 0.1 | 0.715 | 0.00 | 8.0 | 0.006 | 0.12 | 2.7 | 0.108 | 0.04 | 1.5 | 0.219 | 0.03 | 2.5 | 0.118 | 0.04 | 0.7 | 0.420 | 0.01 | 0.1 | 0.784 | 0.00 | 1.0 | 0.312 | 0.02 | 0.9 | 0.344 | 0.02 |
| R_Salience Ventral Attention A: Parietal Medial 1 | 4.5 | 0.001 | 0.41 | 12.6 | 0.001 | 0.18 | 1.9 | 0.178 | 0.03 | 2.9 | 0.091 | 0.05 | 1.4 | 0.246 | 0.02 | 2.3 | 0.131 | 0.04 | 0.1 | 0.704 | 0.00 | 0.0 | 0.833 | 0.00 | 1.3 | 0.260 | 0.02 | 0.8 | 0.371 | 0.01 |
| R_Salience Ventral Attention A: Frontal Medial 1 | 4.9 | 0.001 | 0.43 | 5.7 | 0.021 | 0.09 | 2.0 | 0.161 | 0.03 | 2.1 | 0.152 | 0.04 | 0.8 | 0.390 | 0.01 | 2.2 | 0.148 | 0.04 | 1.1 | 0.303 | 0.02 | 0.3 | 0.561 | 0.01 | 1.4 | 0.238 | 0.02 | 1.0 | 0.330 | 0.02 |
| R_Salience Ventral Attention B: Inferior Parietal Lobule 1 | 3.4 | 0.004 | 0.34 | 1.0 | 0.331 | 0.02 | 7.4 | 0.009 | 0.11 | 2.2 | 0.143 | 0.04 | 0.5 | 0.474 | 0.01 | 1.0 | 0.317 | 0.02 | 0.2 | 0.649 | 0.00 | 0.5 | 0.482 | 0.01 | 0.3 | 0.619 | 0.00 | 0.8 | 0.383 | 0.01 |
| R_Salience Ventral Attention B: Lateral Prefrontal Cortex 1 | 4.4 | 0.001 | 0.41 | 2.4 | 0.130 | 0.04 | 8.9 | 0.004 | 0.13 | 2.4 | 0.124 | 0.04 | 2.0 | 0.161 | 0.03 | 2.9 | 0.096 | 0.05 | 0.8 | 0.384 | 0.01 | 0.2 | 0.665 | 0.00 | 0.2 | 0.638 | 0.00 | 1.1 | 0.304 | 0.02 |
| R_Salience Ventral Attention B: Medial Posterior Prefrontal | 5.8 | 0.000 | 0.47 | 0.5 | 0.494 | 0.01 | 13.5 | 0.001 | 0.19 | 2.4 | 0.123 | 0.04 | 1.7 | 0.193 | 0.03 | 2.4 | 0.125 | 0.04 | 1.6 | 0.205 | 0.03 | 0.2 | 0.647 | 0.00 | 1.6 | 0.205 | 0.03 | 1.5 | 0.233 | 0.02 |
| R_Limbic A: Temporal Pole 1 | 3.1 | 0.007 | 0.32 | 2.0 | 0.158 | 0.03 | 7.1 | 0.010 | 0.11 | 2.4 | 0.129 | 0.04 | 0.9 | 0.339 | 0.02 | 2.0 | 0.159 | 0.03 | 0.7 | 0.391 | 0.01 | 0.1 | 0.796 | 0.00 | 1.4 | 0.247 | 0.02 | 0.5 | 0.500 | 0.01 |
| R_Limbic B: Orbital Frontal Cortex 1 | 2.9 | 0.010 | 0.31 | 0.0 | 0.917 | 0.00 | 11.4 | 0.001 | 0.16 | 2.2 | 0.142 | 0.04 | 1.7 | 0.197 | 0.03 | 2.2 | 0.147 | 0.04 | 1.0 | 0.333 | 0.02 | 0.2 | 0.633 | 0.00 | 0.3 | 0.589 | 0.01 | 0.7 | 0.406 | 0.01 |
| R_Control A: Intraparietal Sulcus 1 | 2.6 | 0.020 | 0.28 | 4.1 | 0.047 | 0.07 | 2.8 | 0.102 | 0.05 | 1.9 | 0.178 | 0.03 | 1.0 | 0.325 | 0.02 | 2.0 | 0.166 | 0.03 | 0.1 | 0.769 | 0.00 | 0.0 | 0.893 | 0.00 | 0.1 | 0.749 | 0.00 | 2.1 | 0.156 | 0.03 |
| R_Control A: Lateral Prefrontal Cortex 1 | 3.4 | 0.004 | 0.35 | 0.0 | 0.921 | 0.00 | 12.1 | 0.001 | 0.17 | 2.6 | 0.115 | 0.04 | 1.1 | 0.297 | 0.02 | 2.0 | 0.160 | 0.03 | 0.3 | 0.581 | 0.01 | 0.3 | 0.575 | 0.01 | 0.4 | 0.553 | 0.01 | 1.0 | 0.310 | 0.02 |
| R_Control A: Lateral Prefrontal Cortex 2 | 3.5 | 0.003 | 0.35 | 2.1 | 0.155 | 0.03 | 8.7 | 0.005 | 0.13 | 2.4 | 0.126 | 0.04 | 2.6 | 0.110 | 0.04 | 2.7 | 0.104 | 0.04 | 0.4 | 0.535 | 0.01 | 0.2 | 0.621 | 0.00 | 0.2 | 0.628 | 0.00 | 1.1 | 0.305 | 0.02 |
| R_Control B: Temporal 1 | 2.9 | 0.010 | 0.31 | 0.1 | 0.814 | 0.00 | 10.0 | 0.003 | 0.15 | 3.4 | 0.069 | 0.06 | 1.8 | 0.181 | 0.03 | 2.3 | 0.135 | 0.04 | 0.6 | 0.433 | 0.01 | 0.1 | 0.714 | 0.00 | 0.6 | 0.447 | 0.01 | 0.7 | 0.401 | 0.01 |
| R_Control B: inferior parietal lobule 1 | 4.3 | 0.001 | 0.40 | 8.0 | 0.006 | 0.12 | 4.8 | 0.032 | 0.08 | 3.0 | 0.090 | 0.05 | 1.9 | 0.179 | 0.03 | 3.0 | 0.091 | 0.05 | 2.0 | 0.161 | 0.03 | 0.5 | 0.461 | 0.01 | 0.1 | 0.702 | 0.00 | 1.0 | 0.315 | 0.02 |
| R_Control B: Lateral Prefrontal Cortexd 1 | 4.6 | 0.001 | 0.42 | 5.6 | 0.021 | 0.09 | 4.8 | 0.032 | 0.08 | 3.5 | 0.066 | 0.06 | 1.2 | 0.269 | 0.02 | 2.3 | 0.138 | 0.04 | 0.6 | 0.436 | 0.01 | 0.3 | 0.576 | 0.01 | 0.1 | 0.722 | 0.00 | 1.7 | 0.192 | 0.03 |
| R_Control B: Lateral Prefrontal Cortexv 1 | 3.6 | 0.003 | 0.36 | 1.3 | 0.263 | 0.02 | 8.5 | 0.005 | 0.13 | 3.5 | 0.066 | 0.06 | 1.6 | 0.218 | 0.03 | 1.8 | 0.183 | 0.03 | 1.0 | 0.319 | 0.02 | 0.1 | 0.737 | 0.00 | 0.3 | 0.615 | 0.00 | 0.2 | 0.680 | 0.00 |
| R_Control C: Cingulate Posterior 1 | 3.2 | 0.006 | 0.33 | 2.1 | 0.148 | 0.04 | 6.8 | 0.012 | 0.10 | 3.6 | 0.064 | 0.06 | 1.9 | 0.175 | 0.03 | 2.4 | 0.125 | 0.04 | 0.9 | 0.340 | 0.02 | 0.0 | 0.840 | 0.00 | 0.7 | 0.398 | 0.01 | 0.9 | 0.336 | 0.02 |
| R_Control C: Precuneus 1 | 2.7 | 0.014 | 0.30 | 1.7 | 0.197 | 0.03 | 4.8 | 0.032 | 0.08 | 3.9 | 0.053 | 0.06 | 2.5 | 0.118 | 0.04 | 3.5 | 0.066 | 0.06 | 1.1 | 0.293 | 0.02 | 0.0 | 0.869 | 0.00 | 0.6 | 0.427 | 0.01 | 0.2 | 0.677 | 0.00 |
| R_Default A: Inferior Parietal Lobule 1 | 3.7 | 0.003 | 0.36 | 6.8 | 0.011 | 0.11 | 1.4 | 0.246 | 0.02 | 3.2 | 0.077 | 0.05 | 1.6 | 0.204 | 0.03 | 2.1 | 0.149 | 0.04 | 1.3 | 0.261 | 0.02 | 0.5 | 0.466 | 0.01 | 0.4 | 0.547 | 0.01 | 0.6 | 0.448 | 0.01 |
| R_Default A: Dorsal Prefrontal Cortex 1 | 5.1 | 0.000 | 0.44 | 5.7 | 0.021 | 0.09 | 9.8 | 0.003 | 0.14 | 2.6 | 0.114 | 0.04 | 3.3 | 0.076 | 0.05 | 3.8 | 0.055 | 0.06 | 2.9 | 0.095 | 0.05 | 0.1 | 0.708 | 0.00 | 0.1 | 0.757 | 0.00 | 1.0 | 0.333 | 0.02 |
| R_Default A: Precuneus Posterior Cingulate Cortex 1 | 2.1 | 0.047 | 0.25 | 0.4 | 0.531 | 0.01 | 4.8 | 0.033 | 0.08 | 3.8 | 0.056 | 0.06 | 1.6 | 0.209 | 0.03 | 2.6 | 0.114 | 0.04 | 0.7 | 0.418 | 0.01 | 0.0 | 0.878 | 0.00 | 0.3 | 0.575 | 0.01 | 0.4 | 0.535 | 0.01 |
| R_Default A: Medial Prefrontal Cortex 1 | 5.0 | 0.000 | 0.44 | 0.1 | 0.805 | 0.00 | 14.5 | 0.000 | 0.20 | 3.2 | 0.080 | 0.05 | 0.7 | 0.412 | 0.01 | 1.2 | 0.272 | 0.02 | 0.8 | 0.376 | 0.01 | 0.4 | 0.524 | 0.01 | 0.4 | 0.523 | 0.01 | 1.4 | 0.245 | 0.02 |
| R_Default B: Dorsal Prefrontal Cortex 1 | 6.2 | 0.000 | 0.49 | 4.0 | 0.049 | 0.07 | 9.8 | 0.003 | 0.14 | 2.9 | 0.096 | 0.05 | 2.7 | 0.106 | 0.04 | 3.3 | 0.074 | 0.05 | 0.8 | 0.388 | 0.01 | 0.2 | 0.645 | 0.00 | 0.5 | 0.483 | 0.01 | 1.3 | 0.255 | 0.02 |
| R_Default B: Ventral Prefrontal Cortex 1 | 3.6 | 0.003 | 0.36 | 1.4 | 0.243 | 0.02 | 8.9 | 0.004 | 0.13 | 3.8 | 0.056 | 0.06 | 1.8 | 0.186 | 0.03 | 2.7 | 0.108 | 0.04 | 0.7 | 0.401 | 0.01 | 0.0 | 0.824 | 0.00 | 0.3 | 0.560 | 0.01 | 0.1 | 0.742 | 0.00 |
| R_Default B: Ventral Prefrontal Cortex 2 | 4.1 | 0.002 | 0.39 | 1.4 | 0.249 | 0.02 | 11.5 | 0.001 | 0.16 | 2.6 | 0.113 | 0.04 | 2.3 | 0.137 | 0.04 | 1.9 | 0.170 | 0.03 | 0.6 | 0.444 | 0.01 | 0.0 | 0.885 | 0.00 | 0.8 | 0.373 | 0.01 | 0.9 | 0.349 | 0.02 |
| R_Default C: Retro Superior Parietal Lobuleenial 1 | 2.4 | 0.027 | 0.27 | 1.9 | 0.171 | 0.03 | 3.1 | 0.085 | 0.05 | 3.0 | 0.088 | 0.05 | 1.5 | 0.223 | 0.03 | 2.5 | 0.118 | 0.04 | 1.0 | 0.311 | 0.02 | 0.0 | 0.886 | 0.00 | 0.9 | 0.336 | 0.02 | 0.1 | 0.745 | 0.00 |
| R_Default C: Parahippocampal Cortex 1 | 2.4 | 0.028 | 0.27 | 2.0 | 0.167 | 0.03 | 3.2 | 0.080 | 0.05 | 2.3 | 0.134 | 0.04 | 0.8 | 0.365 | 0.01 | 2.3 | 0.133 | 0.04 | 1.3 | 0.254 | 0.02 | 0.0 | 0.847 | 0.00 | 0.7 | 0.391 | 0.01 | 0.6 | 0.449 | 0.01 |
| R_Temporal Parietal 1 | 3.6 | 0.003 | 0.36 | 3.9 | 0.054 | 0.06 | 7.1 | 0.010 | 0.11 | 1.9 | 0.172 | 0.03 | 1.2 | 0.279 | 0.02 | 2.0 | 0.165 | 0.03 | 0.1 | 0.708 | 0.00 | 0.1 | 0.803 | 0.00 | 0.7 | 0.404 | 0.01 | 0.5 | 0.486 | 0.01 |
| R_Temporal Parietal 2 | 3.6 | 0.003 | 0.36 | 2.8 | 0.100 | 0.05 | 3.2 | 0.080 | 0.05 | 4.2 | 0.046 | 0.07 | 0.9 | 0.335 | 0.02 | 2.3 | 0.135 | 0.04 | 0.9 | 0.345 | 0.02 | 0.2 | 0.666 | 0.00 | 0.1 | 0.753 | 0.00 | 0.6 | 0.439 | 0.01 |
| R_Temporal Parietal 3 | 2.4 | 0.027 | 0.27 | 3.0 | 0.089 | 0.05 | 2.5 | 0.116 | 0.04 | 1.4 | 0.244 | 0.02 | 1.1 | 0.307 | 0.02 | 1.5 | 0.232 | 0.02 | 0.4 | 0.537 | 0.01 | 0.3 | 0.581 | 0.01 | 0.3 | 0.599 | 0.00 | 0.8 | 0.375 | 0.01 |

### Table S7. General linear models of age group, HOMA-IR2 and age group x HOMA-IR2 effect on regional $CMR_{GLC}$, including cortical thickness as a covariate.

**Left Hemisphere**

| | Overall Model | | | Cortial Thickness | | | Age Category | | | HOMA-IR2 | | | Age Group x HOMA-IR2 | | |
|---|---|---|---|---|---|---|---|---|---|---|---|---|---|---|---|
| | F | p-FDR | $\eta^2_p$ | F | p | $\eta^2_p$ | F | p | $\eta^2_p$ | F | p | $\eta^2_p$ | F | p | $\eta^2_p$ |
| Visual Central: Extra Striate Cortex 1 | 2.5 | 0.053 | 0.117 | 1.1 | 0.299 | 0.015 | 4.4 | 0.039 | 0.056 | 3.7 | 0.057 | 0.048 | 2.4 | 0.127 | 0.031 |
| Visual Central: Extra Striate Cortex 2 | 3.0 | 0.024 | 0.139 | 0.2 | 0.649 | 0.003 | 5.7 | 0.020 | 0.071 | 6.8 | 0.011 | 0.084 | 3.6 | 0.062 | 0.046 |
| Visual Central: Striate Cortex 1 | 3.0 | 0.023 | 0.141 | 0.2 | 0.624 | 0.003 | 6.6 | 0.012 | 0.082 | 5.4 | 0.023 | 0.068 | 2.6 | 0.110 | 0.034 |
| Visual Central: Extra Striate Cortex 3 | 4.9 | 0.002 | 0.209 | 1.2 | 0.283 | 0.016 | 10.5 | 0.002 | 0.124 | 5.5 | 0.022 | 0.069 | 6.2 | 0.015 | 0.077 |
| Visual Peripheral: Extra Striate Inferior 1 | 4.5 | 0.003 | 0.195 | 0.5 | 0.497 | 0.006 | 9.7 | 0.003 | 0.116 | 4.1 | 0.046 | 0.053 | 3.5 | 0.067 | 0.045 |
| Visual Peripheral: Striate Cortex Calcarine 1 | 7.3 | 0.000 | 0.283 | 9.2 | 0.003 | 0.111 | 9.6 | 0.003 | 0.115 | 7.9 | 0.006 | 0.097 | 5.6 | 0.020 | 0.071 |
| Visual Peripheral: Extra Striate CortexSup 1 | 7.2 | 0.000 | 0.281 | 2.2 | 0.147 | 0.028 | 14.9 | 0.000 | 0.168 | 8.0 | 0.006 | 0.097 | 6.4 | 0.013 | 0.080 |
| Somatomotor A: 1 | 10.1 | 0.000 | 0.353 | 7.2 | 0.009 | 0.089 | 13.8 | 0.000 | 0.158 | 7.1 | 0.010 | 0.087 | 4.7 | 0.034 | 0.059 |
| Somatomotor A: 2 | 8.7 | 0.000 | 0.320 | 4.9 | 0.029 | 0.063 | 8.4 | 0.005 | 0.102 | 8.4 | 0.005 | 0.102 | 3.6 | 0.060 | 0.047 |
| Somatomotor B: Auditory 1 | 9.7 | 0.000 | 0.343 | 5.6 | 0.020 | 0.071 | 11.5 | 0.001 | 0.134 | 10.8 | 0.002 | 0.127 | 6.2 | 0.015 | 0.077 |
| Somatomotor B: S2 1 | 7.3 | 0.000 | 0.283 | 1.5 | 0.229 | 0.020 | 13.7 | 0.000 | 0.156 | 7.8 | 0.007 | 0.096 | 4.5 | 0.038 | 0.057 |
| Somatomotor B: S2 2 | 13.6 | 0.000 | 0.423 | 3.5 | 0.064 | 0.045 | 22.4 | 0.000 | 0.232 | 8.9 | 0.004 | 0.107 | 6.4 | 0.014 | 0.079 |
| Somatomotor B: Central 1 | 6.8 | 0.000 | 0.269 | 2.6 | 0.113 | 0.034 | 12.3 | 0.001 | 0.142 | 7.3 | 0.009 | 0.089 | 4.6 | 0.035 | 0.059 |
| Dorsal Attention A: Temporal Occipital 1 | 9.7 | 0.000 | 0.344 | 13.6 | 0.000 | 0.155 | 14.2 | 0.000 | 0.161 | 5.3 | 0.025 | 0.066 | 9.0 | 0.004 | 0.109 |
| Dorsal Attention A: Parietal Occipital 1 | 5.9 | 0.000 | 0.242 | 1.0 | 0.328 | 0.013 | 9.8 | 0.003 | 0.117 | 7.1 | 0.009 | 0.088 | 4.4 | 0.040 | 0.056 |
| Dorsal Attention A: Superior Parietal Lobule 1 | 8.1 | 0.000 | 0.306 | 4.8 | 0.032 | 0.061 | 10.6 | 0.002 | 0.125 | 7.1 | 0.010 | 0.087 | 5.1 | 0.026 | 0.065 |
| Dorsal Attention B: Post Central 1 | 7.8 | 0.000 | 0.296 | 0.1 | 0.732 | 0.002 | 14.6 | 0.000 | 0.165 | 7.1 | 0.009 | 0.088 | 2.7 | 0.104 | 0.035 |
| Dorsal Attention B: Post Central 2 | 12.7 | 0.000 | 0.407 | 15.9 | 0.000 | 0.177 | 6.2 | 0.015 | 0.077 | 7.8 | 0.007 | 0.095 | 2.8 | 0.097 | 0.037 |
| Dorsal Attention B: Post Central 3 | 11.7 | 0.000 | 0.388 | 12.0 | 0.001 | 0.140 | 16.0 | 0.000 | 0.178 | 8.0 | 0.006 | 0.097 | 7.5 | 0.008 | 0.092 |
| Dorsal Attention B: Frontal Eye Fields 1 | 11.6 | 0.000 | 0.386 | 3.3 | 0.074 | 0.042 | 16.2 | 0.000 | 0.180 | 6.8 | 0.011 | 0.084 | 4.1 | 0.046 | 0.053 |
| Salience Ventral Attention A: Parietal Operculum 1 | 10.1 | 0.000 | 0.354 | 1.4 | 0.233 | 0.019 | 16.7 | 0.000 | 0.184 | 10.1 | 0.002 | 0.120 | 4.5 | 0.037 | 0.057 |
| Salience Ventral Attention A: Insula: 1 | 12.9 | 0.000 | 0.411 | 1.1 | 0.307 | 0.014 | 22.1 | 0.000 | 0.230 | 8.6 | 0.005 | 0.104 | 4.5 | 0.036 | 0.058 |
| Salience Ventral Attention A: Insula: 2 | 12.2 | 0.000 | 0.398 | 5.3 | 0.025 | 0.066 | 12.8 | 0.001 | 0.147 | 6.4 | 0.013 | 0.080 | 3.6 | 0.062 | 0.046 |
| Salience Ventral Attention A: Parietal Medial 1 | 12.7 | 0.000 | 0.408 | 13.2 | 0.001 | 0.152 | 7.6 | 0.007 | 0.093 | 13.0 | 0.001 | 0.149 | 3.9 | 0.052 | 0.050 |
| Salience Ventral Attention A: Frontal Medial 1 | 12.1 | 0.000 | 0.395 | 7.7 | 0.007 | 0.094 | 10.1 | 0.002 | 0.120 | 6.8 | 0.011 | 0.084 | 4.4 | 0.038 | 0.057 |
| Salience Ventral Attention B: Lateral Prefrontal Cortex | 16.6 | 0.000 | 0.472 | 14.0 | 0.000 | 0.159 | 14.1 | 0.000 | 0.160 | 10.8 | 0.002 | 0.127 | 3.8 | 0.056 | 0.048 |
| Salience Ventral Attention B: Medial Posterior Prefront | 11.0 | 0.000 | 0.372 | 1.8 | 0.188 | 0.023 | 11.7 | 0.001 | 0.137 | 8.9 | 0.004 | 0.107 | 1.7 | 0.197 | 0.022 |
| Limbic A Temporal Pole 1 | 8.2 | 0.000 | 0.308 | 0.1 | 0.703 | 0.002 | 15.5 | 0.000 | 0.173 | 9.7 | 0.003 | 0.116 | 2.4 | 0.125 | 0.032 |
| Limbic A: Temporal Pole 2 | 7.8 | 0.000 | 0.296 | 3.6 | 0.063 | 0.046 | 15.4 | 0.000 | 0.172 | 6.6 | 0.012 | 0.082 | 3.9 | 0.053 | 0.050 |
| Limbic B: Orbital Frontal Cortex 1 | 6.2 | 0.000 | 0.251 | 0.4 | 0.519 | 0.006 | 12.9 | 0.001 | 0.149 | 8.3 | 0.005 | 0.101 | 3.3 | 0.071 | 0.043 |
| Control A: Intraparietal Sulcus 1 | 10.5 | 0.000 | 0.362 | 6.1 | 0.016 | 0.076 | 12.7 | 0.001 | 0.147 | 7.1 | 0.009 | 0.088 | 4.7 | 0.033 | 0.060 |
| Control A: Lateral Prefrontal Cortex 1 | 9.8 | 0.000 | 0.346 | 3.2 | 0.077 | 0.042 | 11.6 | 0.001 | 0.136 | 6.5 | 0.013 | 0.081 | 3.0 | 0.089 | 0.039 |
| Control A: Lateral Prefrontal Cortex 2 | 10.4 | 0.000 | 0.361 | 6.6 | 0.012 | 0.082 | 4.9 | 0.029 | 0.062 | 5.1 | 0.027 | 0.064 | 1.5 | 0.230 | 0.019 |
| Control B: Lateral Prefrontal Cortexv 1 | 10.4 | 0.000 | 0.359 | 5.8 | 0.018 | 0.073 | 15.3 | 0.000 | 0.171 | 8.0 | 0.006 | 0.098 | 3.8 | 0.055 | 0.049 |
| Control C: Precuneus 1 | 6.2 | 0.000 | 0.250 | 0.8 | 0.383 | 0.010 | 7.1 | 0.009 | 0.088 | 11.5 | 0.001 | 0.135 | 2.1 | 0.148 | 0.028 |
| Control C: Precuneus 2 | 8.1 | 0.000 | 0.305 | 3.7 | 0.060 | 0.047 | 11.1 | 0.001 | 0.130 | 11.8 | 0.001 | 0.138 | 4.6 | 0.035 | 0.058 |
| Control C: Cingulate Posterior 1 | 6.2 | 0.000 | 0.251 | 1.8 | 0.178 | 0.024 | 8.6 | 0.004 | 0.105 | 8.9 | 0.004 | 0.107 | 3.6 | 0.063 | 0.046 |
| Default A: Dorsal Prefrontal Cortex 1 | 16.6 | 0.000 | 0.473 | 10.5 | 0.002 | 0.125 | 15.0 | 0.000 | 0.168 | 8.0 | 0.006 | 0.098 | 2.8 | 0.096 | 0.037 |
| Default A: Precuneus Posterior Cingulate Cortex1 | 6.7 | 0.000 | 0.267 | 2.8 | 0.099 | 0.036 | 8.1 | 0.006 | 0.098 | 11.1 | 0.001 | 0.131 | 3.2 | 0.078 | 0.041 |
| Default A: Medial Prefrontal Cortex 1 | 12.5 | 0.000 | 0.403 | 1.3 | 0.253 | 0.018 | 19.9 | 0.000 | 0.212 | 8.3 | 0.005 | 0.101 | 2.8 | 0.100 | 0.036 |
| Default B: Temp 1 | 9.4 | 0.000 | 0.337 | 3.1 | 0.084 | 0.040 | 15.3 | 0.000 | 0.171 | 7.3 | 0.009 | 0.089 | 5.0 | 0.028 | 0.064 |
| Default B: Temp 2 | 9.5 | 0.000 | 0.338 | 7.7 | 0.007 | 0.094 | 10.0 | 0.002 | 0.119 | 6.5 | 0.013 | 0.081 | 4.2 | 0.043 | 0.054 |
| Default B: Inferior Parietal Lobule 1 | 9.2 | 0.000 | 0.333 | 4.8 | 0.032 | 0.061 | 8.8 | 0.004 | 0.106 | 10.4 | 0.002 | 0.123 | 5.2 | 0.025 | 0.066 |
| Default B: Dorsal Prefrontal Cortex 1 | 15.3 | 0.000 | 0.453 | 3.9 | 0.052 | 0.050 | 14.9 | 0.000 | 0.168 | 8.3 | 0.005 | 0.100 | 3.1 | 0.082 | 0.040 |
| Default B: Lateral Prefrontal Cortex 1 | 10.8 | 0.000 | 0.370 | 5.8 | 0.018 | 0.073 | 12.5 | 0.001 | 0.144 | 6.6 | 0.012 | 0.081 | 3.3 | 0.075 | 0.042 |
| Default B: Ventral Prefrontal Cortex 1 | 11.4 | 0.000 | 0.381 | 0.0 | 0.989 | 0.000 | 19.5 | 0.000 | 0.208 | 10.1 | 0.002 | 0.120 | 2.2 | 0.143 | 0.029 |
| Default B: Ventral Prefrontal Cortex 2 | 10.3 | 0.000 | 0.358 | 2.5 | 0.117 | 0.033 | 14.6 | 0.000 | 0.165 | 9.7 | 0.003 | 0.115 | 4.1 | 0.046 | 0.053 |
| Default C: RetroSuperior Parietal Lobuleenial 1 | 7.4 | 0.000 | 0.286 | 0.6 | 0.430 | 0.008 | 10.3 | 0.002 | 0.122 | 13.1 | 0.001 | 0.150 | 2.0 | 0.164 | 0.026 |
| Default C: Parahippocampal Cortex 1 | 7.3 | 0.000 | 0.284 | 2.2 | 0.144 | 0.029 | 12.4 | 0.001 | 0.143 | 5.9 | 0.018 | 0.074 | 2.9 | 0.095 | 0.037 |
| Temporal Parietal 1 | 10.1 | 0.000 | 0.352 | 5.1 | 0.026 | 0.065 | 15.4 | 0.000 | 0.172 | 10.6 | 0.002 | 0.125 | 6.9 | 0.010 | 0.086 |
| Caudate | 5.8 | 0.000 | 0.238 | 2.8 | 0.098 | 0.037 | 6.6 | 0.012 | 0.081 | 5.8 | 0.019 | 0.073 | 2.8 | 0.100 | 0.036 |
| Putamen | 5.7 | 0.001 | 0.237 | 1.4 | 0.237 | 0.019 | 7.2 | 0.009 | 0.089 | 3.8 | 0.054 | 0.049 | 1.5 | 0.224 | 0.020 |
| Pallidum | 5.7 | 0.001 | 0.235 | 0.8 | 0.362 | 0.011 | 5.8 | 0.018 | 0.073 | 5.4 | 0.023 | 0.068 | 0.6 | 0.457 | 0.007 |
| Thalamus | 5.6 | 0.001 | 0.231 | 1.4 | 0.238 | 0.019 | 4.0 | 0.048 | 0.052 | 9.2 | 0.003 | 0.111 | 0.6 | 0.438 | 0.008 |

**Right Hemisphere**

| | Overall Model | | | Cortial Thickness | | | Age Category | | | HOMA-IR2 | | | Age Group x HOMA-IR2 | | |
|---|---|---|---|---|---|---|---|---|---|---|---|---|---|---|---|
| | F | p-FDR | $\eta^2_p$ | F | p | $\eta^2_p$ | F | p | $\eta^2_p$ | F | p | $\eta^2_p$ | F | p | $\eta^2_p$ |
| Visual Central: Extra Striate Cortex 1 | 8.2 | 0.000 | 0.306 | 1.9 | 0.168 | 0.026 | 12.9 | 0.001 | 0.148 | 9.5 | 0.003 | 0.113 | 2.9 | 0.094 | 0.037 |
| Visual Central: Extra Striate Cortex 2 | 8.1 | 0.000 | 0.305 | 5.8 | 0.019 | 0.072 | 14.8 | 0.000 | 0.167 | 8.7 | 0.004 | 0.105 | 5.4 | 0.023 | 0.068 |
| Visual Central: Extra Striate Cortex 3 | 8.6 | 0.000 | 0.317 | 7.0 | 0.010 | 0.087 | 10.0 | 0.002 | 0.119 | 8.3 | 0.005 | 0.101 | 3.4 | 0.068 | 0.044 |
| Visual Peripheral: Striate Cortex Calcarine 1 | 8.9 | 0.000 | 0.324 | 3.6 | 0.061 | 0.047 | 9.5 | 0.003 | 0.114 | 9.8 | 0.002 | 0.117 | 3.5 | 0.065 | 0.045 |
| Visual Peripheral: Extra Striate Inferior 1 | 11.8 | 0.000 | 0.390 | 9.9 | 0.002 | 0.118 | 11.4 | 0.001 | 0.134 | 12.9 | 0.001 | 0.149 | 8.8 | 0.004 | 0.106 |
| Visual Peripheral: Extra Striate Superior 1 | 7.9 | 0.000 | 0.299 | 0.8 | 0.366 | 0.011 | 14.3 | 0.000 | 0.162 | 8.2 | 0.005 | 0.100 | 3.1 | 0.082 | 0.040 |
| Somatomotor A: 1 | 11.9 | 0.000 | 0.391 | 3.3 | 0.071 | 0.043 | 19.4 | 0.000 | 0.208 | 11.0 | 0.001 | 0.129 | 5.8 | 0.018 | 0.073 |
| Somatomotor A: 2 | 6.1 | 0.000 | 0.247 | 2.8 | 0.101 | 0.036 | 8.1 | 0.006 | 0.099 | 7.9 | 0.006 | 0.097 | 4.0 | 0.048 | 0.052 |
| Somatomotor A: 3 | 5.2 | 0.001 | 0.220 | 0.5 | 0.481 | 0.007 | 12.3 | 0.001 | 0.143 | 7.4 | 0.008 | 0.091 | 6.0 | 0.017 | 0.075 |
| Somatomotor A: 4 | 7.4 | 0.000 | 0.286 | 2.4 | 0.129 | 0.031 | 13.5 | 0.000 | 0.154 | 8.1 | 0.006 | 0.099 | 7.5 | 0.008 | 0.092 |
| Somatomotor B: Auditory 1 | 9.3 | 0.000 | 0.334 | 4.7 | 0.034 | 0.060 | 13.9 | 0.000 | 0.158 | 11.7 | 0.001 | 0.137 | 6.0 | 0.017 | 0.075 |
| Somatomotor B: S2 1 | 8.0 | 0.000 | 0.301 | 2.2 | 0.145 | 0.028 | 7.2 | 0.009 | 0.089 | 8.5 | 0.005 | 0.103 | 3.0 | 0.085 | 0.040 |
| Somatomotor B: S2 2 | 11.3 | 0.000 | 0.379 | 15.9 | 0.000 | 0.177 | 9.7 | 0.003 | 0.116 | 8.6 | 0.004 | 0.105 | 6.4 | 0.013 | 0.080 |
| Somatomotor B: Central 1 | 15.3 | 0.000 | 0.453 | 13.3 | 0.000 | 0.152 | 9.4 | 0.003 | 0.113 | 6.8 | 0.011 | 0.084 | 2.0 | 0.160 | 0.026 |
| Dorsal Attention A: Temporal Occipital 1 | 11.7 | 0.000 | 0.388 | 1.7 | 0.199 | 0.022 | 22.0 | 0.000 | 0.230 | 11.4 | 0.001 | 0.133 | 5.3 | 0.024 | 0.067 |
| Dorsal Attention A: Parietal Occipital 1 | 10.1 | 0.000 | 0.353 | 0.2 | 0.646 | 0.003 | 17.8 | 0.000 | 0.194 | 9.1 | 0.003 | 0.110 | 3.9 | 0.052 | 0.050 |
| Dorsal Attention A: Superior Parietal Lobule 1 | 11.8 | 0.000 | 0.389 | 14.1 | 0.000 | 0.160 | 6.0 | 0.016 | 0.076 | 9.6 | 0.003 | 0.115 | 2.5 | 0.117 | 0.033 |
| Dorsal Attention B: Post Central 1 | 13.1 | 0.000 | 0.414 | 10.0 | 0.002 | 0.119 | 9.7 | 0.003 | 0.116 | 8.5 | 0.005 | 0.103 | 4.8 | 0.032 | 0.061 |
| Dorsal Attention B: Post Central 2 | 9.6 | 0.000 | 0.342 | 1.1 | 0.290 | 0.015 | 17.6 | 0.000 | 0.192 | 8.2 | 0.005 | 0.100 | 4.9 | 0.031 | 0.062 |
| Dorsal Attention B: Frontal Eye Fields 1 | 12.3 | 0.000 | 0.400 | 5.7 | 0.020 | 0.071 | 18.5 | 0.000 | 0.200 | 7.0 | 0.010 | 0.086 | 5.0 | 0.028 | 0.064 |
| Salience Ventral Attention A: Parietal Operculum 1 | 12.7 | 0.000 | 0.408 | 0.9 | 0.350 | 0.012 | 14.9 | 0.000 | 0.167 | 8.4 | 0.005 | 0.102 | 1.3 | 0.264 | 0.017 |
| Salience Ventral Attention A: Insula: 1 | 7.7 | 0.000 | 0.295 | 0.1 | 0.779 | 0.001 | 16.0 | 0.000 | 0.178 | 8.4 | 0.005 | 0.101 | 2.7 | 0.105 | 0.035 |
| Salience Ventral Attention A: Parietal Medial 1 | 8.2 | 0.000 | 0.308 | 3.4 | 0.071 | 0.043 | 15.1 | 0.000 | 0.169 | 7.9 | 0.006 | 0.096 | 3.5 | 0.064 | 0.046 |
| Salience Ventral Attention A: Frontal Medial 1 | 7.2 | 0.000 | 0.280 | 5.9 | 0.018 | 0.073 | 8.7 | 0.004 | 0.105 | 5.0 | 0.028 | 0.063 | 4.7 | 0.034 | 0.060 |
| Salience Ventral Attention B: Inferior Parietal Lobule 1 | 9.4 | 0.000 | 0.336 | 0.0 | 0.997 | 0.000 | 19.3 | 0.000 | 0.207 | 8.7 | 0.004 | 0.105 | 3.9 | 0.051 | 0.051 |
| Salience Ventral Attention B: Lateral Prefrontal Cortex | 10.1 | 0.000 | 0.353 | 3.0 | 0.089 | 0.039 | 15.3 | 0.000 | 0.171 | 8.2 | 0.005 | 0.100 | 4.5 | 0.038 | 0.057 |
| Salience Ventral Attention B: Medial Posterior Prefront | 7.8 | 0.000 | 0.298 | 0.1 | 0.728 | 0.002 | 16.2 | 0.000 | 0.180 | 9.9 | 0.002 | 0.118 | 3.4 | 0.069 | 0.044 |
| Limbic A: Temporal Pole 1 | 11.0 | 0.000 | 0.372 | 6.3 | 0.014 | 0.078 | 14.1 | 0.000 | 0.161 | 10.7 | 0.002 | 0.126 | 4.4 | 0.039 | 0.056 |
| Limbic B: Orbital Frontal Cortex 1 | 13.0 | 0.000 | 0.413 | 9.0 | 0.004 | 0.108 | 13.1 | 0.001 | 0.151 | 9.5 | 0.003 | 0.114 | 4.8 | 0.031 | 0.061 |
| Control A: Intraparietal Sulcus 1 | 11.3 | 0.000 | 0.379 | 5.1 | 0.028 | 0.064 | 18.6 | 0.000 | 0.201 | 10.4 | 0.002 | 0.123 | 5.5 | 0.022 | 0.069 |
| Control A: Lateral Prefrontal Cortex 1 | 7.8 | 0.000 | 0.297 | 1.8 | 0.186 | 0.023 | 11.0 | 0.001 | 0.129 | 10.7 | 0.002 | 0.126 | 3.0 | 0.086 | 0.039 |
| Control A: Lateral Prefrontal Cortex 2 | 7.5 | 0.000 | 0.288 | 2.1 | 0.150 | 0.028 | 12.4 | 0.001 | 0.143 | 12.2 | 0.001 | 0.141 | 5.5 | 0.021 | 0.069 |
| Control B: Temporal 1 | 10.3 | 0.000 | 0.357 | 9.2 | 0.003 | 0.111 | 7.2 | 0.009 | 0.089 | 10.3 | 0.002 | 0.122 | 5.2 | 0.026 | 0.066 |
| Control B: inferior parietal lobule 1 | 13.7 | 0.000 | 0.426 | 6.6 | 0.012 | 0.082 | 18.8 | 0.000 | 0.203 | 5.5 | 0.021 | 0.070 | 4.5 | 0.038 | 0.057 |
| Control B: Lateral Prefrontal Cortexd 1 | 6.5 | 0.000 | 0.259 | 0.6 | 0.436 | 0.008 | 12.3 | 0.001 | 0.143 | 11.0 | 0.001 | 0.129 | 4.8 | 0.032 | 0.061 |
| Control B: Lateral Prefrontal Cortexv 1 | 13.3 | 0.000 | 0.419 | 1.1 | 0.289 | 0.015 | 20.2 | 0.000 | 0.214 | 10.1 | 0.002 | 0.120 | 2.7 | 0.107 | 0.035 |
| Control C: Cingulate Posterior 1 | 15.9 | 0.000 | 0.462 | 7.0 | 0.010 | 0.087 | 16.7 | 0.000 | 0.184 | 9.5 | 0.003 | 0.114 | 4.1 | 0.046 | 0.053 |
| Control C: Precuneus 1 | 10.6 | 0.000 | 0.365 | 1.5 | 0.218 | 0.020 | 16.5 | 0.000 | 0.183 | 11.3 | 0.001 | 0.133 | 3.2 | 0.076 | 0.042 |
| Default A: Inferior Parietal Lobule 1 | 11.1 | 0.000 | 0.374 | 1.8 | 0.180 | 0.024 | 15.3 | 0.000 | 0.171 | 8.9 | 0.004 | 0.108 | 2.7 | 0.103 | 0.036 |
| Default A: Dorsal Prefrontal Cortex 1 | 6.5 | 0.000 | 0.260 | 4.0 | 0.049 | 0.051 | 9.3 | 0.003 | 0.112 | 8.7 | 0.004 | 0.105 | 3.7 | 0.059 | 0.047 |
| Default A: Precuneus Posterior Cingulate Cortex 1 | 7.5 | 0.000 | 0.288 | 5.3 | 0.024 | 0.067 | 13.8 | 0.000 | 0.157 | 8.2 | 0.005 | 0.100 | 6.8 | 0.011 | 0.084 |
| Default A: Medial Prefrontal Cortex 1 | 11.2 | 0.000 | 0.378 | 6.0 | 0.016 | 0.075 | 17.6 | 0.000 | 0.192 | 6.9 | 0.010 | 0.086 | 6.4 | 0.013 | 0.080 |
| Default B: Dorsal Prefrontal Cortex 1 | 11.6 | 0.000 | 0.385 | 5.9 | 0.018 | 0.073 | 13.4 | 0.000 | 0.153 | 12.8 | 0.001 | 0.147 | 6.8 | 0.011 | 0.084 |
| Default B: Ventral Prefrontal Cortex 1 | 7.3 | 0.000 | 0.284 | 3.2 | 0.077 | 0.042 | 11.0 | 0.001 | 0.129 | 7.5 | 0.008 | 0.092 | 5.1 | 0.026 | 0.065 |
| Default B: Ventral Prefrontal Cortex 2 | 4.1 | 0.005 | 0.182 | 3.5 | 0.067 | 0.045 | 5.0 | 0.029 | 0.063 | 7.7 | 0.007 | 0.095 | 3.9 | 0.051 | 0.051 |
| Default C: RetroSuperior Parietal Lobuleenial 1 | 2.8 | 0.032 | 0.131 | 0.0 | 0.896 | 0.000 | 4.4 | 0.040 | 0.056 | 7.6 | 0.007 | 0.093 | 2.3 | 0.133 | 0.030 |
| Default C: Parahippocampal Cortex 1 | 5.1 | 0.001 | 0.215 | 1.0 | 0.321 | 0.013 | 8.3 | 0.005 | 0.100 | 8.8 | 0.004 | 0.106 | 6.1 | 0.016 | 0.076 |
| Temporal Parietal 1 | 3.2 | 0.020 | 0.146 | 0.4 | 0.546 | 0.005 | 8.0 | 0.006 | 0.098 | 4.6 | 0.035 | 0.058 | 3.3 | 0.072 | 0.043 |
| Temporal Parietal 2 | 6.9 | 0.000 | 0.271 | 4.1 | 0.047 | 0.052 | 8.8 | 0.004 | 0.106 | 7.3 | 0.009 | 0.089 | 4.2 | 0.044 | 0.054 |
| Temporal Parietal 3 | 6.2 | 0.000 | 0.252 | 2.9 | 0.093 | 0.038 | 11.4 | 0.001 | 0.134 | 8.1 | 0.006 | 0.098 | 6.9 | 0.011 | 0.085 |
| Caudate | 4.7 | 0.002 | 0.202 | 2.1 | 0.149 | 0.028 | 5.1 | 0.027 | 0.064 | 6.5 | 0.013 | 0.080 | 2.5 | 0.115 | 0.033 |
| Putamen | 5.5 | 0.001 | 0.228 | 1.7 | 0.192 | 0.023 | 6.8 | 0.011 | 0.084 | 5.0 | 0.028 | 0.064 | 2.0 | 0.158 | 0.027 |
| Pallidum | 5.7 | 0.001 | 0.237 | 1.2 | 0.281 | 0.016 | 6.5 | 0.013 | 0.081 | 5.4 | 0.023 | 0.068 | 1.1 | 0.294 | 0.015 |
| Thalamus | 5.4 | 0.001 | 0.227 | 1.3 | 0.264 | 0.017 | 4.0 | 0.051 | 0.051 | 10.0 | 0.002 | 0.119 | 0.7 | 0.407 | 0.009 |
